# Proteomic discovery of chemical probes that perturb protein complexes in human cells

**DOI:** 10.1101/2022.12.12.520090

**Authors:** Michael R. Lazear, Jarrett R. Remsberg, Martin G. Jaeger, Katherine Rothamel, Hsuan-lin Her, Kristen E. DeMeester, Evert Njomen, Simon J. Hogg, Jahan Rahman, Landon R. Whitby, Sang Joon Won, Michael A. Schafroth, Daisuke Ogasawara, Minoru Yokoyama, Garrett L. Lindsey, Haoxin Li, Jason Germain, Sabrina Barbas, Joan Vaughan, Thomas W. Hanigan, Vincent F. Vartabedian, Christopher J. Reinhardt, Melissa M. Dix, Seong Joo Koo, Inha Heo, John R. Teijaro, Gabriel M. Simon, Brahma Ghosh, Omar Abdel-Wahab, Kay Ahn, Alan Saghatelian, Bruno Melillo, Stuart L. Schreiber, Gene W. Yeo, Benjamin F. Cravatt

**Author notes:** These authors contributed equally (Michael R. Lazear, Jarrett R. Remsberg, Martin G. Jaeger). Correspondence –.

## Abstract

Most human proteins lack chemical probes, and several large-scale and generalizable small-molecule binding assays have been introduced to address this problem. How compounds discovered in such “binding-first” assays affect protein function, nonetheless, often remains unclear. Here, we describe a “function-first” proteomic strategy that uses size exclusion chromatography (SEC) to assess the global impact of electrophilic compounds on protein complexes in human cells. Integrating the SEC data with cysteine-directed activity-based protein profiling identifies changes in protein-protein interactions that are caused by site-specific liganding events, including the stereoselective engagement of cysteines in PSME1 and SF3B1 that disrupt the PA28 proteasome regulatory complex and stabilize a dynamic state of the spliceosome, respectively. Our findings thus show how multidimensional proteomic analysis of focused libraries of electrophilic compounds can expedite the discovery of chemical probes with site-specific functional effects on protein complexes in human cells.

## Introduction

Chemical probes are vital tools for perturbing proteins and pathways in biological systems and can serve as starting points for novel therapeutics. The discovery of chemical probes has historically relied on the advent of specific functional assays for proteins of interest and devising such assays can present a major technical hurdle, especially for proteins that lack readily monitorable biochemical activities. Efforts to expand the druggable proteome have accordingly begun to introduce complementary approaches for ligand discovery that leverage binding assays with near-universal applicability to diverse types of proteins (Schreiber, 2019). Examples of technologies for the discovery of small-molecule binders of proteins include fragment-based screening (Lu et al., 2021; Scott et al., 2012), DNA-encoded libraries (DELs) (Brenner and Lerner, 1992; Gironda-Martinez et al., 2021), and chemical proteomics (Backus et al., 2016; Maurais and Weerapana, 2019; Parker et al., 2017; Spradlin et al., 2021). These methods have illuminated the broad small-molecule binding potential of proteins from structurally and functionally distinct classes. Nonetheless, whether and how small molecule-protein interactions emanating from “binding-first” assays impact the functions of proteins remain open and important questions – ones that are particularly challenging to address on a global scale when confronted with the divergent and specialized activities performed by proteins in the cell.

In considering ways to relate small-molecule binding events to functional outcomes, we looked to an emerging category of phenotypic assays that provide broad biochemical signatures of cell states. The impact of compounds on global gene and protein expression profiles of cells has, for instance, been assessed using DNA microarrays (Lamb et al., 2006; Subramanian et al., 2017) and mass spectrometry (MS)-based proteomics (Ruprecht et al., 2020), respectively. These past studies have, however, mostly evaluated compounds with known mechanisms of action, in which the gene/protein expression profiles have served to augment the understanding of established functional effects on proteins or to reveal potential off-target toxicities. When applied to naïve compounds that lack complementary protein-binding profiles, the problem of relating biochemical signatures to specific protein targets persists. We were therefore interested in determining whether the integration of two types of global profiling data – biochemical signatures and protein-binding – could facilitate the *de novo* discovery of chemical probes that produce functional effects in human cells.

Aware that small molecules can perturb other biochemical features beyond gene/protein expression, we also sought to develop complementary “function-first” signatures of compound action – specifically, global readouts of protein complexation states in cells. Many proteins function as parts of larger homo- or hetero-typic complexes (Giurgiu et al., 2019; UniProt Consortium, 2018), and small-molecule inhibitors or stabilizers of protein-protein interactions (PPIs) can serve as valuable chemical probes and therapeutic agents (Arkin et al., 2014; Jin et al., 2014; Schreiber, 2021). Several approaches have been introduced for the large-scale mapping of PPIs, including genetic (e.g., yeast-two hybrid (Fields and Song, 1989)) and biochemical (affinity purification (Huttlin et al., 2021) or co-fractionation (Kirkwood et al., 2013; Kristensen et al., 2012) coupled with MS) methods. Among these options, we viewed co-fractionation-MS as possibly most compatible with devising a streamlined and minimally biased platform for monitoring the effects of small molecules on PPIs in human cells.

Here, we present a proteomic workflow that integrates electrophilic compound effects on the complexation state of proteins with global and site-specific maps of covalent protein binding in human cells. From a modest set of electrophilic small molecules, we discover chemical probes that alter protein complexes by site-specific and stereoselective engagement of individual cysteine residues, including: i) azetidine acrylamides and butynamides that disrupt the proteasome activator complex PA28 by engaging C22 of the adaptor protein PSME1, producing global changes in MHC-I antigenic peptide presentation; and ii) tryptoline acrylamides that stabilize a transient state of the spliceosome by engaging C1111 of the splicing factor SF3B1, resulting in impairments in RNA splicing and cancer cell growth. Taken together, our findings provide a roadmap for the discovery of small-molecule modulators of PPIs that emphasizes the attributes of covalent chemistry and stereochemistry coupled with global proteomic readouts of small-molecule binding and protein complexation states in cells.

## Results

### A proteomic platform to discover small molecules that alter protein complexes in human cells

Chemical probes targeting PPIs can be challenging to discover from conventional compound libraries due to the extensive points of contact required to perturb large protein interfaces (Arkin *et al*., 2014; Smith and Gestwicki, 2012). This problem has been addressed by structure-guided approaches that facilitate the linking of weakly binding fragments into higher-affinity compounds capable of blocking PPIs (Ashkenazi et al., 2017). Covalent compounds offer an alternative and possibly more ligand-efficient strategy, where the permanent bonds formed with proteins may be sufficient to disrupt PPIs even at single points of contact (Akçay et al., 2016; Bar-Peled et al., 2017; Hacker et al., 2017; Harvey et al., 2020; Zhang et al., 2021). Previous chemical proteomic studies using cysteine-directed activity-based protein profiling (ABPP) have further demonstrated that electrophilic small molecule-sensitive, or ligandable, cysteines, are found on a wide array of proteins from different structural and functional classes (Backus *et al*., 2016; Bar-Peled *et al*., 2017; Grossman et al., 2017; Kuljanin et al., 2021; Vinogradova et al., 2020), suggesting that diverse types of protein complexes may be sensitive to covalent compound action.

To more broadly explore the potential of covalent compounds to alter PPIs, we developed a size exclusion chromatography (SEC)/MS-based proteomic method that compares the migration profiles of proteins from human cells treated with different cysteine-directed electrophilic small molecules. Similar co-fractionation-MS, or protein correlation profiling, studies using sophisticated SEC protocols have been introduced to construct large-scale *de novo* protein-protein interaction networks (Heusel et al., 2019; Kirkwood *et al*., 2013; Kristensen *et al*., 2012; Mallam et al., 2019; Skinnider and Foster, 2021). We surmised that a more simplified variant of this approach requiring fewer fractionation steps and less MS instrument time could be leveraged to identify proteins undergoing substantial shifts in SEC migration following exposure of human cells to electrophilic compounds. We specifically established a protocol to compare five SEC fractions from two cell-treatment conditions in a single tandem mass tagging (TMT (Thompson et al., 2003))-based proteomic analysis (**Figure 1A**). We found that proteins quantified from soluble proteomic lysates of the human prostate cancer cell line 22Rv1 collected from SEC experiments performed with a Superdex 200 Increase 10/300GL column spanned a wide range of molecular weights (**Figure S1A**) and showed good correlation with data from previous co-fractionation MS experiments using a much larger number of SEC fractions (40 fractions) (Kirkwood *et al*., 2013) (**Figure 1B**). Likewise, we found that the majority of protein complexes (as defined by the CORUM Core Complex database (Giurgiu *et al*., 2019)) for which two or more protein members were identified in our experiments and previous SEC-MS studies (Kirkwood *et al*., 2013) displayed similar co-elution scores (**Figure S1B** and **Dataset S1**). We additionally confirmed that the mean elution times for proteins were consistent both within replicate SEC-MS experiments performed on the same human cancer cell line and across experiments performed on distinct human cancer cell lines (**Figure S1C** and **Dataset S1**). On average, each SEC-MS experiment quantified ∼3700 total proteins (minimum of two unique quantified peptides per protein) from a starting material of 1.25 mg of soluble human cancer cell proteome. We interpret these data to indicate that a five-fraction SEC-MS protocol exhibited sufficient resolution and sensitivity to evaluate the effects of electrophilic compounds on a diverse array of protein complexes in human cells.

**Figure 1.**
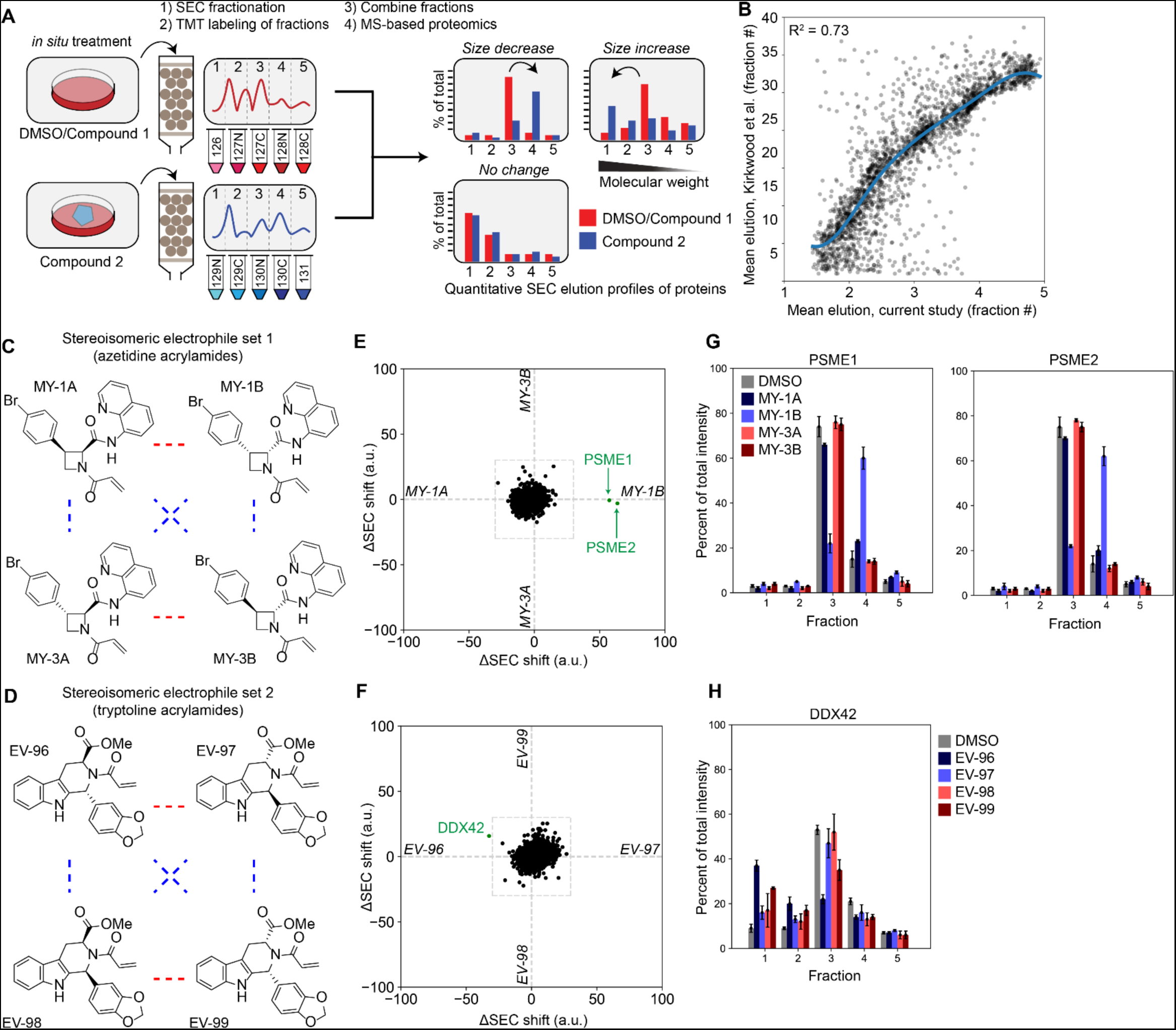
A proteomic platform to discover small-molecule modulators of protein-protein interactions in human cells. A. Schematic for screening electrophilic compounds using an SEC-MS proteomic platform. Cells are treated *in situ* with DMSO or compounds, and lysates are fractionated by size exclusion chromatography (SEC) into 5 fractions, each of which is digested with trypsin and labeled with a separate, isobaric tandem mass tag. Digests are then combined and analyzed by liquid chromatography-mass spectrometry (MS) using an Orbitrap-Fusion Tribrid MS instrument. Protein elution profiles across the five fractions are then assembled and quantified (see **Methods** section for more details). B. Comparison of the mean elution times of proteins from SEC-MS experiments performed with 22Rv1 cells under control (DMSO) conditions (x-axis) vs. SEC-MS data performed with U2OS cells from *Kirkwood* et. al (y-axis). Each dot represents a single protein detected in both experiments. Data are mean elution times (weighted average) from n = 2-11 independent experiments (current study) and n = 3 independent experiments from Kirkwood *et al*. Support vector regression line displayed in blue. C. Structures for the azetidine acrylamide set of stereoisomeric electrophilic compounds MY-1A (**1**), MY-1B (**2**), MY-3A (**3**), and MY-3B (**4**). Red lines represent enantiomers and blue lines correspond to diastereomers. D. Structures for the tryptoline acrylamide set of stereoisomeric electrophilic compounds EV-96 (**5**), EV-97 (**6**), EV-98 (**7**), and EV-99 (**8**). Red lines represent enantiomers and blue lines correspond to diastereomers. E. Protein size shift scores (arbitrary units, a.u.) plotted for SEC-MS experiments performed with proteomes from 22Rv1 cells treated with the indicated compounds (20 μM, 3 h). X-axis represents comparison of protein size shifts caused by MY-1A and MY-1B (difference in SEC shifts from DMSO vs. MY-1A and DMSO vs. MY-1B, see **Equation 4**). Y-axis represents comparison of protein size shifts caused by MY-3A and MY-3B. Data are size shift values calculated from average protein elution profiles from n = 2-11 independent experiments. F. Protein size shift scores (arbitrary units, a.u.) plotted for SEC-MS experiments performed with proteomes from 22Rv1 cells treated with the indicated compounds (20 μM, 3 h). X-axis represents comparison of protein size shifts caused by EV-98 and EV-99. Y-axis represents comparison of protein size shifts caused by MY-3A and MY-3B. Data are size shift values calculated from average protein elution profiles from n = 2-11 independent experiments. G. SEC elution profiles for PSME1 and PSME2 from 22Rv1 cells treated with azetidine acrylamides (20 µM, 3 h), revealing a stereoselective shift to lower MW fractions in MY-1B-treated cells. Data are average values ± SEM of the fractional distribution of protein reporter ion intensities from n = 2-11 independent experiments. H. SEC elution profile for DDX42 from 22Rv1 cells treated with tryptoline acrylamides (20 µM, 3 h), revealing a stereoselective shift to lower MW fractions in EV-96-treated cells. Data are average values ± SEM of the fractional distribution of protein reporter ion intensities from n = 2-11 independent experiments.

We exposed 22Rv1 cells to two structurally distinct sets of four stereoisomeric electrophilic compounds (azetidine acrylamides (**Figure 1C**) (Tao et al., 2022) and tryptoline acrylamides (**Figure 1D**) (Vinogradova *et al*., 2020) ; 20 µM, 3 h treatment), which were constructed based on principles of diversity-oriented synthesis (DOS) (Schreiber Stuart, 2000) – specifically the use of sp^3^-rich, entropically constrained, densely functionalized, and stereochemically defined scaffolds. We previously found that both sets of compounds stereoselectively engaged cysteine residues on diverse classes of proteins in human T-cells (Tao *et al*., 2022; Vinogradova *et al*., 2020). We accordingly sought herein to identify, from cells treated with stereoisomeric electrophilic compounds, proteins that showed stereoselective shifts in SEC migration (size shifts), which could then be correlated with stereoselective changes in cysteine reactivity. We quantified stereoselective size shifts, or ‘shift scores’, by calculating the Euclidean distance between individual protein elution profiles across the five SEC fractions collected from cells treated with enantiomeric compound pairs, where a stereoselective shift score of 30 or greater was considered of potential interest (see **Methods** for more details). An additional comparison to DMSO-treated cells enabled determination of which of the two enantiomeric compounds caused the observed protein size shift(s), and comparisons of the aggregate reporter ion intensity signals for quantified peptides from each protein across the five SEC fractions gave estimates of changes in protein abundance potentially caused by compound treatment. The migration profiles and abundances of most proteins were not affected in cells treated with enantiomeric pairs of electrophilic compounds (**Figure S1D, E** and **Dataset S1**). However, discrete and striking stereoselective size shifts were observed for individual proteins in cells treated with the azetidine acrylamide MY-1B (**2**) (**Figure 1E**) and the tryptoline acrylamide EV-96 (**5**) (**Figure 1F**).

MY-1B caused a size shift for two proteins – PSME1 and PSME2 – both of which moved from a higher molecular weight (MW) fraction 3 to a lower MW fraction 4 (**Figure 1G**). None of the other azetidine acrylamide stereoisomers (enantiomer MY-1A (**1**) and diastereomers MY-3A (**3**) and MY-3B (**4**)) affected the migration profiles of PSME1 and PSME2 (**Figures 1E** and **1G**). PSME1 and PSME2 form the heptameric PA28 proteasomal regulatory complex (Chen et al., 2021; Huber and Groll, 2017) (**Figure S1F**), which influences antigenic peptide processing by the immunoproteasome (Groettrup et al., 1995; Raule et al., 2014; Yamano et al., 2002). The coordinated size shifts observed for PSME1 and PSME2 suggested that MY-1B may disrupt the PA28 complex. In contrast, subunits of the 20S core proteasomal complex were unaffected by MY-1B and generally migrated, as expected, in a higher MW fraction (**Figure S1G**). The major stereoselective effect of EV-96 was an increased size shift for the RNA helicase DDX42 from a lower MW fraction 3 to a higher MW fraction 1 (**Figures 1F** and **1H**). The enantiomeric compound EV-97 (**6**) did not affect DDX42, while a more modest, but observable stereoselective shift for this protein was observed with EV-99 (**8**) compared to enantiomer EV-98 (**7**) (**Figures 1F** and **1H**). These data suggested that EV-96, and to a lesser extent EV-99, may promote the assembly of DDX42 into a larger protein complex.

### Covalent ligands stereoselectively disrupt the PA28 complex by engaging C22 of PSME1

To determine the mechanistic basis for the changes in protein migration caused by the stereoisomeric electrophilic compounds, we performed cysteine-directed ABPP experiments in 22Rv1 cells. We initially focused on the azetidine acrylamides, wherein we aimed to identify cysteines that were stereoselectively engaged by the active compound MY-1B compared to inactive stereoisomers MY-1A and MY-3A/3B. Proteomes from DMSO- or compound-treated cells (20 µM compound, 1 h) were exposed to the broad-spectrum cysteine-reactive probe iodoacetamide-desthiobiotin (IA-DTB), and probe-labeled cysteines enriched and quantified by multiplexed (TMT) MS analysis (Vinogradova *et al*., 2020). Cysteines showing a substantial loss (> 60%) in IA-DTB labeling in cells treated with one or more of the azetidine acrylamides were considered to have been engaged or liganded by those compounds. Of the ∼8,800 total quantified cysteines in 22Rv1 cells, 43 were liganded by MY-1B, and a much smaller subset (seven cysteines) were stereoselectively engaged by this compound (**Figures 2A, B**, and **S2A** and **Dataset S1**). Among the cysteines stereoselectively liganded by MY-1B was PSME1_C22 (**Figure 2A**), which resides near the PSME1-PSME2 interface (**Figure 2C**). One additional cysteine was quantified in PSME1 (C106), as well as one cysteine in PSME2 (C91), and intriguingly, these cysteines both exhibited stereoselective *increases* in IA-DTB reactivity in MY-1B-treated cells (**Figure 2D**). PSME1_C106 and PSME2_C91 are also located proximal to the protein-protein interface of PSME1 and PSME2 (**Figure 2C**), suggesting that these residues may become more solvent accessible and IA-DTB-reactive following engagement of PSME1_C22 by MY-1B, providing further evidence that this compound caused a substantial alteration in the structure of the PA28 complex. A similar profile of stereoselective cysteine reactivity changes was observed for PSME1 in another human cancer cell line (Ramos cells) treated with MY-1B (**Dataset S1**), and we further observed robust stereoselective enrichment of PSME1 by an alkyne analog of this compound (MY-11B) compared to the enantiomer (MY-11A) (**Figure 2E, F** and **Dataset S1**). Finally, we also performed cysteine-directed ABPP experiments with the tryptoline acrylamides, but we did not detect EV-96-sensitive cysteines in DDX42 (**Figure S2B** and **Dataset S1**). We will return to the mechanistic characterization of EV-96 below.

**Figure 2.**
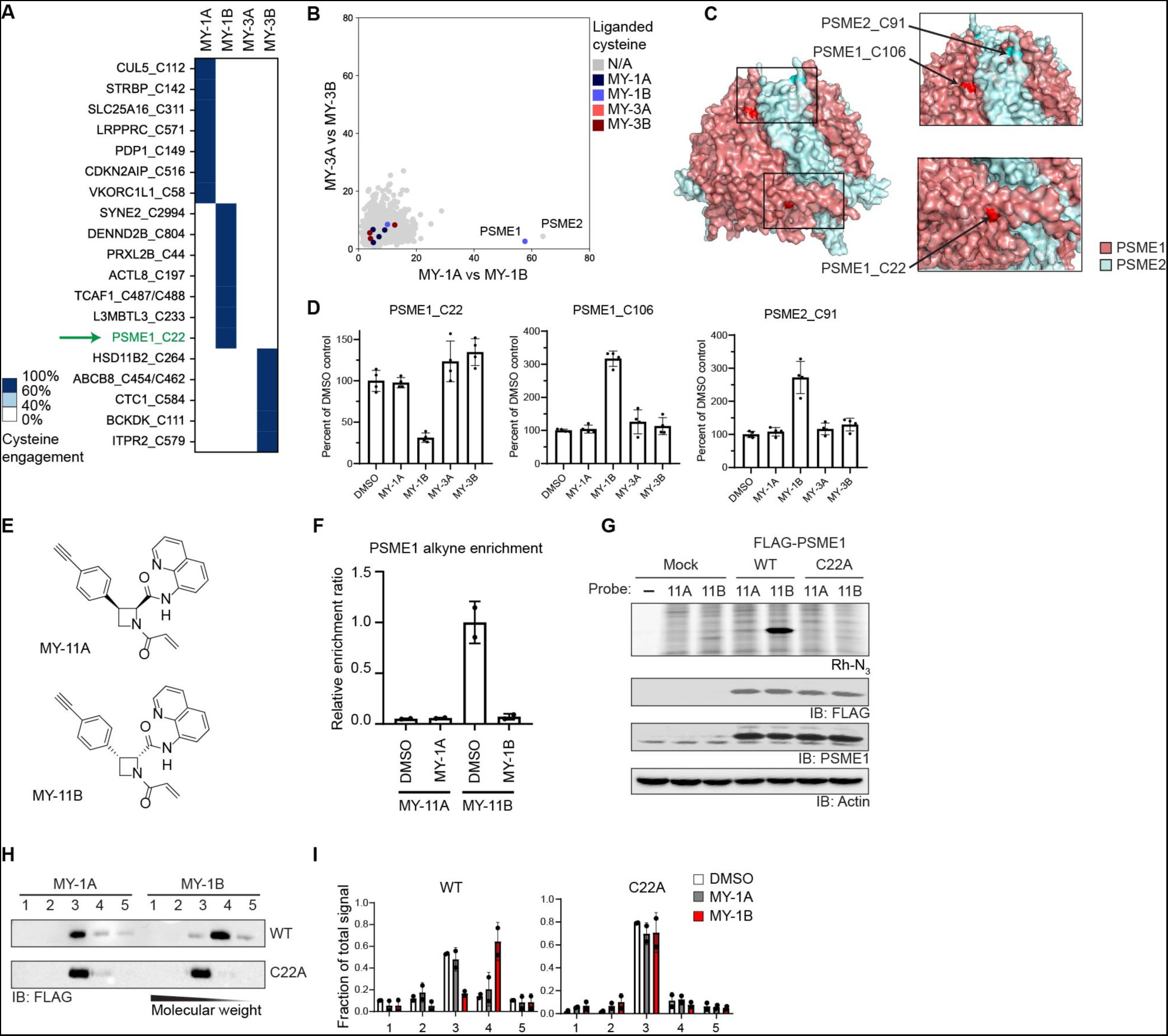
Electrophilic compounds stereoselectively disrupt the PA28 complex by engaging C22 of PSME1. A. Heatmap showing cysteines stereoselectively liganded by azetidine acrylamides in 22Rv1 cells (20 µM compound, 1 h) as determined by cysteine-directed ABPP. Cysteines were considered stereoselectively liganded if they showed > 60% reduction in IA-DTB labeling by one stereoisomeric compound and < 40% reduction for the other three stereoisomeric compounds. Data are average values from n = 4 independent experiments. See **Figure S2A** for larger heat map also showing cysteines that were liganded by more than one of the azetidine acrylamides. B. Graph showing size shifts of proteins in pairwise comparisons of SEC profiles for 22Rv1 cells treated with enantiomeric azetidine acrylamides (MY-1A vs MY-1B, x-axis; MY-3A vs MY-3B; y-axis; reanalysis of data from Figure 1E), where proteins with stereoselectively liganded cysteines are color-coded (cysteine engagement data from panel **A**). C. Crystal structure of PSME1 and PSME2 complex, highlighting the location of PSME1_C22 (bottom inset) and PSME1_C106 and PSME2_C91 (top inset) (PDB: 7DRW). D. Reactivity profiles for cysteines in PSME1 and PSME2 from 22Rv1 cells treated with azetidine acrylamides, highlighting the decrease in IA-DTB labeling of PMSE1_C22 and the increase in labeling of PSME1_C106 and C91 in MY-1B-treated cells compared to cells treated with other azetidine acrylamide stereoisomers. Data are extracted from experiment described in **A**. E. Structures of alkynylated azetidine acrylamide probes MY-11A (inactive) (**9**) and MY-11B (active) (**10**). F. Quantification of stereoselective enrichment and competition of PSME1 by MY-11B (5 µM, 1 h) compared to MY-11A (5 µM, 1 h) and competition by the respective competitor MY-1A and MY-1B (20 µM, 2 h pretreatment) in Ramos cells. Data are average values ± SD normalized to MY-11B treatment group, n = 2 independent experiments. G. MY-11B, but not MY-11A, labels recombinantly expressed WT-PSME1, but not the C22A-PSME1 mutant in a site- and stereo-specific manner. Proteomic lysates from HEK293T cells expressing FLAG-epitope-tagged WT and C22A-PSME1 (or Mock HEK293T cells) were treated with alkyne probe (2.5 μM, 30 min) followed by copper-catalyzed azide alkyne cycloaddition (CuAAC) to an azide-rhodamine tag using and analysis by SDS-PAGE and in-gel fluorescence scanning (top image) or western blotting (bottom three images). Results are from a single experiment representative of two independent experiments. H. MY-1B, but not MY-1A, causes a size shift in WT-, but not C22A-PMSE1. Western blot analysis of SEC profiles for WT and C22A-PSME1 recombinantly expressed as FLAG-epitope tagged proteins in 22Rv1 TetR cells (tetracycline-inducible system; 22Rv1 cells stably expressing Tet repressor and transiently transfected with a plasmid containing PSME1 cDNA under control of a Tet operator/CMV promoter). Cells were treated with MY-1A or MY-1B (10 µM, 3 h) prior to analysis by SEC. Results are from a single experiment representative of two independent experiments. I. Quantification of data shown in panel **H**. Data are average values ± SD from n = 2 independent experiments.

We confirmed by gel-ABPP experiments that MY-11B stereoselectively labeled WT-PSME1, but not a C22A-PSME1 mutant (**Figure 2G**), recombinantly expressed as FLAG epitope-tagged proteins in 22Rv1 cells (**Figure S2C**). Recombinant PSME1 has been shown to form homooligomeric structures in the absence of PSME2 (Huber and Groll, 2017), thus furnishing a convenient assay to further explore the effects of electrophilic compounds on PPIs involving PSME1. Recombinant WT-PSME1 exhibited similar SEC elution profiles to endogenous PSME1 with predominant migration in fraction 3 (**Figure S2D**). Treatment of cells with MY-1B, but not MY-1A, caused a clear size shift for recombinant WT-PMSE1 to fraction 4, while the C22A-PSME1 mutant was unaffected (**Figures 2H, I**). These data thus support that MY-1B disrupts PSME1-mediated PPIs by specifically engaging C22.

Other studies of covalent ligands have shown that substituting an acrylamide with a less reactive butynamide electrophile can improve selectivity for certain protein targets like the B-cell kinase BTK (Barf et al., 2017). We found here that the butynamide analogue of MY-1B – MY-45B (**Figure 3A**) – showed even greater potency for engaging PSME1_C22 (**Figure 3B**), displaying an IC_50_ value of ∼0.4 µM with recombinant WT-PSME1 (**Figure 3C**). We confirmed by cysteine-directed ABPP that MY-45B, but not the enantiomeric control compound MY-45A (**Figure 3A**), engaged C22 of endogenous PSME1 in human and mouse cells with improved potency and selectivity compared to MY-1B (**Figure S2E, F** and **Dataset S1**). Across > 10,000 quantified cysteines in human 22Rv1 cells, MY-45B (5 µM, 3 h) substantially engaged seven cysteines, and only two of these cysteines stereoselectively reacted with MY-45B compared to MY-45A – C22 of PSMSE1 and C258 of the helicase DDX49 (**Figure 3D, E**), a protein that has not been implicated in proteasome regulation. MY-45B, but not MY-45A, caused the expected size shifts for endogenous PSME1 and PSME2 (**Figures 3F** **and S2G**), as well as recombinant WT-PSME1 (**Figure S2H**). The overall properties of MY-45B, including good potency and proteome-wide selectivity, as well as pairing with an inactive enantiomer, led us to designate this compound as a suitable chemical probe for studying the function of PSME1 and the PA28 complex in cells.

**Figure 3.**
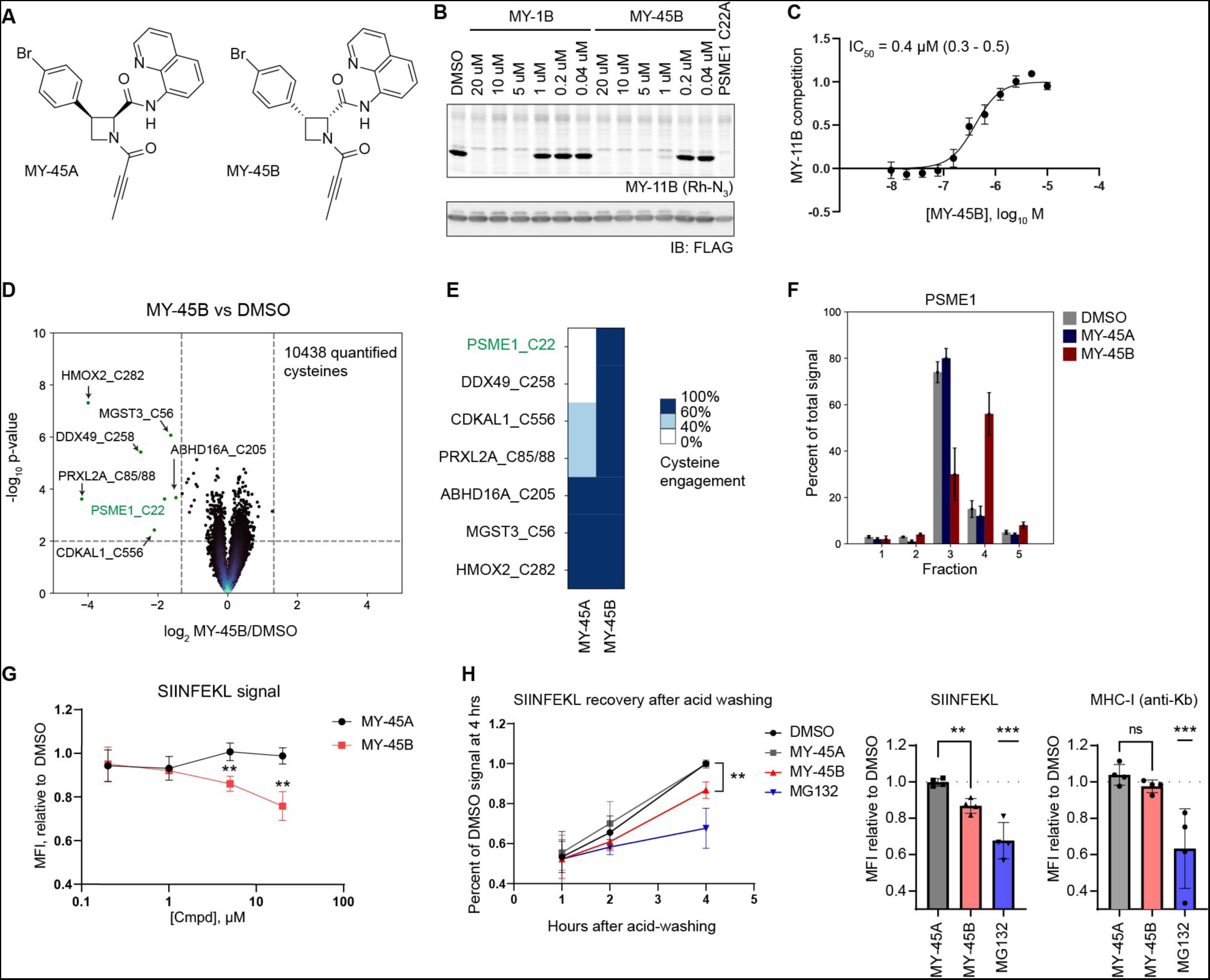
An azetidine butynamide engaging PSME1_C22 functionally impair MHC-I antigenic peptide presentation. A. Structures for azetidine butynamides MY-45A (inactive) (**11**), and MY-45B (active) (**12**). B. Comparison of potency of engagement of PSME1 by MY-1B and MY-45B. Proteomic lysates from HEK293T cells expressing FLAG-epitope tagged WT- or C22A-PSME1 were treated with the indicated concentrations of compound (2 h) followed by MY-11B (2.5 µM, 30 min), CuAAC with rhodamine-azide, and analysis by SDS-PAGE and in-gel fluorescence scanning. Results are from a single experiment representative of two independent experiments. C. Concentration-dependent engagement of PMSE1 by MY-45B as measured by blockade of MY-11B probe labeling (as shown in panel **B**). Experiments performed as described in panel (B). Data are average values ± SEM from n = 6 independent experiments, IC_50_ and 95% CI (confidence intervals) are listed. D. Volcano plot showing cysteines substantially (> 60% reduction in IA-DTB labeling) and significantly (*p* value < 0.01) liganded by MY-45B (5 µM, 3 h) in 22Rv1 cells as determined by cysteine-directed ABPP. Data are average values from n = 4 independent experiments. E. Heatmap displaying MY-45B-liganded cysteines (from panel **D**) and their reactivity with enantiomer MY-45A. F. SEC-MS elution profile for endogenous PSME1 in 22Rv1 cells treated with DMSO, MY-45A, or MY-45B (20 μM, 3 h). Data are average values ± SEM of the fractional distribution of reporter ion intensities from n = 2-11 independent experiments. G. Functional effects of MY-45B treatment on antigen processing. Mouse T lymphoma cells constitutively expressing chicken ovalbumin (E.G7-Ova) were treated with DMSO or compound (5 µM for MY-45A and MY-45B; 10 µM for MG132) prior to analysis. Left panel, SIINFEKL median fluorescence intensity (MFI) for the indicated compound treatment groups relative to DMSO at 1, 2, or 4 h post-acid-wash. Right panel, MFI for the indicated compound treatment groups relative to DMSO 4 h after acid wash for both SIINFEKL and overall MHC-I for 5 µM MY-45A, 5 µM MY-45B, and 10 µM MG132. Data are average values ± SD relative to DMSO from n = 4 independent experiments. **, p < 0.01, *** p < 0.001 compared to MY-45A treatment. H. Concentration-dependent impairment of antigen processing by MY-45B, 4 h post-acid wash. Data are average values ± SD relative to DMSO from n = 3 independent experiments. **, p < 0.01 compared to MY-45A treatment.

The PA28 complex has been found to impact MHC class I (MHC-I) antigen presentation through influencing the proteasomal processing of a select, but not exhaustively determined, set of antigens (Keller et al., 2015; Yamano *et al*., 2002). The chicken ovalbumin peptide SIINFEKL is commonly used as a model peptide in antigen presentation studies and is among the antigens regulated by the PA28 complex (Yamano *et al*., 2002). Using a kinetic assay that measures the rate of recovery of MHC-I peptide presentation following acid washing (Shunji et al., 1987; Sturm et al., 2021) (**Figures S3A and S3B**), we found that mouse T lymphoma cells constitutively expressing chicken ovalbumin (E.G7-Ova) showed a time- and concentration-dependent reduction in SIINFEKL presentation – but not overall MHC-I surface expression – following treatment with MY-45B, but not MY-45A (**Figures 3G** and **3H**), and these data correlated with stereoselective engagement of mouse PSME1_C22 by MY-45B as determined by cysteine-directed ABPP (**Figure S3C and Dataset S1**). In contrast, the direct proteasome inhibitor MG132 suppressed both SIINFEKL peptide and MHC-I presentation (**Figure 3H**).

These data, taken together, indicated that azetidine acrylamides and butynamides stereoselectively disrupt the structure and function of the PA28 complex by engaging PSME1_C22. We next asked how these covalent PSME1 ligands more globally impact MHC-I peptide presentation.

### PSME1 disruption modulates MHC-I-peptide interactions in human leukemia cells

We first generated cell models genetically disrupted for PSME1 or PSME2 using CRISPR/Cas9 methods in the near-haploid human leukemia cell line KBM7 (**Figure 4A**). Notably, genetic disruption of either PSME1 or PSME2 led to loss of both proteins in KBM7 cells (**Figure 4A**), suggesting that the stability of each protein depends on its preferred interaction partner. MS-based proteomic analysis of sgControl versus sgPSME1 cells, as well as cells treated with MY-45B or MY-45A or DMSO, revealed few changes in the abundance of proteins (**Figure S3D**), indicating that genetic or chemical disruption of the PA28 complex did not perturb the general protein-degrading function of the proteasome. In this proteomic experiment, MY-45B-treated sgControl cells, but not MY-45A-treated sgControl cells, displayed a dramatic decrease (> 90%) in signals for the tryptic peptide containing PSME1_C22 (R.EDLCTK), while other PSME1-derived tryptic peptides were unaffected (**Figure S3E**), thus providing clear evidence for robust site-specific and stereoselective engagement of PSME1_C22 by MY-45B.

**Figure 4.**
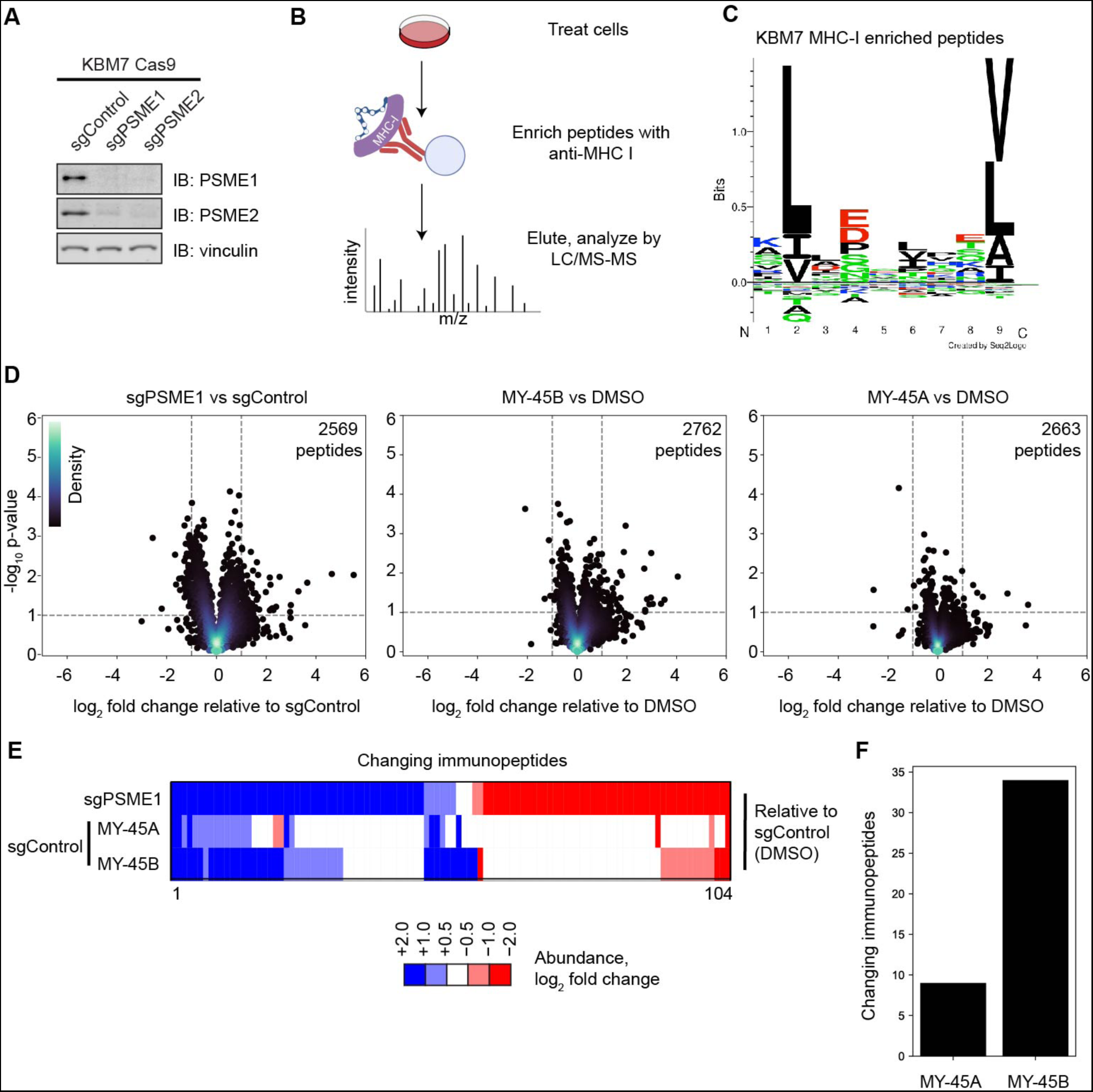
Chemical or genetic perturbation of PSME1 modulates MHC-I-immunopeptide interactions in human leukemia cells. A. Western blots showing PSME1 and PSME2 expression in KBM7 CRISPR/Cas9 control (sgAAVS1), PSME1 (sgPSME1), or PSME2 (sgPSME2) cell lines. Vinculin was used as a loading control. Results are from a single experiment representative of two independent experiments. B. Cartoon schematic for anti-MHC class I immunopeptidomics protocol. After treatment with compounds, cells are lysed, and MHC-I bound peptides immunoprecipitated with anti-MHC-I antibody, eluted, and analyzed by LC-MS. C. Motif analysis of peptides enriched by MHC-I co-immunoprecipitation from KBM7 cells. D. Volcano plots showing substantially (> two-fold increase or decrease) and significantly (*p* value < 0.05) changing MHC-I bound immunopeptides in sgPSME1 vs sgControl (sgAAVS1) cells (left), DMSO- vs MY-45B-treated sgControl cells (middle), or DMSO- vs MY-45A-treated sgControl cells (right) (10 μM compound, 8 h). Data are average values from n= 3-4 independent experiments. E. Heatmap of MHC-I-bound immunopeptides that are substantially and significantly changing in at least one comparison group (sgPSME1 vs sgControl; MY-45A- vs DMSO-treated sgControl; or MY-45B- vs DMSO-treated sgControl) as defined in panel **D**. F. Bar graph showing the number of MHC-I-bound immunopeptides that are substantially and significantly changing in MY-45A- vs DMSO-treated sgControl or MY-45B- vs DMSO-treated sgControl cells as defined in panel **D**.

We next evaluated the antigenic peptide profiles of sgPSME1 cells compared to sgControl cells, as well as sgControl cells treated with MY-45B (10 µM, 8 h) compared to MY-45A- or DMSO-treated cells, following an established protocol for the affinity enrichment, identification, and quantification of MHC-I-bound peptides by LC-MS (Purcell et al., 2019) (**Figure 4B**). Supporting the technical quality of the protocol, motif analysis of the eluted immunopeptides revealed a consensus motif for the KBM7 HLA haplotype HLA-A*02:01 (Duncan et al., 2012) (**Figure 4C**). sgPSME1 cells exhibited substantial (log_2_ fold change > 1) and significant (*p* < 0.05) changes in ∼100 of > 2,500 quantified immunopeptides, which reflected both decreases and increases in co-enrichment with MHC-I (**Figure 4D, E**). A smaller subset of MHC-I-associated peptides were altered by MY-45B (34 peptides) in sgControl cells (**Figure 4D-F**), and these peptides showed a striking (∼70%) overlap with the peptides altered in sgPSME1 cells (**Figure 4D-F**). In contrast, the inactive enantiomer MY-45A caused far fewer changes in the antigenic peptide profile of KBM7 cells (< 10 peptide changes; **Figure 4D-F**). These findings, taken together, indicate that the acute, pharmacological disruption of the PA28 by covalent ligands engaging C22 of PSME1 produces discrete changes in MHC-I-associated peptides that are largely consistent with those caused by chronic genetic loss of PSME1/PSME2. We speculate that the antigenic peptide changes observed in sgPSME1 cells, but not MY-45B-treated cells, may be caused by longer-term perturbations of the PA28 complex. While too few antigenic peptides were altered by genetic or pharmacological disruption of PSME1 in KBM7 cells to assess potential sequence motif preferences associated with these changes, it is possible that such features may be revealed in future studies that apply PSME1 ligands to a greater number of cell types or under immunostimulatory conditions.

### Tryptoline acrylamides stereoselectively alter spliceosome composition and function

Returning to our finding that EV-96 promotes a stereoselective shift of DDX42 to a higher molecular weight form despite not apparently engaging any cysteines in this protein (**Figure 1F, H**), we gathered additional clues pertaining to the mechanism of action of EV-96 by quantitative proteomics, which revealed a striking, stereoselective reduction in several proteins within 8 h post-treatment in 22Rv1 cells (**Figure 5A**). None of the affected proteins appeared to possess EV-96-sensitive cysteines (**Dataset S1**), which argued against a direct, ligand-induced degradation model. Gene Ontology (GO)-term analysis revealed that the EV-96-sensitive proteins were enriched in cell division and cell cycle functions (**Figure 5B**), suggesting that these protein changes may be an indirect consequence of EV-96 effects on cancer cell proliferation. Consistent with this hypothesis, we found that EV-96 caused a stereoselective blockade in the growth of 22Rv1 cells (**Figure 5C**). EV-96 also impaired the growth of other human cancer cells lines, such as Ramos and THP1 cells (**Figure S4A**). Interestingly, the EV-96-induced protein changes showed only modest overlap with a “frequent responder” group of proteins previously shown to represent common changes caused by diverse cytotoxic compounds (Ruprecht *et al*., 2020) (**Figure S4B**), suggesting a discrete mechanism of anti-proliferative action for EV-96.

**Figure 5.**
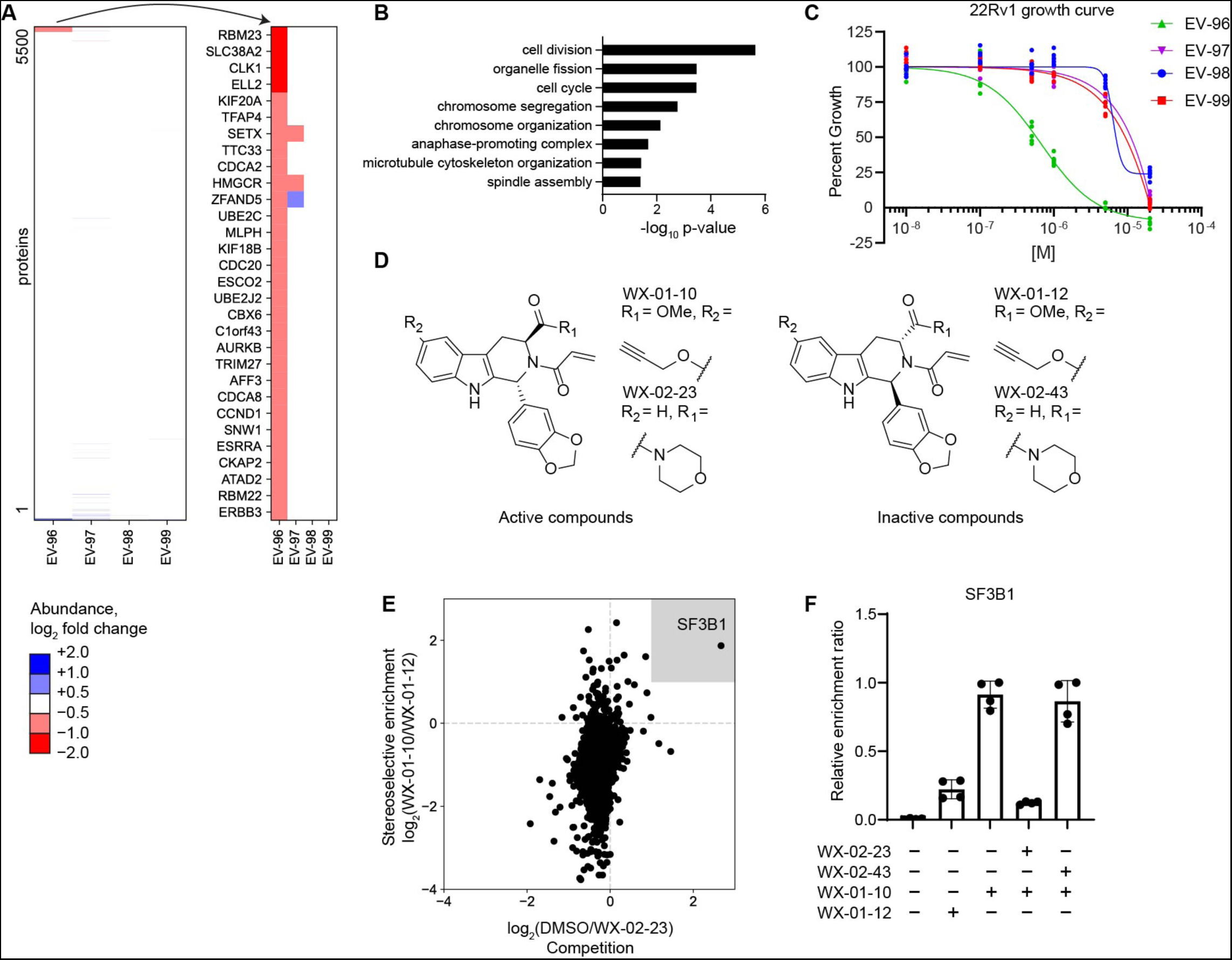
Tryptoline acrylamides that stereoselectively engage SF3B1 alter protein abundances and block the proliferation of cancer cells. A. EV-96 treatment stereoselectively alters protein abundances in 22Rv1 cells. Left panel, Heatmap of protein abundance changes in 22Rv1 cells treated with tryptoline acrylamides (20µM, 8 h). Right panel, Blow up of heatmap showing proteins with > 33% decreases in abundance in 22Rv1 cells treated with EV-96. Data are average values from n = 4-6 independent experiments. B. Gene ontology enrichment for proteins stereoselectively decreased in abundance by EV-96 (panel **A**) reveals an enrichment for cell cycle-related proteins. C. EV-96 stereoselectively inhibits 22Rv1 cell growth. Cells were treated with the indicated concentrations of tryptoline acrylamides for 72 h, after which cell growth was measured by CellTiter-Glo. Data are relative to DMSO control from n = 6 independent experiments. D. Structures of alkyne (WX-01-10 and WX-01-12) and morpholino amide (WX-02-23 and WX-02-43) analogues of EV-96 and EV-97. E. Chemical proteomic identification of SF3B1 as a protein that is stereoselectively enriched by WX-01-10 and stereoselectively competed in enrichment by WX-02-23. X-axis: log_2_ competition ratio values for proteins enriched by alkyne probe WX-01-10 (10 µM, 1 h) in 22Rv1 cells pretreated with DMSO or WX-02-23 (5 µM, 2 h pretreatment) as a competitor. Y-axis: log_2_ enrichment ratio values for proteins treated with active alkyne probe WX-01-10 vs inactive probe WX-01-12 (10 µM, 1 h). Data are average values from n = 4 independent experiments (See also **Figure S5F** for schematic of this experiment). F. Quantification of stereoselective enrichment and competition of SF3B1 by active alkyne probe (WX-01-10) and competitor (WX-02-23) vs inactive enantiomer alkyne probe (WX-01-12) and inactive enantiomer competitor (WX-02-43). Data are average values ± SD normalized to WX-01-10 treatment group for n = 4 independent experiments.

One possible explanation for the inability to identify an EV-96-sensitive cysteine relevant to the observed anti-proliferative activity would be that this compound produces its proteomic and cell growth effects through a non-covalent mechanism. Arguing against this hypothesis, however, we found that a non-electrophilic propanamide analogue of EV-96 did not inhibit cancer cell growth (**Figures S4C and S4D**). Recognizing instead that some electrophilic compound-sensitive cysteines may evade detection by cysteine-directed ABPP if, for instance, they reside on non-proteotypic peptides (i.e., tryptic peptides that are infrequently detected by MS-based proteomics due to size and/or poor physicochemical properties (Mallick et al., 2007)), we adopted a complementary chemical proteomic strategy to identify stereoselectively engaged protein targets of EV-96 using alkyne-modified analogues of this compound. First, we generated alkyne-modified analogues of EV-96 and its inactive enantiomer EV-97 – WX-01-10 and WX-01-12, respectively (**Figure 5D**) – and confirmed that these compounds maintained differential cell growth inhibitory effects (**Figure S4E**). We concurrently found, through screening a larger set of tryptoline acrylamides, a morpholine amide analogue of EV-96 – WX-02-23 – that exhibited ∼4-fold greater antiproliferative activity (IC_50_ of 170 nM (150-180nM)) and maintained stereoselectivity in comparison to the enantiomeric compound WX-02-43 (**Figures 5D** **and S4F, G**). WX-02-23 also induced apoptosis in a subset of the cancer cell lines examined (**Figure S4H**). With this set of chemical tools, we pursued protein targets by MS-based proteomic experiments involving the pre-treatment of cancer cells with WX-02-23 or its inactive enantiomer WX-02-43, followed by treatment with active alkyne WX-01-10 or inactive alkyne enantiomer WX-01-12, where a protein target responsible for the observed anti-proliferative effect should display: i) stereoselective enrichment by WX-01-10 (in comparison to WX-01-12); and ii) competition in its enrichment by WX-02-23, but not WX-02-43 (**Figure S4I**). Only a single protein – the spliceosome factor SF3B1 – was found to meet these criteria (**Figures 5E, F** and **Dataset S1**). We also found that SF3B1 showed a similar stereoselective competition profile in cells pretreated with EV-96 versus EV-97 before alkyne WX-01-10 enrichment (**Figure S4J** and **Dataset S1**).

SF3B1 is an ∼150 kDa member of the spliceosome and is essential for stabilizing the branch point adenosine prior to intron removal (Larsen, 2021). Natural product-based modulators of SF3B1 have been described, and these compounds show broad spectrum anti-proliferative activity (Lee and Abdel-Wahab, 2016). Consistent with SF3B1 being a direct, stereoselective target of tryptoline acrylamides, we found in gel-ABPP experiments that WX-01-10, but not its enantiomer WX-01-12 labeled an ∼150 kDa protein in 22Rv1 cells, and this labeling was blocked by WX-02-23, but not its enantiomer WX-02-43 (**Figures 6A** **and S5A**). Interestingly, WX-01-10 labeling of the 150 kDa protein was also blocked by pre-treatment with pladienolide B (**Figure 6A** and **S5A**), a natural product modulator of SF3B1 (Kotake et al., 2007). WX-02-23 and pladienolide B both induced expression of p27 (**Figure 6A**), a previously described feature of spliceosome modulators reflecting an aberrant splicing event that removes a C-terminal ubiquitination site from p27 and prevents cells from progressing through the G2/M checkpoint (Kaida et al., 2007).

**Figure 6.**
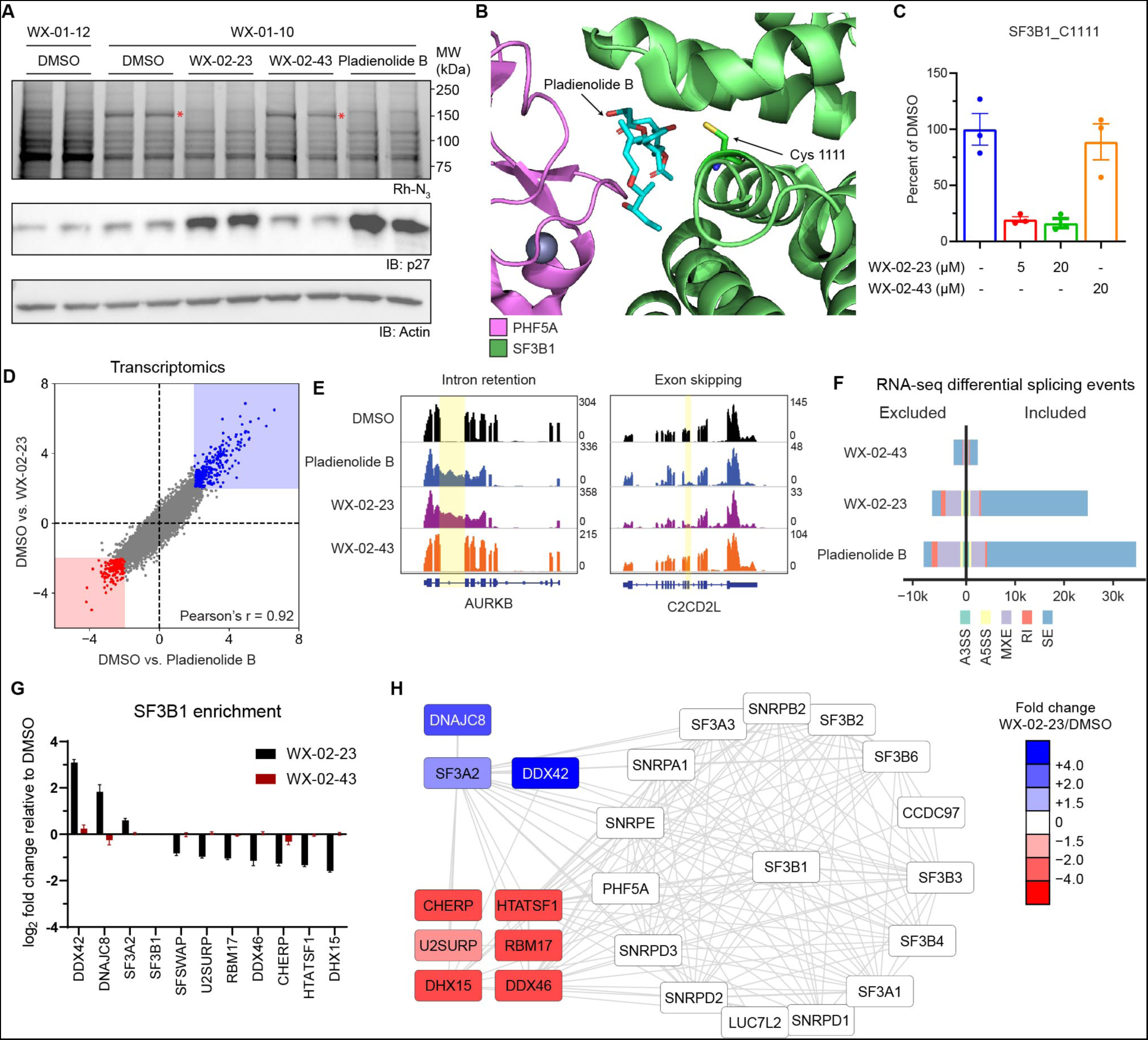
Tryptoline acrylamides engage C1111 of SF3B1 and stereoselectively modulate spliceosome structure and function. A. Stereoselective labeling of a 150 kDa protein in 22Rv1 cells as determined by gel-ABPP. Top panel, Gel-ABPP showing proteins labeled by alkyne probes in 22Rv1 cells. Cells were pre-treated with DMSO, WX-02-23 (1 µM), WX-02-43 (1 µM) or pladienolide B (10 nM) for 24 h followed by treatment with WX-01-10 or WX-01-12 (1 µM, 1 h), lysis, CuAAC with rhodamine azide, SDS-PAGE, and in-gel fluorescence scanning. Red asterisk marks a 150 kDa protein that is stereoselectively labeled by WX-01-10 (compared to cells treated with inactive enantiomer alkyne probe WX-01-12) and blocked in labeling by WX-02-23 and pladienolide B, but not inactive enantiomer WX-02-43. This profile is consistent with MW and compound interaction profile of SF3B1. Middle panel, western blot demonstrating increased p27 expression in WX-02-23 and pladienolide B-treated cells. Bottom panel, western blot of actin as a loading control. Results are from a single experiment representative of two independent experiments. B. Crystal structure of SF3B1-PHF5A complex bound to pladienolide B, highlighting the location of C1111 (PDB: 6EN4). C. Quantification of stereoselective engagement of SF3B1_C1111 by WX-02-23 as measured by targeted cysteine reactivity profiling experiments performed in 22Rv1 cells 5 μM or 20 μM compound, 3 h). Data are average values ± SD relative to DMSO from n = 2-3 independent experiments. D. Scatter plot of mRNA transcript abundance changes in 22Rv1 cells treated with WX-02-23 (5 µM), pladienolide B (10 nM), or DMSO for 8 h. RNA-seq data are average values shown as log_2_ fold change relative to DMSO for n = 3 independent experiments. E. Examples of intron retention in Aurora Kinase B and exon skipping in C2CD2L caused by pladienolide B and WX-02-23 in 22Rv1 cells. F. Summary of alternative splicing events caused by pladienolide B (10 nM), WX-02-23 (5 µM), and inactive enantiomer WX-02-43 (5 µM) in 22Rv1 cells (8 h) compared to DMSO treatment, as identified with rMATS by threshold of |PSI| > 0.1 and FDR < 0.05. Data represent values from three independent experiments. G. WX-02-23 remodels SF3B1 protein-protein interactions. Quantification of proteins showing differential co-immunoprecipitation with SF3B1 (> 1.5 fold change increase or decrease) in HEK293T cells treated with WX-02-23, WX-02-43, or DMSO (5 μM, 3 h). Co-immunoprecipitated was performed with anti-SF3B1 antibody (CST #14434). Data are average log_2_ fold changes ± SD relative to DMSO from n = 4-7 independent experiments. H. Interactome map from STRING database filtered for proteins identified as SF3B1 interactors in the co-immunoprecipitation described in panel **I**. Red and blue proteins indicate interactions that are weakened or strengthened by WX-02-23 treatment. Data are average log_2_ fold changes relative to DMSO from n = 4-7 independent experiments.

The co-crystal structure of a pladienolide B-SF3B1 complex has confirmed that this interaction is reversible involving contacts with both SF3B1 and PHF5A (Cretu et al., 2018). Integrating this information with our data on the tryptoline acrylamides indicating a covalent mechanism of action and a shared binding pocket with pladienolide B, we surmised that this pocket might contain a WX-02-23-sensitive cysteine. Review of the SF3B1-pladienolide B structure identified a single candidate cysteine C1111 in the binding pocket (**Figure 6B**). SF3B1_C1111 has rarely been quantified in our cysteine reactivity profiling datasets (current and past (Backus *et al*., 2016; Bar-Peled *et al*., 2017; Vinogradova *et al*., 2020)), suggesting that either this cysteine is poorly reactive with IA probes or that it resides on a non-proteotypic tryptic peptide that is difficult to detect by MS-based proteomics. Consistent with the latter hypothesis, we found that the IA-DTB adduct with a tryptic peptide containing C1111, which is 28 amino acids in length with 14 hydrophobic residues (A, F, I, L, M, V), eluted at the tail end of our standard liquid chromatography gradients (140 min, ∼38% acetonitrile) (**Figure S5B**). We established a targeted proteomic assay using parallel reaction monitoring to more consistently quantify the IA-DTB-C1111 tryptic peptide adduct along with several other IA-DTB-labeled cysteines in SF3B1, which revealed that WX-02-23 stereoselectively blocked IA-DTB reaction with C1111 (**Figures 6C** **and S5C**), but did not affect other cysteines in SF3B1 (**Figure S5D**). The tryptic peptide containing C1111 also has another cysteine C1123; however, this cysteine is distal to the pladienolide B-binding pocket (**Figure S5E**), and chemical proteomic experiments performed with trypsin-GluC double digests revealed that the peptide from amino acids 1122-1137 of SF3B1 was unaffected in IA-DTB reactivity in WX-02-23-treated cells (**Figure S5F** and **Dataset S1**), indicating that C1123 is not directly engaged by WX-02-23. Finally, our cysteine-directed ABPP data also confirmed that EV-96 and WX-02-23 did not engage PHF5A_C26 (**Dataset S1**), which was recently found to be targeted by spliceostatin A, another natural product modulator of the spliceosome (Cretu et al., 2021; Kaida *et al*., 2007).

WX-02-23 and pladienolide B caused strikingly similar changes to the transcriptomes and proteomes of cancer cells as measured by RNA-seq (**Figure 6D**) and MS-based proteomics (**Figure S6A**), respectively. These changes were not observed with the inactive enantiomer WX-02-43 (**Figure S6B**). Both compounds also showed greater anti-proliferative effects in Panc 05.04 cells expressing the cancer-associated K700E-SF3B1 mutant compared to Panc 04.03 cells expressing WT-SF3B1 (Foy and McMullin, 2019), although the differential impact was modest in magnitude (**Figure S6C**), which is consistent with previous findings with other pladienolide B-related SF3B1 ligands (Seiler et al., 2018). A deeper analysis of the RNA-seq data revealed that WX-02-23 and pladienolide B also altered mRNA splicing in similar ways, including the induction of both exon skipping and – to a lesser extent – intron retention events (**Figures 6E, F** and **S6D**). No such splicing effects were observed with WX-02-43 (**Figure 6E, F** and **S6D**). Finally, we found that 22Rv1 cells stably overexpressing Y36C-PHF5A, a point mutant that has been found to confer resistance to the anti-proliferative activity of pladienolide B (Teng et al., 2017), also protected against the growth inhibitory effects of WX-02-23 (**Figure S6E**). SF3B1 reactivity with the alkyne probe WX-01-10 was additionally disrupted in 22Rv1 cells expressing Y36C-PHF5A (**Figure S6F**), consistent with this mutation impairing binding of covalent tryptoline acrylamides to the spliceosome. Modeling studies also supported the importance of PHF5A_Y36 in promoting interactions with WX-02-23 (**Figure S6G**).

We interpret these data, taken together, as strong evidence that WX-02-23 and related tryptoline acrylamides produce their anti-proliferative effects through covalent modification of SF3B1, which in turn perturbs spliceosome function in a manner that resembles the effects of structurally unrelated natural product modulators of the spliceosome such as pladienolide B.

### Covalent SF3B1 ligands stabilize a spliceosome state with enhanced binding to DDX42

Curious to understand whether and how covalent modification of SF3B1 might relate to the SEC migration change in DDX42 caused by tryptoline acrylamides (**Figure 1H**), which was also observed for the more potent SF3B1 ligand WX-02-23 (**Figure S7A**), we noted some literature precedence for DDX42 (or SF3b125) physically associating with the spliceosome (Will et al., 2002). Interestingly, we found that fraction 1, to which DDX42 shifted in EV-96 or WX-02-23-treated cells, also contained other spliceosome components, including SF3B1 (**Figure S7A**), suggesting that SF3B1 ligands may promote DDX42 binding to the spliceosome. Consistent with this hypothesis, immunoprecipitation (IP)-MS proteomics revealed that DDX42 associated with SF3B1 to a much greater extent in WX-02-23-treated cancer cells compared to DMSO- or WX-02-43-treated cancer cells (**Figure 6G** and **Dataset S1**). WX-02-23 also caused a broader remodeling of SF3B1 interactions, including enhanced association with splicing factor DNAJC8 and decreased interactions with other spliceosome components (**Figures 6G, H**).

DDX42 contributions to the spliceosome remain poorly understood, and the limited functional studies performed on this helicase to date have mainly focused on spliceosome-independent activities (Bonaventure et al., 2022; Uhlmann-Schiffler et al., 2006). To investigate DDX42’s role in splicing, we introduced an N-terminal degradation tag (dTAG) into the endogenous DDX42 locus of HCT-116 cells (Erb et al., 2017). The resulting homozygous clonal cell line expressed a DDX42-dTAG fusion protein that could be degraded to near-completion within 1 h of treatment with the Von-Hippel-Lindau (VHL)-recruiting heterobifunctional small molecule dTAG^v^-1 (Nabet et al., 2020) (**Figure 7A**). Longer-term treatment with dTAG^v^-1 (72 h) caused noticeable growth impairment, but did not appear to be cytotoxic (**Figure 7B**). This chemical genetic system provided a way to study the mechanistic interactions of DDX42, SF3B1 modulatory compounds, and the broader spliceosome on a time scale that is comparable to fast-acting cellular processes such as splicing (Jaeger and Winter, 2021).

**Figure 7.**
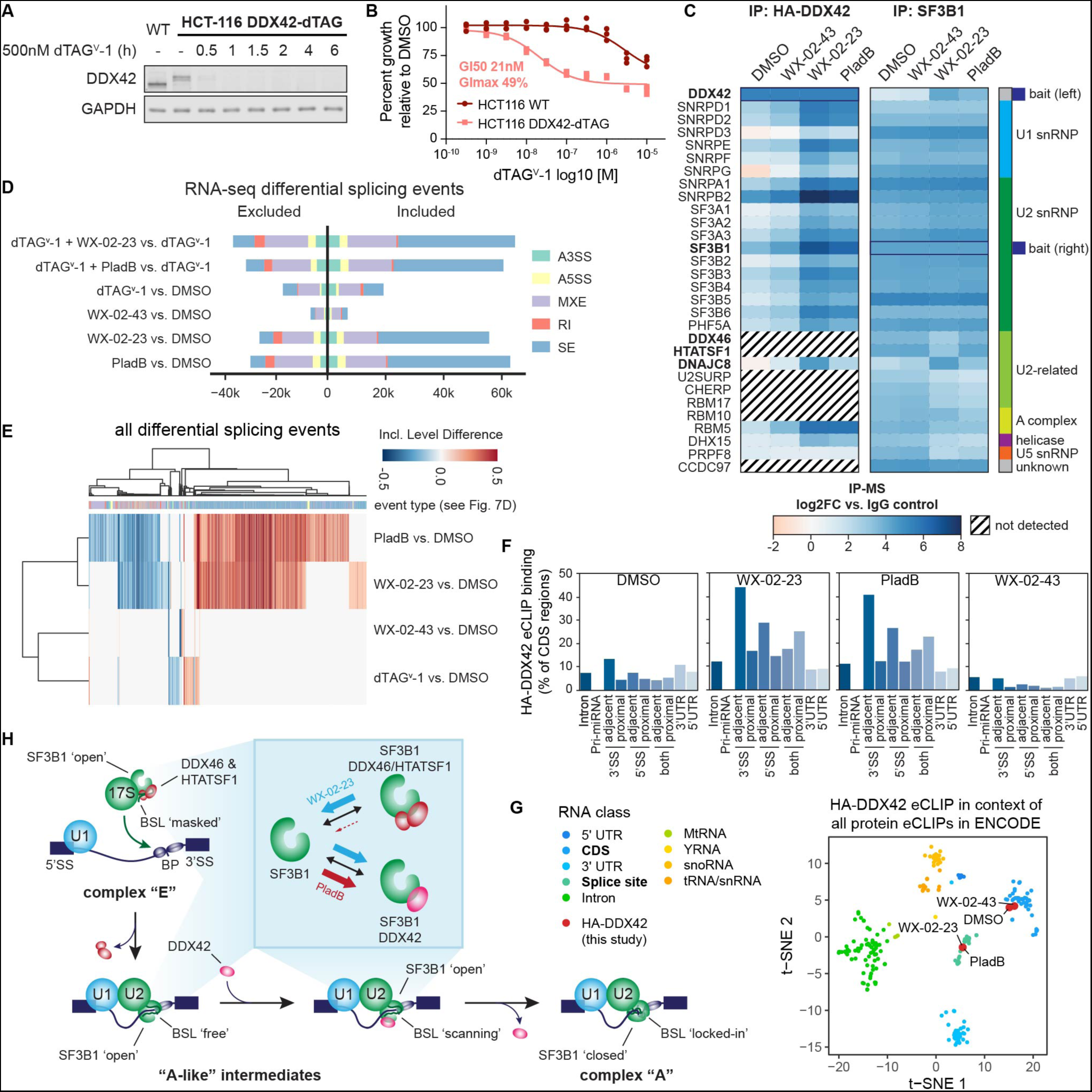
DDX42 facilitates spliceosome branch point selection. A. Time-resolved western blot analysis of dTAG^v^-1 ligand-induced DDX42 degradation in endogenously engineered DDX42-dTAG HCT-116 cells. B. Acute removal of DDX42 leads to inhibition of cell proliferation. DDX42-dTAG or wild-type HCT-116 cells were treated with the indicated concentrations of dTAG^v^-1 for 72 h, after which cell growth was measured by CellTiter-Glo. Data are relative to DMSO control from n = 3 independent experiments. C. Heatmap showing proteins that enriched in HA-DDX42 (left) or SF3B1 (right) immunoprecipitation-mass-spectrometry (IP-MS) experiments in DDX42-dTAG HCT-116 cells treated with WX-02-23 (5 µM), WX-02-43 (5 µM), pladienolide B (PladB, 10 nM), or DMSO for 3 h. Proteins were required to be detected in at least two independent experiments and included if they were either enriched in DMSO vs. IgG (log2FC >1 for HA, >2 for SF3B1) or in WX-02-23 vs. DMSO (log2FC >2). Results are average values from 2-4 independent experiments. The resulting list of 41 high-confidence interactors was input into the StringDB protein-protein interaction database and the largest connected component of 30 proteins form the basis of the heatmap. Data were normalized to corresponding bait and are shown as log2 fold-enrichment vs. IgG control. Proteins were annotated based on their proposed function in spliceosome sub-complexes. Diagonal black lines indicate proteins that were not detected in HA-DDX42 IP-MS experiments. Bold type mark proteins either directly liganded by WX-02-23 and pladienolide B (SF3B1) or substantially affected in interactions with SF3B1 by WX-02-23 and/or pladienolide B. D. Alternative splicing events triggered by indicated combinations of DDX42 degradation +/- treatment with pladienolide B (PladB, 10 nM), WX-02-23 (5 µM) or WX-02-43 (5 µM) in DDX42-dTAG HCT-116 cells. Cells were pre-treated for 1 h with either 500 nM dTAGv-1 or DMSO, followed by addition of either DMSO or compounds to the same supernatant for 8 h. Splicing changes were identified with rMATS threshold of |PSI| > 0.1 and FDR < 0.05. Data represent values from three independent experiments. E. Clustered heatmap of inclusion level differences between indicated compound treatments and DMSO control for alternative splicing events from panel **D**. Columns are annotated by the type of alternative splicing event in the color scheme of panel **D**. Data represent values from three independent experiments. F. DDX42 RNA-binding profiles in DMSO-vs compound-treated DDX42-dTAG HCT-116 cells measured by eCLIP-seq using the HA-tag of the dTAG fusion. DDX42-dTAG HCT-116 cells were treated with WX-02-23 (5 µM), WX-02-43 (5 µM), or Pladienolide B (100 nM) for 3 h. Data are average values from two independent experiments. eCLIP enriched windows (FDR<0.2) were called and annotated using the Skipper analysis pipeline (Boyle *et al*., 2022) and are depicted as percent binding relative to coding sequence (CDS). Proximal denotes within 500 bases and adjacent denotes within 100 bases from the annotated splice site (SS). G. tSNE of HA-DDX42 eCLIP samples in the context of all available eCLIP datasets that were generated by the ENCODE consortium. DDX42-dTAG HCT-116 cells were treated with WX-02-23 (5 µM), WX-02-43 (5 µM), or Pladienolide B (100 nM) for 3 h. Data are average values from n = 2 independent experiments. H. Proposed model depicting the role of DDX42 in facilitating the branch point selection step of spliceosome function. Inset summarizes differential SF3B1 complexation effects caused by synthetic, covalent (WX-02-23) vs natural product, reversible (pladienolide B) SF3B1 ligands.

One model to explain the increased association of DDX42 with SF3B1 in WX-02-23-treated cells would be that tryptoline acrylamides selectively reacted with SF3B1 when this protein was bound to DDX42. However, our dTAG^v^-1 system revealed that degradation of DDX42 did not substantially alter the stereoselective reactivity of alkyne probe WX-01-10 with SF3B1 (**Figure S7B, C**). Additionally, cells lacking DDX42 still displayed robust anti-proliferative responses to WX-02-23 and Pladienolide B (**Figure S7D**). These data thus suggested an alternative model wherein ligand binding to SF3B1 stabilized a spliceosome assembly state that displays enhanced affinity for DDX42. Consistent with this model, IP-MS studies using the HA-tag of the DDX42-dTAG fusion construct revealed that DDX42 principally interacted with components of the U1, U2, and SF3b spliceosome subcomplexes, and these interactions were strongly and stereoselectively enhanced by WX-02-23 treatment (**Figure 7C** and **Dataset S1**). These IP-MS data thus indicated that DDX42 measurably, but transiently binds an A-like spliceosome complex (Wahl et al., 2009), and this interaction is profoundly stabilized by SF3B1 ligands.

### DDX42 facilitates spliceosome branch point selection

We next assessed the impact of acute DDX42 degradation on splicing. RNA-seq analysis of cells post-degradation of DDX42 (dTAG^v^-1 treatment, 9 h) revealed substantial splicing changes, including exon skipping and – to a lesser degree – intron retention events (**Figure 7D**). These splicing alterations were less dramatic than those caused by the SF3B1 ligands WX-02-23 and pladienolide B, and these ligands also maintained their major splicing effects in cells lacking DDX42 (**Figure 7D**). These data are consistent with our chemical proteomic results indicating that DDX42 is not required for the engagement of SF3B1 by covalent ligands (**Figure S7B, C**). Interestingly, the splicing changes promoted by DDX42 loss qualitatively differed from those triggered by SF3B1 ligands (**Figure 7E**). We specifically observed that DDX42 loss increased the usage of stronger branch point sequences and decreased the usage of weaker branch points at differentially included exons (**Figure S7E**), suggesting that DDX42 may facilitate branch point selection. To test this hypothesis more directly, we evaluated DDX42-mRNA interactions by eCLIP-seq (enhanced crosslinking and immunoprecipitation followed by sequencing) (Van Nostrand et al., 2016) using the hemagluttinin (HA) epitope tag component of the dTAG fusion. These eCLIP-seq experiments revealed that, in untreated cells, DDX42 mainly bound to the coding region of (pre-)mRNAs, but, following exposure of cells to SF3B1 ligands WX-02-23 or pladienolide B, the DDX42-RNA interaction preferences strongly shifted towards regions within or near splice sites (**Figure 7F and G** and **Dataset S2**) (Boyle et al., 2022; Van Nostrand et al., 2020). Our results, taken together, thus suggest that DDX42 dynamically binds SF3B1 as part of an A-like spliceosome complex that is in close contact with (pre-)mRNAs and splice sites to facilitate branch point selection. The distinct splicing effects caused by DDX42 degradation versus SF3B1 ligands may in turn reflect differences in the functional outcomes associated with physical loss of DDX42 versus ligand-induced trapping of the spliceosome in specific subcomplexes that include a state showing enhanced interactions with DDX42.

As part of the 17S U2 snRNP and before binding pre-mRNA, SF3B1 is found in an ‘open’ conformation with the U2 snRNA branchpoint-interacting stem-loop (BSL) masked by two proteins, HTATSF1 and the RNA helicase DDX46 (Cretu *et al*., 2021; Zhang et al., 2020). Upon binding pre-mRNA and interacting with other spliceosome components, DDX46, through its ATPase activity, releases HTATSF1 and itself, presumably to allow scanning of the BSL for pre-mRNA branchpoint sequences. Upon recognition of a branchpoint sequence, SF3B1 normally switches to a closed conformation to stabilize the newly formed BSL-branchpoint helix (Haselbach et al., 2018). Structural work has revealed that pladienolide B impairs this conformational switch to the closed state by acting as a wedge in an SF3B1 hinge region, leading to reduced fidelity of branchpoint selection (Cretu *et al*., 2018). Intriguingly, DDX46 and HTATSF1 were not among the proteins that co-immunoprecipitated with DDX42 (**Figure 7C** and **Dataset S1**), and both DDX46 and HTATSF1 displayed reduced binding to SF3B1 in WX-02-23-treated cells (**Figure 6G, H**, **Figure 7C**, and **Dataset S1**). It thus appears that HTATSF1/DDX46 and DDX42 binding to SF3B1 might be mutually exclusive. Consistent with this conclusion, a recently deposited cryo-EM structure of a reconstituted DDX42-SF3b complex has revealed that the binding interface for SF3B1-DDX42 directly overlaps with that of SF3B1-HTATSF1/DDX46 (**Figure S7F**) (Zhang, 2022a; b; Zhang *et al*., 2020). Finally, we found that the WX-02-23-dependent disruption of DDX46 and HTATSF1 interactions with SF3B1 was preserved in cells lacking DDX42 (**Figure S7H**), providing further evidence that covalent SF3B1 ligands bind to an open state of the spliceosome lifecycle that shows enhanced affinity for DDX42 and reduced interactions with DDX46 and HTATSF1. Considering the prevailing model that DDX46 activity and HTATSF1 release prime the scanning of the U2 BSL for branch point sequences (Cretu *et al*., 2021; Zhang *et al*., 2020), we suggest that DDX42 subsequently facilitates the stable selection of branch points as part of the SF3B1 open complex (**Figure 7H**).

Before concluding, we call attention to a curious and as-of-yet unexplained finding that WX-02-23 and pladienolide B, despite binding to the same SF3B1-PHF5A pocket, appear to differentially reshape SF3B1 interactions. While both compounds promoted SF3B1 binding to DDX42, only WX-02-23 impaired interactions with DDX46 and HTATSF1 (**Figure 7C** and **Dataset S1**). Even experiments performed with 10X higher concentrations of pladienolide B (100 nM) that fully suppress cancer cell growth, and including the compound in the lysis and immunoprecipitation buffers, did not result in changes in DDX46/HTATSF1 binding to SF3B1 (**Figure S7H**). Considering further that a structure has been reported of pladienolide B bound to an SF3B1-DDX46/HTATSF1 complex (Zhang, 2022b), we conclude that this natural product, unlike WX-02-23, does not perturb SF3B1 interactions with DDX46 and HTATSF1 (**Figure 7H**). Moreover, while WX-02-23 and pladienolide B generally caused similar splicing changes, some differences were also observed (**Figure 7E**). While we do not yet understand how such compound-specific effects on spliceosome composition and function may distinctly impact cellular biochemistry and physiology, we believe that the covalent SF3B1 ligands reported herein offer a mechanistically distinct and synthetically accessible class of compounds for further structural and functional exploration of this important topic.

## Discussion

The pursuit of chemical probes stands to benefit from the introduction of generalized assays that can be applied to diverse types of proteins, preferably, in physiologically relevant settings. Chemical proteomics has emerged as an attractive strategy to discover small-molecule binders for proteins in native biological systems (Backus, 2019; Cuesta and Taunton, 2019; Jones, 2019; Spradlin *et al*., 2021). Original ABPP strategies using active site-directed chemical probes produced data where small-molecule binding to a protein could be inferred to cause a functional effect (i.e., a compound discovered to bind the active site of a kinase or hydrolase could be assumed to inhibit this enzyme (Greenbaum et al., 2000; Liu et al., 1999; Patricelli et al., 2007)). As the concepts of ABPP have been extended to enable assessment of small-molecule binding far beyond enzyme active sites to include virtually any protein in the human proteome (Backus *et al*., 2016; Bar-Peled *et al*., 2017; Vinogradova *et al*., 2020), fundamental challenges emerge in relating binding events to functional outcomes. Establishing this connection with assays that assess the functional features of many proteins in parallel could greatly accelerate chemical probe discovery versus a more traditional one-at-a-time investigation of individual small molecule-protein interactions. Here, we have introduced a “function-first” proteomic platform aimed at translating a broad swath of small molecule-protein binding events into a subset of interactions that perturb the complexation state of proteins in cells. Through doing so, we discovered chemical probes that alter the structure and function of key protein complexes involved in proteasomal peptide processing and mRNA splicing in human cells.

To our knowledge, the azetidine acrylamides and butynamides described herein that engage C22 of PSME1 represent the first chemical probes targeting the PA28 proteasome regulatory complex. Consistent with the proposed role of the PA28 complex in regulating MHC class I antigenic peptide processing (Raule *et al*., 2014; Yamano *et al*., 2002), we found that covalent PSME1 ligands altered a discrete set of MHC-I immunopeptides. Most of these immunopeptide changes were also observed in cells genetically disrupted for PSME1. Future studies where cells are treated for longer periods of time with PSME1 ligands may clarify whether the additional immunopeptide changes found in sgPSME1 cells, but not MY-45B-treated cells are due to chronic versus acute disruption of the PA28 complex or reflect more fundamental differences between genetic versus pharmacological mechanisms of targeting this complex. Additional areas of interest would include evaluating the impact of PMSE1 disruption on the MHC-I antigenic peptide repertoire of disease-relevant systems, such as cancer cells that have developed resistance to proteasome inhibitors (Dytfeld et al., 2016).

The tryptoline acrylamides that engage C1111 of SF3B1 are distinct from other chemical probes described for this protein, which are reversible natural product-derived compounds of greater structural complexity. The simplified and synthetically accessible tryptoline acrylamides may thus facilitate a distinct and more extensive exploration of the structure-activity relationships for SF3B1 modulators. Our initial studies already suggest that WX-02-23 and pladienolide B affect spliceosome complexes in different ways, as both compounds stabilized SF3B1-DDX42 interactions while WX-02-23, but not pladienolide B, disrupted SF3B1 association with DDX46 and HTATSF1. How such differences in spliceosome remodeling may translate into functional effects on splicing is an important topic for future investigation, as the evaluation of analogs of pladienolide B has indicated that individual SF3B1 ligands can differentially modulate splicing outcomes (Seiler *et al*., 2018).. Considering further that pladienolide B analogs are in clinical development for cancers, including those with high-frequency mutations in spliceosome components like SF3B1 (Brunner and Steensma, 2020; DeNicola and Tang, 2019), it would be of interest to investigate whether the covalent tryptoline acrylamide ligands can be optimized to preferentially impact the functions of common cancer-related SF3B1 mutants (e.g., K700E-SF3B1). As covalent ligands, the tryptoline acrylamides also provided versatile target engagement probes for SF3B1, which facilitated key mechanistic insights, including the discovery that these compounds maintain engagement in cells lacking DDX42, while losing SF3B1 engagement in resistant cells expressing a Y36C-PHF5A mutant. Finally, our studies have also contributed to a more fundamental understanding of the function of DDX42 as a dynamic component of the spliceosome, where this protein appears to specifically contribute to branch site selection. The transient nature of DDX42 binding to the spliceosome may explain why a recent structure of the DDX42-associated SF3b complex was solved by reconstitution of recombinantly expressed proteins (Zhang, 2022a), and we are hopeful that the stabilization of endogenous DDX42 binding to the spliceosome by covalent SF3B1 ligands may offer a path to additional structures that include a greater proportion of spliceosome components, including additional dynamic subunits that are stabilized in binding to SF3B1 in the presence of WX-02-23 (e.g., DNAJC8; (**Figure 6G, H**, **Figure 7C, Figure S7H,** and **Dataset S1**).

Reflecting on why our approach was successful at identifying selective, cell-active chemical probes that perturb PPIs from a small number of test compounds (two sets of four stereoisomeric acrylamides), we posit a few important factors. First, covalent ligands, by leveraging a combination of reactivity and recognition features, may provide an advantage for chemically targeting historically challenging protein classes such as adaptor proteins like PSME1. Second, we believe that the DOS principles of constructing focused compound libraries bearing sp^3^-rich cores that are stereochemically defined, entropically constrained, and densely functionalized may further improve the probability of engaging PPI sites, at least when compared to more fragment-like and/or sp^2^-based small-molecule libraries. An additional advantage of DOS-constructed libraries is that they can furnish chemical probes paired with physicochemically matched, inactive enantiomeric control compounds for cell biological studies (Feldman et al., 2022; Kim et al., 2004; Vinogradova *et al*., 2020). Finally, the function-first screening approach likely also facilitated the discovery of chemical probes by prioritizing follow-up studies on a subset of covalent liganding events that alter protein complexation states in cells.

Our studies also highlight some of the challenges and potential limitations of combining SEC-MS with cysteine-directed ABPP to relate functional effects on protein complexes to covalent liganding events in human cells. Some of the proteins showing electrophilic compound-induced size shifts by SEC, for instance, did not possess obvious cysteines engaged by these compounds. There are multiple potential reasons for this apparent disconnect. First, as we found for PSME2 and DDX42, electrophilic compounds may cause size shifts for proteins within a complex in addition to, or even other than, the direct target of the compounds. Fortunately, proteomic efforts to globally map protein complexes in human cells (Huttlin *et al*., 2021) have provided a rich set of data to draw upon when relating covalent ligandability maps to protein size shifts measured by SEC, such that compound effects on both direct targets and indirect associated partners in a complex may be interpretable. We also note the potential for cysteine reactivity profiling itself to illuminate such relationships, as exemplified by the stereoselective increase in reactivity of a specific cysteine in PSME2 in cells treated with chemical probes targeting PSME1 (**Figure 2D**). An electrophilic compound-induced size shift may also prove difficult to connect to a specific liganding event using cysteine-directed ABPP if the relevant cysteine is on a non-proteotypic peptide (as we initially found for C1111 of SF3B1) or is engaged by the compound with low stoichiometry. Either of these challenges should be addressable by using alkyne variants of electrophilic compounds, which enable direct enrichment and analysis of covalent protein targets (as we demonstrated for chemical probes targeting SF3B1). It is also important to acknowledge that some protein size shifts caused by electrophilic compounds may be overlooked by our current SEC protocol. For instance, detecting proteins that shift between two large complexes may require alternative SEC columns that better separates proteins in the very high MW range, and the analysis of membrane protein complexes would require adding a detergent solubilization step to the protocol. Finally, many protein complexes are dynamic or low affinity and may therefore disassemble following cell lysis. To enhance the discovery of compound-induced changes in very large, membrane-associated, or dynamic protein complexes, alternative approaches for mapping protein-protein interactions, such as *in situ* chemical crosslinking (Kleiner et al., 2018; Larance et al., 2016; Wheat et al., 2021), cellular thermal shift profiling (Tan et al., 2018), or proximity labeling (Cho et al., 2020; Huttlin *et al*., 2021), may prove useful.

Projecting forward, we are intrigued by the prospects of profiling larger electrophilic compound libraries with function-first proteomic assays. The SEC-MS method, while not high-throughput, should prove capable of assaying up to 30 compounds per week with a single FPLC/LC-MS system. And, if one elected to perform targeted (rather than untargeted) proteomic analyses (Uzozie and Aebersold, 2018) focused on specific protein complexes of interest, the throughput of the method could likely be further improved. Our findings also point to the potential importance of the design features, rather than sheer size, of compound libraries when seeking PPI modulators, as the DOS-inspired compound sets examined herein furnished chemical probes that modulate protein complexes with well-defined structure-activity relationships (e.g., stereoselectivity and site-specificity). In the future, it would be interesting to better understand the respective contributions of reversible binding affinity (*K_i_*) and cysteine reactivity (*k_inact_*) to the stereoselective interactions observed with DOS-inspired electrophilic compounds. Additionally, many PPI interfaces may lack cysteine residues, and it is therefore provocative to consider whether incorporating alternative electrophiles targeting other nucleophilic amino acids (Abbasov et al., 2021; Brulet et al., 2020; Cuesta and Taunton, 2019) into DOS designs, or replacing the electrophile with a photoreactive group to capture reversible binding interactions (Parker *et al*., 2017), might further enhance the chemical proteomic discovery of compounds that modulate protein complexes in cells.

We finally call attention to the numerous electrophilic compound-cysteine interactions discovered herein that did not apparently affect protein complexation state (e.g., **Figures 2A, B**, **3E**, and **S2A,** and **Dataset S1**). These liganded cysteines were found on structurally and functional diverse protein types, including enzymes, transporters, transcriptional regulators, RNA-binding proteins, and adaptor proteins. While it may be tempting to interpret that such liganding events occur at “silent” sites on proteins (i.e., sites for which compound binding does not produce a functional effect on the protein), we are reluctant to draw such a conclusion at this stage of analysis. First, we only assessed one of many possible functional features of proteins in cells (protein-protein interactions) through a single analytical method (SEC-MS) that may overlook shifts in very large or membrane-associated protein complexes as discussed earlier. Some apparently silent sites may thus manifest functionality when analyzed by the complementary methods to globally profile protein-protein interactions noted above (Kleiner *et al*., 2018; Larance *et al*., 2016; Wheat *et al*., 2021) or that assess other types of biomolecular interactions (e.g., protein-DNA interactions (Ruprecht et al., 2022)), as well as the turnover, post-translational modification state (Huang et al., 2019; Navarrete-Perea et al., 2018) and localization (Huh et al., 2003; Thul et al., 2017) of proteins. These findings, along with the continued discovery of allosteric inhibitors for diverse proteins (Chen et al., 2016; Nussinov and Tsai, 2015; Ostrem et al., 2013) and the structural mapping of hotspots for small-molecule binding at protein interfaces (Wells and McClendon, 2007), suggest that many, perhaps even most, ligandable sites on proteins have functional potential. Fully unlocking this functionality appears poised to gain from integrating structurally diversified electrophilic chemistry with generalized assays that record the myriad ways compounds modulate protein activities in human cells.

## AUTHOR CONTRIBUTIONS

Conceptualization: M.R.L., J.R.R, and B.F.C; Data curation: M.R.L., J.R.R., M.G.J., G.M.S., B.F.C; Formal analysis: M.R.L., J.R.R., M.G.J., K.R., H.H, S.J.H., J.R., H.L., B.F.C; Investigation: M.R.L., J.R.R., M.G.J., K.E.D., E.N., L.R.W., G.L.L., J.G., S.B., J.V., T.W.H., V.F.V., C.J.R., M.M.D.; Methodology: M.R.L., J.R.R., M.G.J., B.F.C; Resources: S.J.W., M.A.S., D.O., M.Y., J.V., S.J.K, I.H., J.R.T., G.M.S., B.G., O.A.-W., K.A., A.S., B.M., S.L.S., G.W.Y.; Software: M.R.L.; Supervision: G.W.Y., B.F.C.; Visualization: M.R.L., J.R.R., M.G.J., K.R., H.H, S.J.H., J.R.; Writing – original draft: M.R.L. and J.R.R.; Writing – review & editing: M.R.L., J.R.R., M.G.J., G.W.Y., B.F.C.

## STAR METHODS

### RESOURCE AVAILABILITY

#### Lead contact

Further information and requests for resource and reagents should be directed to and will be fulfilled by the lead contact, Benjamin F. Cravatt

#### Materials availability

All chemical probes and other elaborated electrophilic compounds generated in this study are available from the Lead Contact with a completed Materials Transfer Agreement.

#### Data and code availability

The mass spectrometry proteomics data have been deposited to the ProteomeXchange Consortium via the PRIDE (Perez-Riverol et al., 2019) partner repository with the dataset identifier PXD029655. Raw RNA-sequencing data has been reposited to NCBI under GEO: GSE185373, GSE220185. All other data that support the findings of this study are available from the corresponding authors upon reasonable request. Processed proteomic and RNA-sequencing data are provided in Dataset S1 and S2. This paper does not report original code.

Any additional information required to reanalyze the data reported in this paper is available from the lead contact upon request.

### EXPERIMENTAL MODEL AND SUBJECT DETAILS

#### Cell lines

22Rv1 (ATCC, CRL-2505), MCF7 (ATCC, HTB-22), Ramos (ATCC, CRL-1596), THP1 (ATCC, TIB-202), HEK293T (ATCC, CRL-3216), KBM7 (Georg Winter, CeMM, Vienna; RRID:CVCL_A426), HCT-116 (ATCC, CCL-247), Panc 04.03 (ATCC, CRL-2555), Panc 05.04 (ATCC, CRL-2557), and E.G7-Ova (ATCC, CRL-2113) were grown in in RPMI (22Rv1, Ramos, THP1, Panc 04.03, Panc 05.04, E.G7-Ova), DMEM (HEK293T, HCT-116), IMDM (KBM7), or EMEM (MCF7), supplemented with 10% fetal bovine serum (FBS), 2 mM L-alanyl-L-glutamine (GlutaMAX), penicillin (100 U/ml), and streptomycin (100 μg/ml) and maintained at 37 °C with 5% CO_2_.

*-Generation of stably transduced clonal 22Rv1 cells expressing TetR:* HEK293FT cells (Thermo R70007, supplemented with MEM nonessential amino acids, 25-025-Cl Corning) (5x10^6^) were plated in 10-cm plates and allowed to attach overnight. To 1mL of serum-free DMEM the following were added: 3 µg of pLenti3.3/TR, 2.25 μg psPAX2.0 plasmid (Addgene, catalog number: 12260) and 0.75 μg CMV VSV-G (Addgene, catalog number: 98286) and 36 μL lipofectamine 2000 transfection reagent (Invitrogen). Reagents were flicked to mix, and after 15 min incubation, the transfection mixture was added dropwise to plates containing cells. Medium was exchanged approximately 16 h post transfection, virus-containing supernatants were then collected 48 h post transfection and filtered, and then used to infect 22Rv1 cells in 6-well plates in the presence of 6 μg/ml Polybrene (Santa Cruz). The media was replaced 24 h later, cells were allowed to recover for an additional 24 h, and then geneticin (400 μg/mL) was added for selection. Clonal pLenti3.3/TR expressing cells were selected by cloning cylinder and subsequently transfected with constructs of interest.

*-Transfection of epitope tagged and mutant proteins of interest:* Clonal pLenti3.3/TR expressing 22Rv1 cells were seeded into 6cm dishes and 24 h later transfected with proteins of interest cloned into pLenti6.3. 1 μg of pLenti6.3 plasmid with gene of interest and 3 μL of PEI (1 μg/μL) were added to 200 μL serum-free RPMI and incubated for 15 minutes. Medium of cells was replaced to contain tetracycline (0.1-1 μg/mL final concentration) and then transfection mixture was added dropwise. Cells were assayed 24-48 h later.

*-Generation of DDX42-dTAG knock-in HCT-116 cells:* Knock-in of a dTAG cassette into the endogenous DDX42 locus was performed as previously described (Jaeger et al., 2020). A pX330A_sgPITCh_sgDDX42_N sgRNA/Cas9 plasmid was cloned via standard oligo-annealing and ligation methods using two sense/antisense DNA oligos (sgDDX42_N_sense 5’-CACCGattcctaacaggtcagtcat; sgDDX42_N_anti 5’-AAACatgactgacctgttaggaatC). A pCRIS-PITChv2 repair template plasmid was cloned using pCRIS-PITChv2-Puro-dTAG (BRD4) (Addgene #91793) as a PCR template to introduce 20-22 bp microhomology arms corresponding to the genomic DNA sequences immediately 5’ and 3’ of the sgRNA cut site (primers: RepDDX42N_1_P_F 5’-gcgttacatagcatcgtacgcgtacgtgtttggcttattcctaacaggtcagtatgaccgagtacaagcccacg; RepDDX42N_1_P_R 5’-agcattctagagcatcgtaCGCGTACGTGTTTGGtccagttcatggtgccaatgGAagatccgccgccacc). The resulting fragment was gel-purified and re-introduced into an MluI-linearized pCRIS-PITChv2 backbone using NEBuilder 2xHiFi Gibson assembly master mix (New England Biolabs, USA). The sgRNA (6 μg) and repair (6 μg) plasmids were then co-electroporated into 2 million wild-type HCT-116 cells using Lonza Nucleofector II with reagent kit V and program D-032. After recovery for 3 days, selection for successfully tagged clones was initiated by DMEM containing 2 μg/mL puromycin. Surviving colonies appeared after 5 days of puromycin selection and cells were seeded in limiting dilution of 0.1-0.5 cells per 50 μL to four 384-well plates in puro-containing DMEM. After 14 days, wells with a single colony were expanded to 24-well plates and tested for positive dTAG integration via HA-tag and DDX42 antibody western blot. Clones with positive western blot signal were next genotyped by extracting genomic DNA and amplifying the region surrounding the sgRNA cut site (primers DDX42_N_seqF 5’-gcccttggggctatacacttt; DDX42_N_seqR 5’-ccagcactgatggcaaaacc). Clones showing a large band (integrated cassette), but no short band (no integration at the cut site) were selected for Sanger sequencing of the purified PCR reaction. One successful clone was selected and designated the HCT-116 dTAG-DDX42 working clone, which showed homozygous and seamless, indel-free integration of the dTAG cassette. Four more clones with varying integration outcomes (2x seamless, 2x in-frame insertions) were kept as backup clones but not used for any experiments.

*- Generation of sgPSME1, sgPSME2 and sgControl KBM7 cells:* Stable knock-out cell lines were generated by transduction of KBM7_Cas9-P2A-blast (Cas9 lentivirus produced from Addgene #52962) with lentiGuide-puro virus (sgRNAs cloned into Addgene #52963) using standard CRISPR/Cas9 and lentivirus protocols. Briefly, sgRNAs were cloned into lentiGuide-puro via BsmBI restriction and annealed oligo ligation cloning (sgPSME1_sense 5’-CACCGccagcccgaggcccaagcca; sgPSME2_sense 5’ CACCGaaatccagagacttacctcc; sgControl-AAVS1_sense 5’-caccgGGGGCCACTAGGGACAGGAT).

Lentivirus was produced using the polyethylenimine (PEI) protocol (Joung et al., 2017). Briefly, 2 million Lenti-X 293T cells (Takara) were seeded in 5 mL DMEM to 6cm dishes 20h before transfection. The next day, a PEI master mix was prepared (61.8 μL serum-free DMEM, 1.7 μg pSPAX2, 0.85 μg pMD2.G, 1.29 μg expression plasmid DNA per transfection), mixed with 18.5 μL PEI (1 mg/mL in water, neutralized with NaOH to pH 7.0; 40,000 MW) and incubated for 10min at room temperature. The Lenti-X supernatant was changed to 3mL fresh full DMEM and PEI/plasmid mix was added to the medium dropwise. Viral supernatant was harvested once after 70h, sterile filtered using .45 μm syringe filters and frozen in aliquots at -80°C.

For transduction, 1 million KBM7 cells were mixed with 50 μL of freshly thawed sgPSME1, sgPSME2, or sgControl/AAVS1 viral supernatant in a total of 3mL full IMDM supplemented with 8 μg/mL polybrene in 12-well plates. Cells were spin-infected at 2000rpm and 30°C for 1h and incubated for 24h at 37°C. Selection was initiated with 1 μg/mL puromycin and considered complete after 4 days, when the no-virus control cells were completely killed. Selected pools were characterized using gDNA PCR and western blot and used for knock-out cell line experiments.

*- Generation of 22Rv1 cells stably transduced with PHF5A_WT or Y36C:* Lentiviral expression plasmids pLEX304_PHF5A-WT-STOP and pLEX304_PHF5A-Y36C-STOP were cloned using the gateway recombinase cloning strategy (Invitrogen, USA) from commercially synthesized gene blocks (IDT, USA). Lentivirus production and transduction of 22Rv1 cells was performed as described above. 22Rv1 cells stably transduced with pLEX304_PHF5A were selected with 10 μg/mL blasticidin for 7 days, at which point non-transduced control cells were completely killed.

#### Antibodies

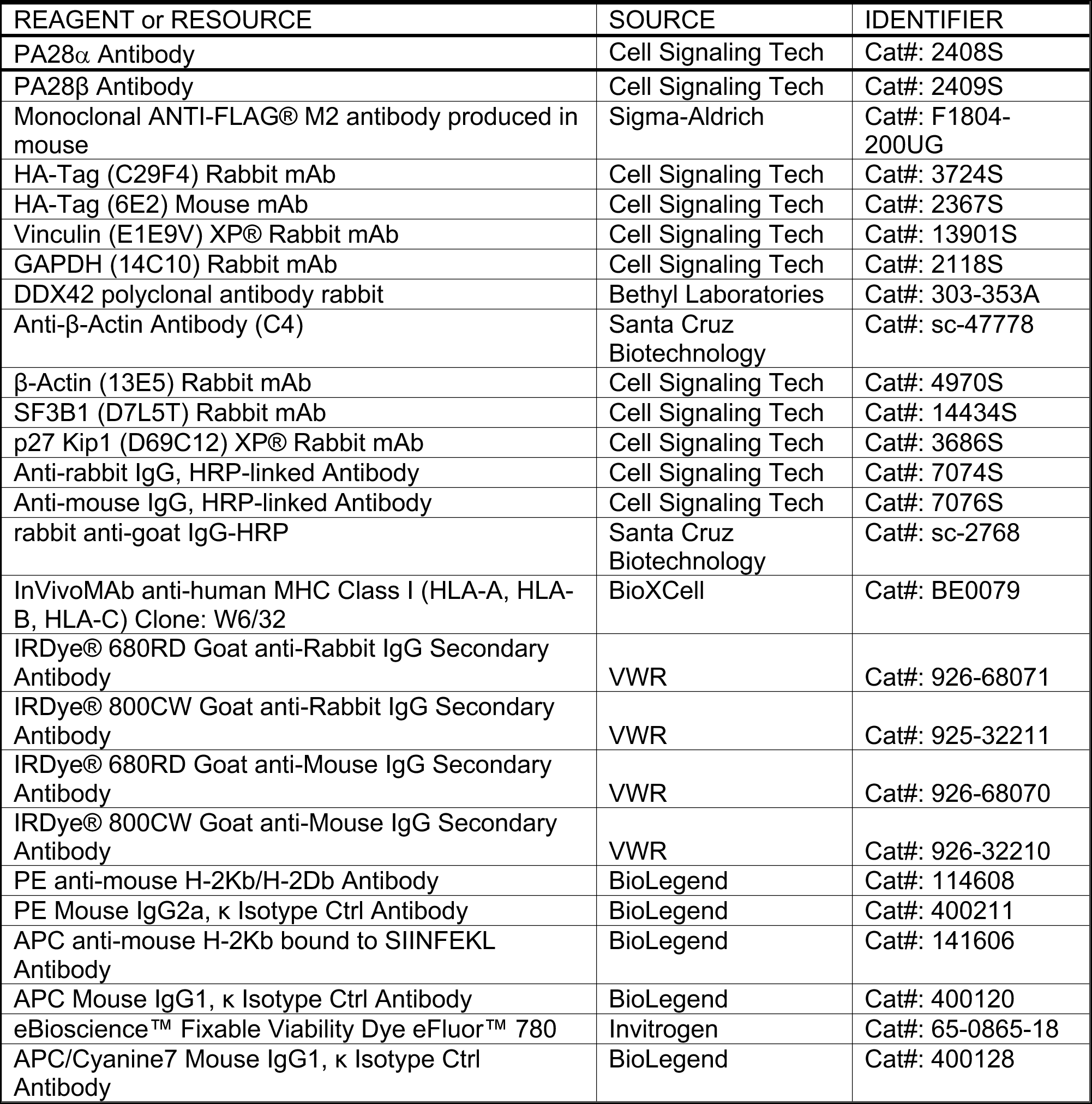

| REAGENT or RESOURCE | SOURCE | IDENTIFIER |
| --- | --- | --- |
| PA28α Antibody | Cell Signaling Tech | Cat#: 2408S |
| PA28β Antibody | Cell Signaling Tech | Cat#: 2409S |
| Monoclonal ANTI-FLAG® M2 antibody produced in mouse | Sigma-Aldrich | Cat#: F1804-200UG |
| HA-Tag (C29F4) Rabbit mAb | Cell Signaling Tech | Cat#: 3724S |
| HA-Tag (6E2) Mouse mAb | Cell Signaling Tech | Cat#: 2367S |
| Vinculin (E1E9V) XP® Rabbit mAb | Cell Signaling Tech | Cat#: 13901S |
| GAPDH (14C10) Rabbit mAb | Cell Signaling Tech | Cat#: 2118S |
| DDX42 polyclonal antibody rabbit | Bethyl Laboratories | Cat#: 303-353A |
| Anti-β-Actin Antibody (C4) | Santa Cruz Biotechnology | Cat#: sc-47778 |
| β-Actin (13E5) Rabbit mAb | Cell Signaling Tech | Cat#: 4970S |
| SF3B1 (D7L5T) Rabbit mAb | Cell Signaling Tech | Cat#: 14434S |
| p27 Kip1 (D69C12) XP® Rabbit mAb | Cell Signaling Tech | Cat#: 3686S |
| Anti-rabbit IgG, HRP-linked Antibody | Cell Signaling Tech | Cat#: 7074S |
| Anti-mouse IgG, HRP-linked Antibody | Cell Signaling Tech | Cat#: 7076S |
| rabbit anti-goat IgG-HRP | Santa Cruz Biotechnology | Cat#: sc-2768 |
| InVivoMAb anti-human MHC Class I (HLA-A, HLA-B, HLA-C) Clone: W6/32 | BioXCell | Cat#: BE0079 |
| IRDye® 680RD Goat anti-Rabbit IgG Secondary Antibody | VWR | Cat#: 926-68071 |
| IRDye® 800CW Goat anti-Rabbit IgG Secondary Antibody | VWR | Cat#: 925-32211 |
| IRDye® 680RD Goat anti-Mouse IgG Secondary Antibody | VWR | Cat#: 926-68070 |
| IRDye® 800CW Goat anti-Mouse IgG Secondary Antibody | VWR | Cat#: 926-32210 |
| PE anti-mouse H-2Kb/H-2Db Antibody | BioLegend | Cat#: 114608 |
| PE Mouse IgG2a, $\kappa$ Isotype Ctrl Antibody | BioLegend | Cat#: 400211 |
| APC anti-mouse H-2Kb bound to SIINFEKL Antibody | BioLegend | Cat#: 141606 |
| APC Mouse IgG1, $\kappa$ Isotype Ctrl Antibody | BioLegend | Cat#: 400120 |
| eBioscience™ Fixable Viability Dye eFluor™ 780 | Invitrogen | Cat#: 65-0865-18 |
| APC/Cyanine7 Mouse IgG1, $\kappa$ Isotype Ctrl Antibody | BioLegend | Cat#: 400128 |

### METHOD DETAILS

#### Proteomic platforms: SEC-MS (related to Figures 1, 2)

*- Sample preparation:* Cells (adherent: 15cm dishes, suspension: 2 million cells/mL) were treated with electrophilic probes *in situ* for indicated times and concentrations (20 μM, 3 h), washed with ice-cold PBS, and flash frozen in liquid nitrogen. Cell pellets were lysed by sonication (8 pulses, 40% power) in 600 μL ice-cold PBS supplemented with cOmplete protease and PhosSTOP phosphatase inhibitors (Roche; 1 tablet/10mL of PBS), ultracentrifuged for 20 min at 100,000 g, then normalized to a standardized concentration (typically 1.5 to 2.5 mg/mL) using a standard DC protein concentration assay (Bio-Rad). 500 μL of clarified lysate was injected into a Superdex 200 Increase 10/300 GL column attached to an ÄKTA pure FPLC system (Cytiva). Proteins were fractionated using an isocratic gradient (PBS) running at 0.5 mL/min into 5 fractions 2 mL wide, beginning at 8 mL and ending at 18 mL. Eluate was collected into 15 mL tubes containing 12 mL of acetone at 4 °C. After completion of one set of 5 fractions, the above protocol was repeated a second time for the remaining 5 fractions. Proteins were precipitated overnight at -20 °C, then centrifuged at 4500 g for 20 minutes, yielding a white protein pellet. Acetone/PBS mixture was decanted off the pellets, which were dried at room temperature and then resuspended in 125 μL of 8 M urea, 10 mM DTT in 100 mM EPPS for 15 minutes at 65 °C, followed by probe sonication to complete resuspension. 6.25 μL of 500 mM iodoacetamide was added to samples for alkylation (30 minutes, 37 °C), followed by dilution

with 370 μL of 100 mM EPPS, and addition of 2 μg trypsin per fraction, and digested overnight at 37 °C. 70 μL of tryptic digest was aliquoted into a clean microcentrifuge tube, followed by 30 μL of dry acetonitrile and 4.5 μL (20 μg/μL) of TMT tag in dry acetonitrile. Samples were labeled at room temperature for 75 minutes, then quenched with 6 μL of 5% hydroxylamine for 15 minutes and acidified with 5 μL of formic acid. The set of 10 fractions were then combined into a single tube and evaporated by speed vac. Samples were desalted via Sep-Pak and then high pH fractionated as described below.

- *Data processing*: Fractional distributions for each peptide-spectra match with a total TMT reporter ion intensity were calculated by dividing the reporter ion intensity for each TMT channel by the summed intensity across the 5 TMT channels corresponding to a single treatment condition (**Equation 1**). Protein-level SEC elution profiles were then generated by averaging together peptide-level elution profiles from unique peptides with summed reporter ion intensities >5,000. Two unique peptide sequences were required per-protein. Bar graph elution profiles are represented as the mean ± standard error of the mean across all replicates. Euclidean distances (SEC shift scores, **Equation 2**) were calculated using the average elution profile for each protein across replicates, combined by treatment condition and cell line. Figures reporting mean elution times used **Equation 3** for calculation. Figures 1E, 1F use **Equation 4** for calculating the delta-SEC shift score for an enantiomeric pair.

- *Filtering of SEC-MS data*: Replicates were combined based on cell line and treatment condition (probe, concentration, duration) by averaging SEC elution profiles for each protein across all experiments. A coefficient of variation (CV) filter was applied by taking the average the per-fraction CV across all 5 fractions. Proteins with a CV > 0.5 were removed from analysis.

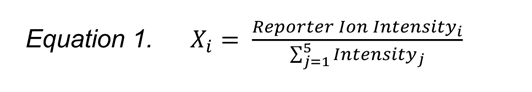

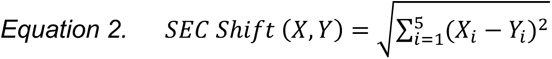

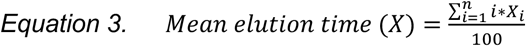

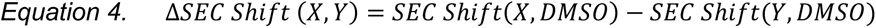

where X_i_, Y_i_ represent protein-level fractional distributions for individual treatment conditions.

#### SEC analysis by Western blot (related to Figure 2)

For experiments using transiently expressed constructs (FLAG-PSME1 WT/C22A), stable clonal 22Rv1 pLenti3.3/TR were transfected using polyethylenimine and treated with tetracycline (0.1 μg/mL) for 48 h prior to treatment with electrophilic probes.

Cells were lysed and fractioned by SEC as described above. After acetone precipitation and resuspension, 4x gel loading buffer was added to eluates, and proteins were resolved using SDS-PAGE (4-20%, Tris-glycine gel), and transferred to a nitrocellulose membrane (350 mA for 90 min). The membrane was blocked with 5% milk in Tris-buffered saline (20 mM Tris-HCl 7.6, 150 mM NaCl) with 0.1% tween (TBST) and incubated with primary antibody overnight at 4 C. After TBST wash (3 times), the membrane was incubated with secondary antibody for 1 h at room temperature, washed with TBST again, developed with ECL western blotting detection reagents, and imaged on a Bio-Rad ChemiDoc MP. Relative band intensities were analyzed using ImageJ.

#### Gel-based ABPP for PSME1 cysteine engagement (related to Figure 3)

C-terminally FLAG-tagged PSME1 WT or C22A were transiently expressed in HEK293T cells, harvested 48 h later, then cells were divided into aliquots, flash frozen and stored at -80 °C. On the day of the experiment, an aliquot of cells were thawed, lysed by sonication in ice-cold PBS supplemented with cOmplete protease inhibitors (1 tablet/10mL of PBS). normalized to 2.0 mg/mL using a standard DC protein assay (Bio-Rad) and divided into 25 μL aliquots. 1 μL of DMSO or compound (25x stock) was incubated with lysates for 2 h at RT, followed by incubation with 1 μL of 62.5 μM MY-11B click probe (final concentration 2.4 μM) for an additional 30 minutes. Reagents for the CuAAC click reaction were pre-mixed prior to addition to the samples, as described previously (Backus *et al*., 2016). After 1 h of click labeling, 4x SDS loading buffer was added to samples, which were then analyzed by SDS-PAGE. ImageJ was used to quantify rhodamine band intensities, and GraphPad PRISM Version 9.0.0 was used to generate IC_50_ curves (four-parameter variable slope least squares regression)

#### FACS analysis with citric acid wash (related to Figure 3)

E.G7-Ova cells (2 million/mL) were treated with compounds for 4 h, washed with PBS, and then washed with 1 mL of mild acid elution buffer (131 mM citric acid, 66 mM NaH_2_PO_4_; pH 3.0) for 2 minutes, and then washed with media three times. Cells were then resuspended in warm RPMI and allowed to recover for the indicated period, after which cells were washed with PBS and transferred to 96 well plates for staining. Each well was washed with 200 μL PBS, and then stained for 20 minutes with 50 μL of staining solution: 1:1000 fixable viability dye stain (Invitrogen) and 1:200 dilution of anti-SIINFEKL (BioLegend) or anti-mouse H-2K^b^ (BioLegend) antibody in FACS buffer (2% FBS in PBS supplemented with PenStrep and sterile filtered). Additional wells were used as unstained or singly-stained controls. After staining, cells were washed with PBS, then fixed in 200 μL 4% paraformaldehyde in PBS for 15 minutes. Cells were resuspended in 75 μL FACS buffer and analyzed on a Novocyte flow cytometer. The corresponding data in Figure 3G are presented as the average percentage of median fluorescence intensity (MFI) relative to DMSO (4 h timepoint) treated control ± SD from n = 4 replicates. The corresponding data in Figure 3H are presented as the average percentage of MFI relative to DMSO treated control ± SD from n = 3 replicates. Statistical analysis was performed using GraphPad PRISM Version 9.0.0.

#### Proteomic platform: Cysteine ligandability profiling (Related to Figures 2, 3)

- *Sample preparation*: Cells (adherent: 15 cm dishes, suspension: 2 million cells/mL) were treated with electrophilic probes *in situ* for indicated times and concentrations, washed with ice-cold PBS, and flash frozen in liquid nitrogen. Cell pellets were lysed by sonication (8 pulses, 40% power) in 500 μL ice-cold PBS supplemented with cOmplete protease inhibitors (1 tablet/10 mL of PBS). Whole cell lysates protein content was measured using a standard DC protein assay (Bio-Rad) and 500 μL (2 mg/mL protein content) were treated with 5 μL of 10 mM IA-DTB (in DMSO) for 1 h at ambient temperature with occasional vortexing. Samples were methanol-chloroform precipitated with the addition of 500 μL ice-cold MeOH and 200 μL CHCl3, vortexed, and centrifuged (10 min, 10,000 g). Without disrupting the protein disk, both top and bottom layers were aspirated, and the protein disk was washed with 1 mL ice-cold MeOH and centrifuged (10 min, 10,000 g). The pellets were allowed to air dry, and then re-suspended in 90 μL buffer (9 M urea, 10 mM DTT, 50 mM TEAB pH 8.5). Samples were reduced by heating at 65 °C for 20 minutes and water bath sonicated as needed to resuspend the protein pellets, followed by alkylation via addition of 10 μL (500 mM) iodoacetamide and incubated at 37 °C for 30 min with shaking. Samples were diluted with 300 μL buffer (50 mM TEAB pH 8.5) to reach final concentration of 2 M urea, briefly centrifuged, and probe sonicated once more to ensure complete resuspension. Trypsin (4 μL of 0.25 μg/μL in trypsin resuspension buffer with 25 mM CaCl_2_) was added to each sample and digested at 37 °C for 2 h to overnight. Digested samples were then diluted with 300 μL wash buffer (50 mM TEAB pH 8.5, 150 mM NaCl, 0.2% NP-40) containing streptavidin-agarose beads (50 μL of 50% slurry in wash buffer) and were rotated at room temperature for 2 h. Enriched samples were pelleted by centrifugation (2000 g, 2 min) and transferred to BioSpin columns and washed (3x1 mL wash buffer, 3x1 mL PBS, 3x1mL water). Enriched peptides were eluted with 300 μL of 50% acetonitrile with 0.1% formic acid and eluate evaporated to dryness via speedvac. IA-DTB labeled and enriched peptides were resuspended in 100 μL EPPS buffer (200 mM, pH 8.0) with 30% acetonitrile, vortexed, and water bath sonicated. Samples were TMT labeled with 3 μL of corresponding TMT tag (20 μg/μL), vortexed, and incubated at room temperature for 1 h. TMT labeling was quenched with the addition of hydroxylamine (5 μL 5% solution in H_2_O) and incubated for 15 minutes at room temperature. Samples were then acidified with 5 μL formic acid, combined and dried using a SpeedVac. Samples were desalted via Sep-Pak and then high pH fractionated as described below.

- *Data processing*: Cysteine engagement ratios (probe vs DMSO) were calculated for each peptide-spectra match by dividing each TMT reporter ion intensity by the average intensity for the channels corresponding to DMSO treatment. Peptide-spectra matches were then grouped based on protein ID and residue number (e.g., PSME1 C22), excluding peptides with summed reporter ion intensities for the DMSO channels < 10,000, coefficient of variation for DMSO channels > 0.5, and non-unique or non-tryptic peptide sequences. TMT reporter ion intensities were normalized to the median summed signal intensity across channels.

- *Filtering of cysteine data*: Replicates were combined based on cell line and treatment condition (probe, concentration, duration) by averaging cysteine engagement ratios across all experiments.

#### Proteomic platform: Immunopeptidomics (related to Figure 4)

- *Sample preparation*: KBM7 cells (10 million cells/mL, 500 million cells per sample) were treated with electrophilic probes (from 50 mM stocks) as indicated for 8 h, washed with ice-cold PBS, and flash frozen in liquid nitrogen. MHC enrichments were processed as previously described (Purcell et al., 2019). Briefly, cells were lysed with 3 mL of lysis buffer (0.5% (vol/vol) NP-40 (Igepal CA-630), 50 mM Tris pH 8.0, 150 mM NaCl, 1 mM EDTA, 0.2 mM Iodoacetamide, and protease inhibitor cocktail (Roche)), rotating at 4 C for 30 minutes in a 5 mL low-bind Eppendorf. Samples were centrifuged at 18,000 g for 10 minutes and supernatant was taken for enrichment by anti-MHC antibody (BioXCell, BE0079) conjugated to Affi-gel 10 matrix (∼2 mg of antibody per sample). Enrichments rotated at 4 C for 2 h and transferred to Bio-spin filter column for washing with 3x1 mL lysis buffer, wash buffer 1 (50 mM Tris pH 8, 150 mM NaCl), wash buffer 2 (50 mM Tris pH 8, 400 mM NaCl), and wash buffer 3 (50 mM Tris pH 8). MHC-conjugated peptides were eluted with 1 mL of 10% acetic acid in HPLC grade water. Samples were immediately desalted using Sep-Pak, dried via speedavac, and then ready for injection into the mass spectrometer.

- *Mass spectrometry*: Samples were analyzed by liquid chromatography tandem mass-spectrometry using an Orbitrap Eclipse Tribrid Mass Spectrometer (Thermo Scientific) coupled to an UltiMate 3000 Series Rapid Separation LC system and autosampler (Thermo Scientific Dionex). The peptides were eluted onto an EASY-Spray HPLC column (Thermo ES902, ES903) using an Acclaim PepMap 100 (Thermo 164535) loading column, and separated at a flow rate of 0.25 μL/min. Peptides were separated across a 90 minute gradient of 2-15%, 5 minutes 15-20%, and then 5 minutes 20-95% acetonitrile (0.1% formic acid) followed by column equilibration. Data was acquired using a data-dependent acquisition in profile mode. Briefly, the scan sequence began with an MS1 master scan (Orbitrap analysis, resolution 120,000, 350-1400 m/z, RF lens 30%, automatic gain control [AGC] target 250%, maximum injection time 50 ms, profile mode) with dynamic exclusion enabled (repeat count 1, duration 15 s). The top precursors were then selected for MS2 analysis within a 2.5 duty cycle. MS2 analysis consisted of quadrupole isolation (isolation window 0.7) of precursor ion followed by collision-induced dissociation (CID) in the orbitrap (AGC 300%, normalized collision energy 35%, maximum injection time 250 ms). Raw files were searched using MaxQuant (v2.0.3.1) with 5% PSM FDR again the Human UniProt database (release 2016), unspecific digestion, 8-12 amino acids in length, 20 ppm peptide tolerance, and match-between-runs enabled (1 minute match time window, 20 minute alignment time window). Subsequent results were further filtered to require an identification posterior error probability < 0.01.

#### Proteomic platform: Whole proteome profiling (related to Figures 5, 6)

- *Sample preparation*: Cells (adherent: 15cm dishes, suspension: 2 million cells/mL) were treated with electrophilic probes *in situ* for indicated times and concentrations (1 h or 3 h as noted), washed with ice-cold PBS, and flash frozen in liquid nitrogen. Cell pellets were lysed by sonication (8 pulses, 40% power) in 200 μL ice-cold PBS supplemented with cOmplete protease inhibitors (1 tablet/10 mL of PBS) and 1 mM PMSF (100 mM stock in ethanol). Whole cell lysates protein content was measured using a standard DC protein assay (Bio-Rad) and a volume corresponding to 200 μg was transferred to a new low-bind Eppendorf tube (containing 48 mg urea) and brought to 100 μL total volume. Samples were reduced with DTT (5 μL 200 mM stock in H_2_O, 10 mM final concentration) and incubated at 65 °C for 15 minutes, then alkylated with iodoacetamide (5 μL 400 mM stock in H_2_O, 20 mM final concentration) incubated at 37 °C for 30 minutes in the dark. Samples were precipitated with the addition of ice-cold MeOH (600 μL), CHCl3 (180 μL), and H_2_0 (500 μL), and vortexed and centrifuged (16,000 g, 10 minutes, 4 °C). The top layer above the protein disc was aspirated and an additional 1mL of ice-cold MeOH was added. The samples were again vortexed and centrifuged (16,000 g, 10 minutes, 4 °C), the supernatant aspirated and the protein pellets were allowed to air dry and be stored at -80 °C or proceeded to resuspension and digestion. Samples were resuspended in 160 μL EPPS buffer (200 mM pH 8.0) using a probe sonicator (10-15 pulses). Proteomes were first digested with LysC (0.5 μg/μL in HPLC grade water, 4 μL per sample) for 2 h at 37 °C. Then trypsin (8 μL per sample, 0.5 μg/μL in trypsin resuspension buffer with 20 mM CaCl_2_) was added and samples were incubated at 37 °C overnight. Peptide concentrations were estimated using Micro BCA^TM^ assay (Thermo Scientific), and a volume corresponding to 25 μg was transferred to a new low-bind Eppendorf tube and brought to 35 μL with EPPS buffer. Samples were diluted with 9 μL acetonitrile and then TMT labeled with 5 μL of corresponding TMT tag (20 μg/μL), vortexed, and incubated at room temperature for 1 h. TMT labeling was quenched with the addition of hydroxylamine (5 μL 5% solution in H_2_O) and incubated for 15 minutes at room temperature. Samples were then acidified with 2.5 μL formic acid and an equal volume (20 μL corresponding to ∼8.85 μg per channel) was combined and dried using a SpeedVac. Samples were desalted via Sep-Pak and then high pH fractionated as described below.

- *Data processing*: Protein abundance ratios (probe vs DMSO) were calculated for each peptide-spectra match by dividing each TMT reporter ion intensity by the average intensity for the channels corresponding to DMSO treatment. Peptide-spectra matches were then grouped based on protein ID, excluding peptides with summed reporter ion intensities for the DMSO channels < 10,000, coefficient of variation for DMSO channels > 0.5, non-unique or non-tryptic peptide sequences. TMT reporter ion intensities were normalized to the median summed signal intensity across channels.

- *Filtering of whole proteome data*: Whole proteome data was filtered at peptide-level by removing any peptide-spectra matches with a standard deviation of abundance ratio > 100%. At a replicate-level, proteins with at least 2 distinct peptide-spectra matches were retained for analysis. Replicates were then combined based on cell line and treatment condition (probe, concentration, duration) by averaging protein abundance ratios across all experiments.

#### Offline fractionation

High pH offline fractionation was performed as previously described (Remsberg et al., 2021; Vinogradova *et al*., 2020). Briefly, samples fractionated via Peptide Desalting Spin Columns (Thermo 89852) were resuspended in buffer A (5% acetonitrile, 0.1% formic acid) and bound to the spin columns. Bound peptides were then washed in water, 10 mM NH_4_HCO_3_ containing 5% acetonitrile, and eluted in fractions of increasing acetonitrile, concatenated, and then dried using a SpeedVac vacuum concentrator. Resulting fractions were resuspended in buffer A (5% acetonitrile, 0.1% formic acid) and analyzed by mass spectrometry.

Samples fractionated by HPLC were resuspended in buffer A (500 μL) and fractionated into a 96 deep-well plate using HPLC (Agilent). The peptides were eluted onto a capillary column (ZORBAX 300Extend-C18, 3.5 μm) and separated at a flow rate of 0.5 mL/min using the following gradient: 100% buffer A from 0-2 min, 0%–13% buffer B from 2-3 min, 13%–42% buffer B from 3-60 min, 42%–100% buffer B from 60-61 min, 100% buffer B from 61-65 min, 100%–0% buffer B from 65-66 min, 100% buffer A from 66-75 min, 0%–13% buffer B from 75-78 min, 13%–80% buffer B from 78-80 min, 80% buffer B from 80-85 min, 100% buffer A from 86-91 min, 0%–13% buffer B from 91-94 min, 13%–80% buffer B from 94-96 min, 80% buffer B from 96-101 min, and 80%–0% buffer B from 101-102 min (buffer A: 10 mM aqueous NH_4_HCO_3_; buffer B: acetonitrile). Each well in the 96-well plate contained 20 μL of 20% formic acid to acidify the eluting peptides. The eluent was evaporated to dryness in the plate using SpeedVac vacuum concentrator. The peptides were resuspended in 80% acetonitrile, 0.1% formic acid buffer (125 μL/column) and every 12th fraction was combined. Samples were dried using SpeedVac vacuum concentrator, the resulting 12 combined fractions were re-suspended in buffer A (5% acetonitrile, 0.1% formic acid) and analyzed by mass spectrometry.

#### Parallel reaction monitoring (Related to Figure 6)

Dry peptide samples were reconstituted in a water/acetonitrile (85:15) mixture containing 0.1% formic acid (100 µL) and 15 µL was injected on to an EASY-Spray C18 loading column (5 µm particle size, 100 µm × 2 cm; Fisher Scientific, DX164564) and resolved on a custom analytical column (2 µM particle size, 75 µm x 15 cm) using a Dionex Ultimate 3000 nano-LC (Thermo Fisher Scientific). Peptides were separated over a 60-min gradient of 15 to 33% acetonitrile (0.1% formic acid) and analyzed on a Q-Exactive instrument (Thermo Fisher Scientific) using a parallel reaction monitoring method targeting the SF3B1 peptide containing C1111 (amino acids 1110 - 1149, missed tryptic, +3 charge state). Selected precursor ions were isolated and fragmented by high-energy collision dissociation and fragments were detected in the Orbitrap at 17,500 resolution.

Raw data files were uploaded analyzed in Skyline (v.21.1.0.278) to determine the abundance of each peptide in inhibitor-treated samples relative to vehicle-treated samples. Peptide quantification was performed by calculating the sum of the peak areas corresponding to six fragment ions from each peptide. The peptides and fragment ions were preselected from in-house reference spectral libraries acquired in data-dependent acquisition mode to identify authentic spectra for each peptide and normalized to internal retention time standards.

#### TMT liquid chromatography-mass-spectrometry (LC-MS) analysis

Samples were analyzed by liquid chromatography tandem mass-spectrometry using an Orbitrap Fusion Tribrid Mass Spectrometer (Thermo Scientific) or an Orbitrap Eclipse Tribrid Mass Spectrometer coupled to an UltiMate 3000 Series Rapid Separation LC system and autosampler (Thermo Scientific Dionex). The peptides were eluted onto a capillary column (75 μm inner diameter fused silica, packed with C18 (Waters, Acquity BEH C18, 1.7 μm, 25 cm)) or an EASY-Spray HPLC column (Thermo ES902, ES903) using an Acclaim PepMap 100 (Thermo 164535) loading column, and separated at a flow rate of 0.25 μL/min. Data was acquired using an MS3-based TMT method on Orbitrap Fusion or Orbitrap Eclipse Tribrid Mass Spectrometers. Briefly, for Fusion acquisition, the scan sequence began with an MS1 master scan (Orbitrap analysis, resolution 120,000, 400−1700 m/z, RF lens 60%, automatic gain control [AGC] target 2E5, maximum injection time 50 ms, centroid mode) with dynamic exclusion enabled (repeat count 1, duration 15 s). The top ten precursors were then selected for MS2/MS3 analysis. MS2 analysis consisted of quadrupole isolation (isolation window 0.7) of precursor ion followed by collision-induced dissociation (CID) in the ion trap (AGC 1.8E4, normalized collision energy 35%, maximum injection time 120 ms). Following the acquisition of each MS2 spectrum, synchronous precursor selection (SPS) enabled the selection of up to 10 MS2 fragment ions for MS3 analysis. MS3 precursors were fragmented by HCD and analyzed using the Orbitrap (collision energy 55%, AGC 1.5E5, maximum injection time 120 ms, resolution was 50,000). For MS3 analysis, we used charge state–dependent isolation windows. For charge state z = 2, the MS isolation window was set at 1.2; for z = 3-6, the MS isolation window was set at 0.7. Scan parameters on Eclipse Tribrid Mass Spectrometers were the same with the following exceptions; full MS scan used RF lens 30%, standard AGC target and auto maximum injection time. MS2 scans in the Ion Trap used normalized collision energy 36%, standard AGC target and auto maximum injection time. MS3 SPS analysis used 30,000 resolution and 500% normalized AGC target and auto maximum injection time. The MS2 and MS3 files were extracted from the raw files using RAW Converter (version 1.1.0.22; available at http://fields.scripps.edu/rawconv/), uploaded to Integrated Proteomics Pipeline (IP2), and searched using the ProLuCID algorithm (publicly available at http://fields.scripps.edu/downloads.php) using a reverse concatenated, non-redundant variant of the Human UniProt database (release 2016). Cysteine residues were searched with a static modification for carboxyamidomethylation (+57.02146 Da). For cysteine profiling experiments, a dynamic modification for IA-DTB labeling (+398.25292 Da) was included with a maximum number of 2 differential modifications. N-termini and lysine residues were also searched with a static modification corresponding to the TMT tag (+229.1629 Da). Peptides were required to be at least 6 amino acids long, and to be fully tryptic, except for the GluC digested cysteine profiling samples which included K, R, E, and D cleavage sites. ProLuCID data was filtered through DTASelect (version 2.0) to achieve a peptide false-positive rate below 1%. The MS3-based peptide quantification was performed with reporter ion mass tolerance set to 20 ppm with Integrated Proteomics Pipeline (IP2).

#### Gene Ontology analysis (related to Figure 5)

A list of stereoselectively affected proteins was analyzed using GOnet tools(Pomaznoy et al., 2018) (ontology version 2019-07-01) for GO term enrichment of biological processes using hierarchical output.

#### Proliferation and apoptosis assays (related to Figures 5-7)

Cells were seeded at 5000 cells per well (50 μL medium) in 96-well flat bottom white wall plates. After 24 h, 50 μL of medium containing DMSO or compound dilutions (2x final concentration from 1000x stocks) were added to the wells. At the time of compound addition, a reference plate to determine the cell population density at time 0 (*T*_0_) was assayed using CellTiter-Glo (Promega) reagent (50 μL added to each well). After 72 h in culture, the remaining plates were assayed using CellTiter-Glo. After 30 min of shaking at room temperature, luminescence was analyzed using a CLARIOstar (BMG Labtech) plate reader. Apoptosis assays were performed similarly as described above using Caspase-Glo 3/7 (Promega) following 24 h compound treatments. Raw values were normalized using GraphPad PRISM software version 9.0.0. Lines of best fit were generated using a four-parameter variable slope least squares regression. Proliferation assays linked to HCT-116 DDX42-dTAG cells were performed similarly, except that compound was prepared in a new 96-well plate as a 1:sqrt(10) serial dilution with 9 dose points (plus 1 untreated well) and adding 50 μL of this drug/medium solution to the 50 μL cells.

#### MS-based ABPP for SF3B1 cysteine engagement (related to Figure 5)

22Rv1 cells were treated with indicated compounds for 2 h, chased with click probe for 1 h *in situ*, harvested and lysed in PBS with protease inhibitors (cOmplete), and then normalized to 2.0 mg/mL. Reagents for the CuAAC click reaction were pre-mixed prior to addition to the samples (55 μL of click reaction mix for 500 μL of lysate), as described previously described (Wang et al., 2019). After 1 h of click labeling, samples were precipitated with the addition of ice-cold MeOH (600 μL), CHCl3 (180 μL), and H_2_0 (500 μL), and vortexed and centrifuged (16,000 g, 10 minutes, 4 °C). The top layer above the protein disc was aspirated and an additional 1mL of ice-cold MeOH was added. The samples were again vortexed and centrifuged (16,000 g, 10 minutes, 4 °C), the supernatant aspirated and the protein pellets were allowed to air dry and be stored at -80 °C or proceeded to resuspension and enrichment. Pellets were resuspended in 500 µL 8 M urea (in DPBS) with 10 µL 10% SDS and sonicated to ensure no large precipitate remained. Samples were reduced with 25 µL of 200 mM DTT and incubated at 65 °C for 15 minutes, then reduced with addition of 25 µL of 400 mM iodoacetamide and incubated in the dark at 37 °C for 30 minutes. For enrichment, 130 µL 10% SDS was added and then samples were diluted to 5.5 mL DPBS. Samples were enrichment with streptavidin beads (Thermo cat # 20353; 100 µL/sample) and incubated at room temperature for 1.5 h while rotating. After incubation, samples were pelleted by centrifugation (2,000 g, 2 min) and beads were washed with 0.2% SDS in DPBS (2 x 10 mL), DPBS (1 x 5 mL), then transferred to low-bind Eppendorf (cat # 0030108442), washed with water (2 x 1 mL), and 200 mM EPPS (1mL). Pelleted beads were resuspended in 200 µL 2 M urea (in 200 mM EPPS) and digested with 2 µg Trypsin with 1 mM CaCl_2_ (final concentration) overnight at 37 °C. Digested peptides were transferred to new low-bind Eppendorf and acetonitrile added (to 30% final) and TMT labeled with 6 μL of corresponding TMT tag (20 μg/μL), vortexed, and incubated at room temperature for 1 h. TMT labeling was quenched with the addition of hydroxylamine (6 μL 5% solution in H_2_O) and incubated for 15 minutes at room temperature. Samples were then acidified with 15 μL formic acid, combined and dried using a SpeedVac. Samples were desalted and high pH fractionated using Peptide Desalting Spin Columns (Thermo 89852) to a final of 3 fractions and analyzed by mass spectrometry.

#### Gel-based ABPP for SF3B1 cysteine engagement (related to Figure 6)

22Rv1 or HCT-116 cells were treated with indicated compounds for 2-24 h, chased with click probe for 1 h *in situ*, harvested and lysed in PBS with protease inhibitors (cOmplete), and then normalized to 2.0 mg/mL. Reagents for the CuAAC click reaction were pre-mixed prior to addition to the samples (3 μL of click reaction mix for 25 μL of lysate), as previously reported (Backus *et al*., 2016). After 1 h of click labeling, 4x SDS loading buffer was added to samples, which were then analyzed by SDS-PAGE.

- *IP-ABPP*: HCT-116 DDX42-dTAG cells were treated *in situ* with DMSO or dTAGv-1 (500 nM) for 1 h. The samples were then treated with WX-01-10 or WX-01-12 (2.5 μM) for 1 h. The cells were washed with ice-cold DPBS and then collected via scraping. The pellets were flash frozen in liquid nitrogen and stored at -80 °C until analysis. The pellets were suspended in 0.5 mL DPBS containing 1.5 mM MgCl_2_, 1% NP40, and 1 mM PMSF (1% DMSO v/v final). The cells were lysed via sonication (1 x 10, 10%) on ice and then treated with Pierce Universal nuclease (1 μL). The samples were incubated for 1.5 h at 4 °C with rotation. The samples were then re-sonicated (1 x 10, 10%) and centrifuged at 16,000 RPM for 10 min. The supernatant was quantified via DC assay (BioRad) and normalized to 2 mg/mL total (550 μL each). 50 μL input was collected and treated with CuAAC click mixture for 1 h at room temperature and then it was quenched with 4x loading buffer (store at -20 °C prior to analysis). The remaining sample was treated with 1 μg rabbit anti-SF3B1 overnight at 4 °C with rotation. Then 25 μL Pierce Protein A plus resin (pre-washed in DPBS supplemented with 0.2% NP40, suspended in 75 μL final) was added to each sample and allowed to incubate at 4 °C for 4 h. The beads were washed 3 x 1 mL with ice-cold DPBS containing 0.2% NP40 (pellet at 2000 rpm for 2 min to pellet beads). The beads were suspended in 50 μL DPBS and treated with CuAAC click mixture for 1 h at room temperature. The SF3B1 was eluted by addition of 4x loading buffer and then heated for 10 min at 95 °C. The supernatant was collected and analyzed by SDS-PAGE. proteins were resolved using SDS-PAGE (10% Tris-glycine gel) and transferred to a methanol pre-activated PVDF membrane (60 V for 2 h). The membrane was blocked with 5% milk in Tris-buffered saline (20 mM Tris-HCl 7.6, 150 mM NaCl) with 0.1% tween (TBST) and incubated with primary antibody overnight. After TBST wash (3 times), the membrane was incubated with secondary antibody (mouse anti-rabbit, sc-2357), washed with TBST again, developed with ECL western blotting detection reagents (SuperSignal™ West Pico PLUS Chemiluminescent Substrate), and recorded on a Bio-Rad ChemiDoc MP.

#### RNA sequencing and analysis of differential gene expression (related to Figures 6 and 7)

Total RNA from compound or DMSO treated 22Rv1 or HCT-116 DDX42-dTAG cells was isolated using QIAshredder and RNeasy Mini Plus Kits (QIAGEN) according to manufacturer’s protocol and stored at -80 °C until further analysis. RNA sequencing libraries were prepared using the NEBNext Ultra II RNA Library Prep Kit for Illumina following manufacturer’s instructions (NEB, Ipswich, MA, USA). Briefly, mRNAs were first enriched with Oligo(dT) beads. Enriched mRNAs were fragmented for 15 minutes at 94 °C. First strand and second strand cDNAs were subsequently synthesized. cDNA fragments were end repaired and adenylated at 3’ends, and universal adapters were ligated to cDNA fragments, followed by index addition and library enrichment by limited-cycle PCR. The sequencing libraries were validated on the Agilent TapeStation (Agilent Technologies, Palo Alto, CA, USA), and quantified by using Qubit 2.0 Fluorometer (Invitrogen, Carlsbad, CA) as well as by quantitative PCR (KAPA Biosystems, Wilmington, MA, USA). FASTQ files were first trimmed using Trim_galore (v0.6.4) to remove sequencing adapters and low quality (Q<15) reads. Trimmed sequencing reads were aligned to the human Hg19 reference genome (GENCODE, GRCh37.p13) using STAR (v2.7.5) (Dobin et al., 2013). SAM files were subsequently converted to BAM files, sorted, and indexed using samtools (v1.9). BAM files were used to generate bigwig files using bamCoverage (part of the Deeptools package; v3.3.1). Read counting across genomic features was performed using featureCounts (part of the subread package; v1.5.0) (Liao et al., 2014) using the following parameters: -p -T 20 -O -F GTF -t exon. Differential gene expression analysis was performed using the edgeR (v3.32.1), DESeq2 (v1.30.1) (Love et al., 2014), and limma voom (v3.46.0) (Law et al., 2014) R packages. Data visualization and figure generation was performed in Rstudio (v1.3.1073) using the following packages: ggplot2 (v3.3.5), ggpubr (v0.4.0), complexHeatmap (v2.6.2), and VennDiagram (v1.6.0).

For quantification of alternative RNA splicing, fastq files are adapter-trimmed by cutadapt 3.4 and aligned to GRCh38 by STAR 2.7.6 (Dobin *et al*., 2013) and analyzed using rMATS (v4.1.1) (Shen et al., 2014) using the GENCODE (v35) GTF annotation for GRCh38 and the following parameters: -t paired --libType fr-unstranded –readLength 150 --novelSS. Enumeration of isoform counts was performed using only reads that span the splice junction directly. To identify high confidence AS events, events were considered significant if (i) the inclusion level difference was greater than 10% compared to DMSO, (ii) the False Discovery Rate (FDR) was smaller than 0.05. For comparison of splicing events between conditions, we further limited to highly covered events where at least a total of 100 junctional reads cover the isoform for all conditions. The Inclusion level difference for each AS event were used to cluster the conditions. Data analysis and visualization was performed using custom scripts in python (3.7.12), pandas, scikit-learn, seaborn, Rstudio (v1.3.1073) using the following packages: ggplot2 (v3.3.5), ggrepel (v0.9.1), maser (v1.8.0), and VennDiagram (v1.6.0).

- *Branchpoint analysis*: Branchpoint analysis was performed using Branchpointer (https://github.com/signalbash/branchpointer) (Signal et al., 2018), which is a machine learning model trained using sequence feature surrounding the 3′ splice site. To identify branchpoint associated with each isoform, we found -18 to -44 nucleotide upstream of 3′ splice site and fed into the model. We filtered with the suggested scoring threshold (prediction probability > 0.52) from the software for high confidence branchpoints. U2 binding energy predicted by branchpointer was then used to compare the difference between isoforms.

#### SF3B1 Co-immunoprecipitation studies (related to Figure 6 and 7)

HEK293T or HCT-116 DDX42-dTAG cells were treated *in situ* with DMSO, 5 μM of WX-02-23/WX-02-43, or 10 nM pladienolide B for 3 h. Cells were harvested, washed with ice-cold PBS, and lysed in CoIP lysis buffer (0.5% NP-40, 100 mM EPPS, 150 mM NaCl, cOmplete protease inhibitor). Lysates were clarified at 16,000 g for 3 minutes, normalized to 2.0 mg/mL and 500 μL of each lysate was aliquoted to a clean eppendorf tube.

Lysates were incubated with 5 μL of anti-IgG or anti-SF3B1 (CST) for 2 h at 4 °C, followed by 1 h with 25 μL of protein-A agarose beads. Beads were washed twice with CoIP lysis buffer, and then twice with 100 mM EPPS, followed by elution with 8 M urea in EPPS at 65 °C for 10 min. Eluates were reduced with DTT at 65 °C for 15 min (2.5 μL of 200 mM = 12.5 mM final), alkylated with iodoacetamide at 37 °C for 30 min (2.5 μL of 400 mM = 25 mM final), and diluted to 2 M urea by addition of 115 μL of EPPS. Samples were then trypsinized at 37 °C overnight (2 μg of trypsin per sample), then TMT labeled and desalted as described above.

A similar experiment was performed, where HCT-116 DDX42-dTAG cells were treated with 100 nM pladienolide B and the compound was included in the lysis buffer during lysis and immunoprecipitation to assess if any observed interactome differences between WX-02-23 and pladienolide B were due to washout of the reversibly binding pladienolide B during cell lysis/IP.

#### Enhanced CLIP-seq (related to Figure 7)

- *Library preparation*: 5 million cells were seeded in 10 mL media in a 10 cm plate and grown for 20h. Media was then aspirated and replaced with 7 mL media containing the indicated compounds or 0.1% DMSO for 3h. Following this incubation period, media was aspirated, cells were washed with PBS and protein–RNA interactions were stabilized with UV crosslinking (254 nm, 400 mJ/cm2) in cold DPBS. Cells were scraped, harvested and pelleted in 1.5 mL tubes (2000 g, 3 min, 4°C). Cell pellets were flash-frozen in liquid N2 and stored at -80°C until further analysis. Thawed pellets were then lysed in 1 mL of iCLIP lysis buffer and digested with RNase I (Ambion). Immunoprecipitation of HA-DDX42–RNA complexes with HA-Tag (C29F4) Rabbit mAb (10 μg per mL of lysate) was performed using magnetic beads with pre-coupled secondary antibody (M-280 Sheep Anti-Rabbit IgG Dynabeads, ThermoFisher Scientific 11204D) and beads were stringently washed. After dephosphorylation with FastAP (ThermoFisher) and T4 PNK (NEB), a barcoded RNA adapter was ligated to the 3′ end (T4 RNA Ligase, NEB). Ligations were performed on-bead (to allow washing away unincorporated adapter) in high concentration of PEG8000. Samples were then run on standard protein gels and transferred to nitrocellulose membranes, and a region 75 kDa (∼220 nt of RNA) above the protein size was isolated and proteinase K (NEB) treated to isolate RNA. RNA was reverse transcribed with Superscript III (Invitrogen), and treated with ExoSAP-IT (Affymetrix) to remove excess oligonucleotides. A second DNA adapter (containing a random-mer of 10 (N10) random bases at the 5′ end) was then ligated to the cDNA fragment 3′ end (T4 RNA Ligase, NEB), performed with high concentration of PEG8000 (to improve ligation efficiency) and DMSO (to decrease inhibition of ligation due to secondary structure). After cleanup (Dynabeads MyOne Silane, ThermoFisher), an aliquot of each sample was first subjected to qPCR (to identify the proper number of PCR cycles), and then the remainder was PCR amplified (Q5, NEB) and size selected via agarose gel electrophoresis. Samples were sequenced on the Illumina NovaSeq 6000 platform as two single-end 75bp reads.

- *Data analysis*: After standard NovaSeq demultiplexing, eCLIP libraries processed with Skipper (https://github.com/YeoLab/skipper/) (Boyle *et al*., 2022). Briefly, reads were adapter trimmed with skewer (Jiang et al., 2014) (https://github.com/relipmoc/skewer) and reads less than 20 bp were discarded. UMI were extracted using fastp 0.11.5 (https://github.com/OpenGene/fastp) (Chen et al., 2018). Mapping was then performed against the full human genome (hg38) including a database of splice junctions with STAR (v 2.7.6) (Dobin *et al*., 2013), allowing up to 100 multimapped regions. When multimapping occurred, reads are randomly assigned to any of the matching position. Reads were then PCR duplicated using UMIcollaps (https://github.com/Daniel-Liu-c0deb0t/UMICollapse). Enriched “windows” (IP versus SM-Input) was called on deduplicated reads using a GC-bias aware beta-binomial model. Each window (∼100 b.p.) were partitioned from Gencode v38 and associated with a specific type of genomic region (CDS, UTR, proximal introns near splice site… etc). Windows are filtered using FDR < 0.2 and only reproducible windows between two replicates are used.

- *Differential binding analysis*: To analyze how WX-02-23, WX-02-43 and PladB change DDX42 binding, we ran skipper with the same method as described above but replaced the SM-Input library with the DMSO IP library. This way, the output from skipper represents windows that contain enriched binding after treatment, compared to DMSO.

- *Metagene analysis*: To visualize differential eCLIP densities at near-nucleotide resolution, we used Metadensity (Her et al., 2022) to calculate relative information content (RI). RI values here represents differential binding of WX-02-23/WX-02-43/PladB compared to DMSO as the background. Specifically, for each nucleotide position in a transcript, 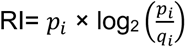, where *p_i_* and *q_i_* are the fraction of total reads in IP from either WX-02-23/WX-02-43/PladB and DMSO-treated IP libraries respectively that map to nucleotide i. We averaged the relative information over all transcripts with at least one differential enriched window detected by skipper (see section “Differential binding analysis”.

### QUANTIFICATION AND STATISTICAL ANALYSIS

Unless otherwise stated, quantitative data are expressed in bar and line graphs as mean ± SD (error bar) shown. SEC elution profiles are expressed as mean ± SEM (error bar). Unless otherwise stated, differences between two groups were examined using an unpaired two-tailed Student’s t-test with equal or unequal variance as noted. Significant P values are indicated (\**P*<0.05, \*\**P*<0.01, \*\*\**P*<0.001, and \*\*\*\**P*<0.0001).

## SUPPLEMENTAL INFORMATION

**Methods S1.** Synthetic Procedures and Characterization of Compounds

**Dataset S1.** Proteomic and RNA-seq data and Reference Tables, Related to Figures 1, 2, 3, 4, 5, 6, and 7.

**Dataset S2.** eCLIP data, Related to Figure 7

## Supporting information

Chemistry Methods

## ACKNOWLEDGEMENTS

We thank M. Hayward, S. Niessen, H. Murrey, B. Martin, R. Park, and A. Cognetta for helpful discussions, and B. Chen and X. Lieu (WuXi AppTec) for assistance with compound synthesis. This work was supported by the NIH (CA231991), American Cancer Society (PF-18-217-01-CDD), Vividion Therapeutics, Janssen Pharmaceuticals, and Pfizer. M. G. J. is supported by an EMBO long-term fellowship (EMBO ALTF 255-2021). G.W.Y. is supported by NIH R01 HG004659, U24 HG009889 and an Allen Distinguished Investigator Award, a Paul G. Allen Frontiers Group advised grant of the Paul G. Allen Foundation.

---

**Figure S1.**
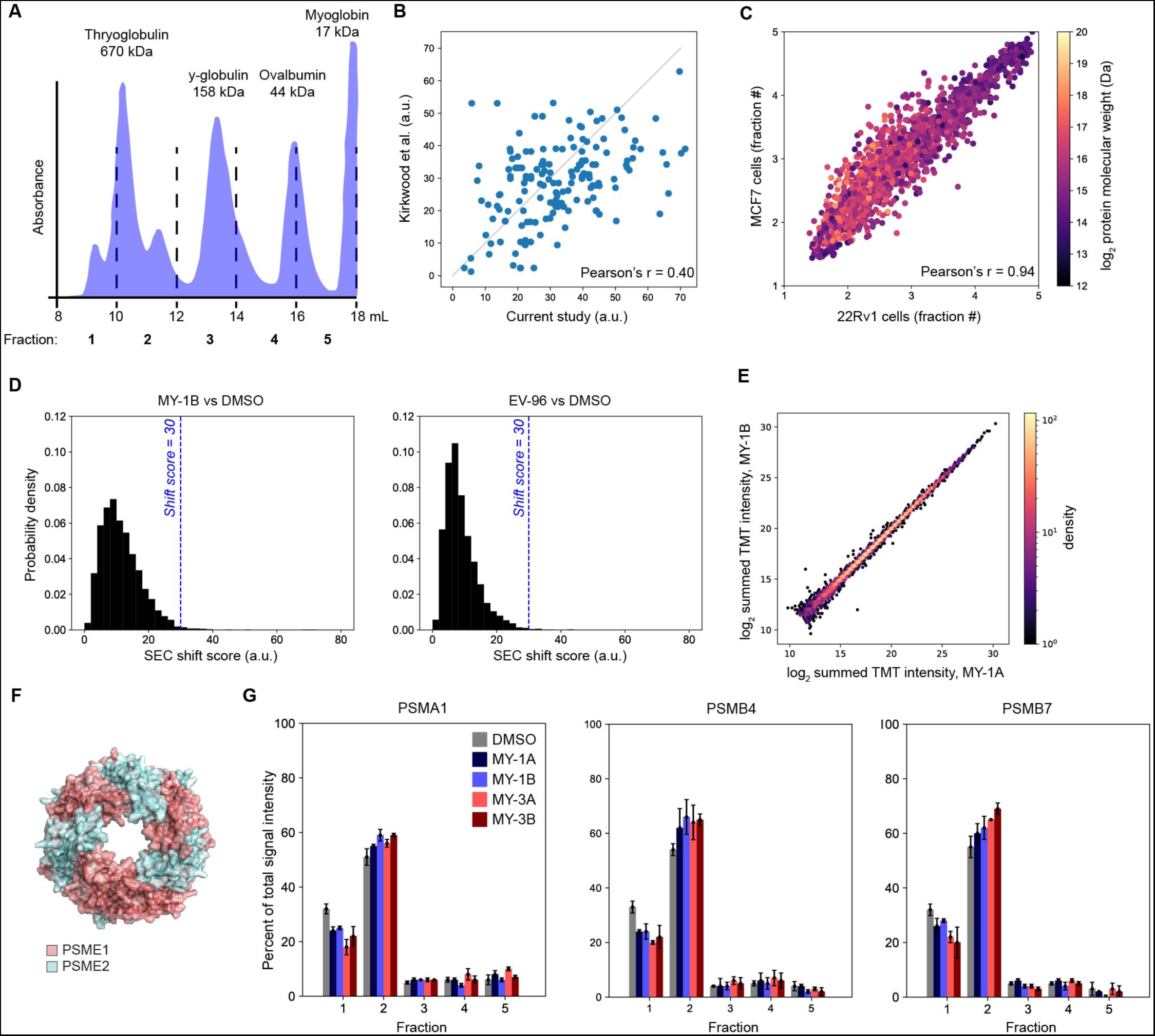
A proteomic platform to discover small-molecule modulators of protein-protein interactions in human cells, related to Figure 1. A. Distribution of proteins of indicated MWs as fractionated by SEC-MS method. B. Co-elution scores for protein complexes in the CORUM Core Complex dataset for both SEC-MS with 5 fractions (x-axis) and the data from Kirkwood *et. al.* with 40 fractions (y-axis). Each dot represents a single annotated protein complex – lower score represents tighter co-elution of complex members. Data are mean pairwise Euclidean distance across constituent proteins in each annotated protein complex from n = 3-11 independent experiments (current study) and n = 3 independent experiments from Kirkwood *et al*. C. Comparison of the weighted average elution times in SEC-MS experiments for proteins from 22Rv1 (x-axis) for MCF7 (y-axis) cells under control (DMSO) conditions. Each dot represents a single protein detected in both cell lines, and proteins are colored based on their annotated MW (UniProt). Large proteins elute in higher-MW fractions, while small proteins elute across all fractions, indicating the capture of native protein complexes. Data are mean elution times (weight average) of n = 2-11 and n = 2-6 independent experiments for 22Rv1 and MCF7 cells, respectively with Pearson’s r indicated. D. Distribution of SEC shift scores (a.u.) for proteins detected in MY-1B or EV-96 treated samples (20 µM, 3 h). The vertical dashed line marks a size shift value of 30, which was used as a minimal shift magnitude to designate potential compound-induced changes of interest. Data are shift scores calculated from average protein elution profiles from n = 2-11 independent experiments. E. Comparison of protein abundances in MY-1A vs MY-1B-treated 22Rv1 cells. Data are log_2_ transformed summed TMT intensities for proteins quantified from cells treated with MY-1A (TMT channels 1 – 5) or MY-1B (channels 6 – 10). Data for a total of 6395 unique quantified proteins is shown. Individual values from n = 2 independent experiments are shown. F. Structure of PA28 complex, showing heteroheptameric arrangement of PSME1 and PSME2 subunits (PDB 7DRW). G. SEC elution profiles for representative 20S core proteasome subunits, revealing that these proteins are not size-shifted by MY-1B or other stereoisomeric azetidine acrylamides (20 µM, 3 h). Data are average values ± SEM of the fractional distribution of protein reporter ion intensities from n = 2-11 independent experiments.

**Figure S2.**
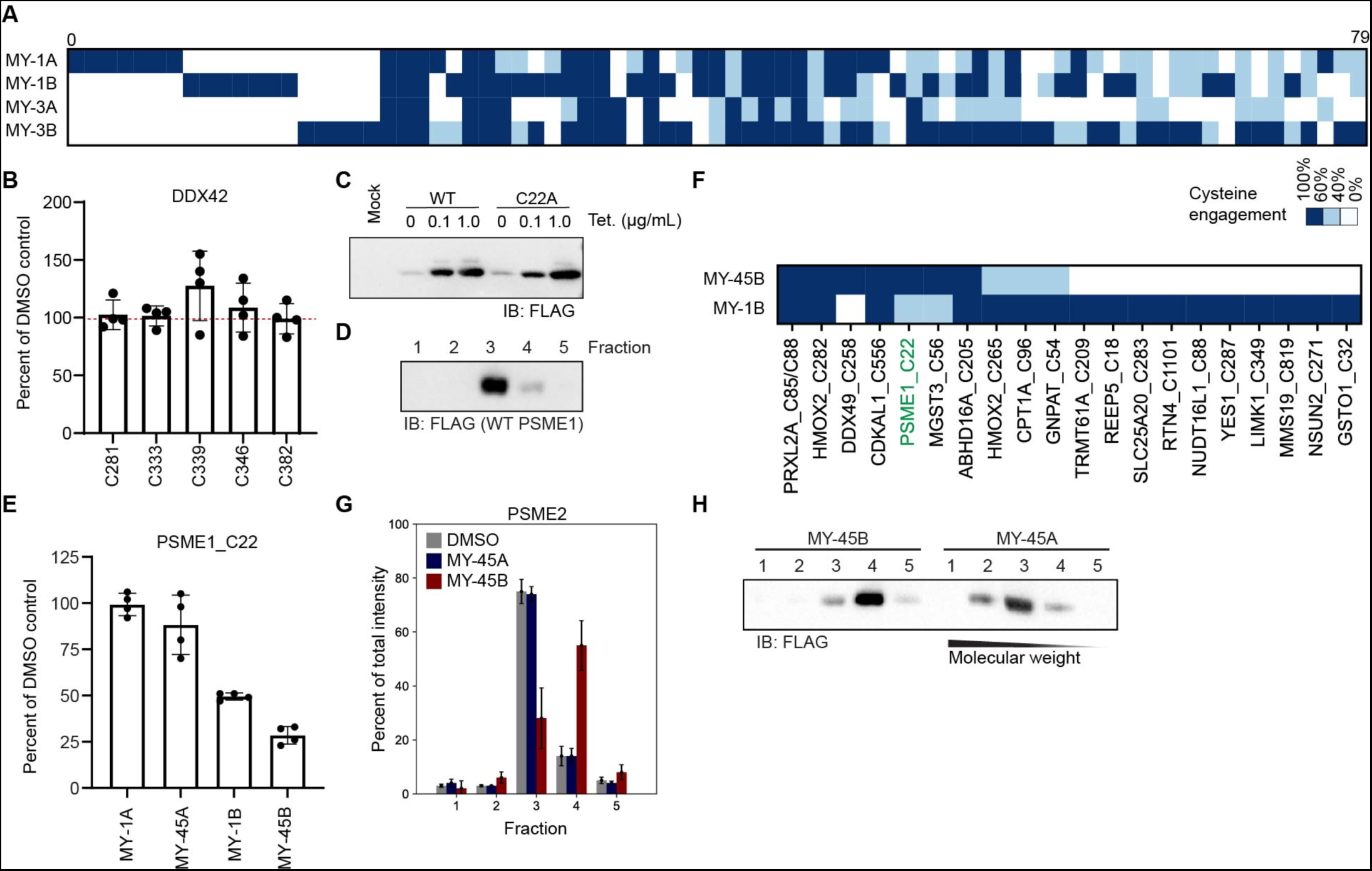
Electrophilic compounds stereoselectively disrupt the structure of the PA28 complex by engaging C22 of PSME1, Related to Figures 2 and 3. A. Heatmap of cysteines liganded (> 60% engagement) by azetidine acrylamides in 22Rv1 cells (20 µM compound, 1 h). Data are average values from n = 4 independent experiments. B. Lack of engagement of cysteines in DDX42 by EV-96 as measured by cysteine reactivity profiling experiments performed in 22Rv1 cells (20 µM EV-96, 1 h). Data are average values ± SD relative to DMSO from n = 4 independent experiments. C. Western blot of 22Rv1 TetR cells expressing recombinant FLAG epitope-tagged WT- or C22A-PSME1 (10 cm dish, 1 µg plasmid, 48 h transfection) in response to treatment with the indicated amounts of tetracycline. 0.1 µg/mL of tetracycline was selected as the concentration for inducing expression of PSME1 protein in 22Rv1 TetR cells. D. SEC elution profile as determined by western blotting for recombinant WT-PSME1 in 22Rv1 TetR cells. Data are from a single experiment representative of two independent experiments. E. Quantification of PSME1_C22 engagement by the indicated compounds as measured by cysteine-directed ABPP performed in 22Rv1 cells (5 µM compound, 3 h). Data are average values ± SD relative to DMSO from n = 4 independent experiments. F. Heatmap of cysteines liganded by MY-1B or MY-45B (5 µM, 3 h) in 22Rv1 cells as determined by cysteine-directed ABPP. A cysteine was considered liganded if it showed a > 60% reduction in IA-DTB labeling in compound-treated cells. Data are average values from n = 4 independent experiments. G. SEC-MS elution profile for endogenous PSME2 in 22Rv1 cells treated with DMSO, MY-45A, or MY-45B (20 μM, 3 h). Data are average values ± SEM of the fractional distribution of reporter ion intensities from n = 2-11 independent experiments. H. SEC elution profile as determined by western blotting recombinant WT-PSME1 in 22Rv1 TetR cells treated with MY-45A or MY-45B (5 µM, 3 h). Data are from a single experiment representative of 2 independent experiments.

**Figure S3.**
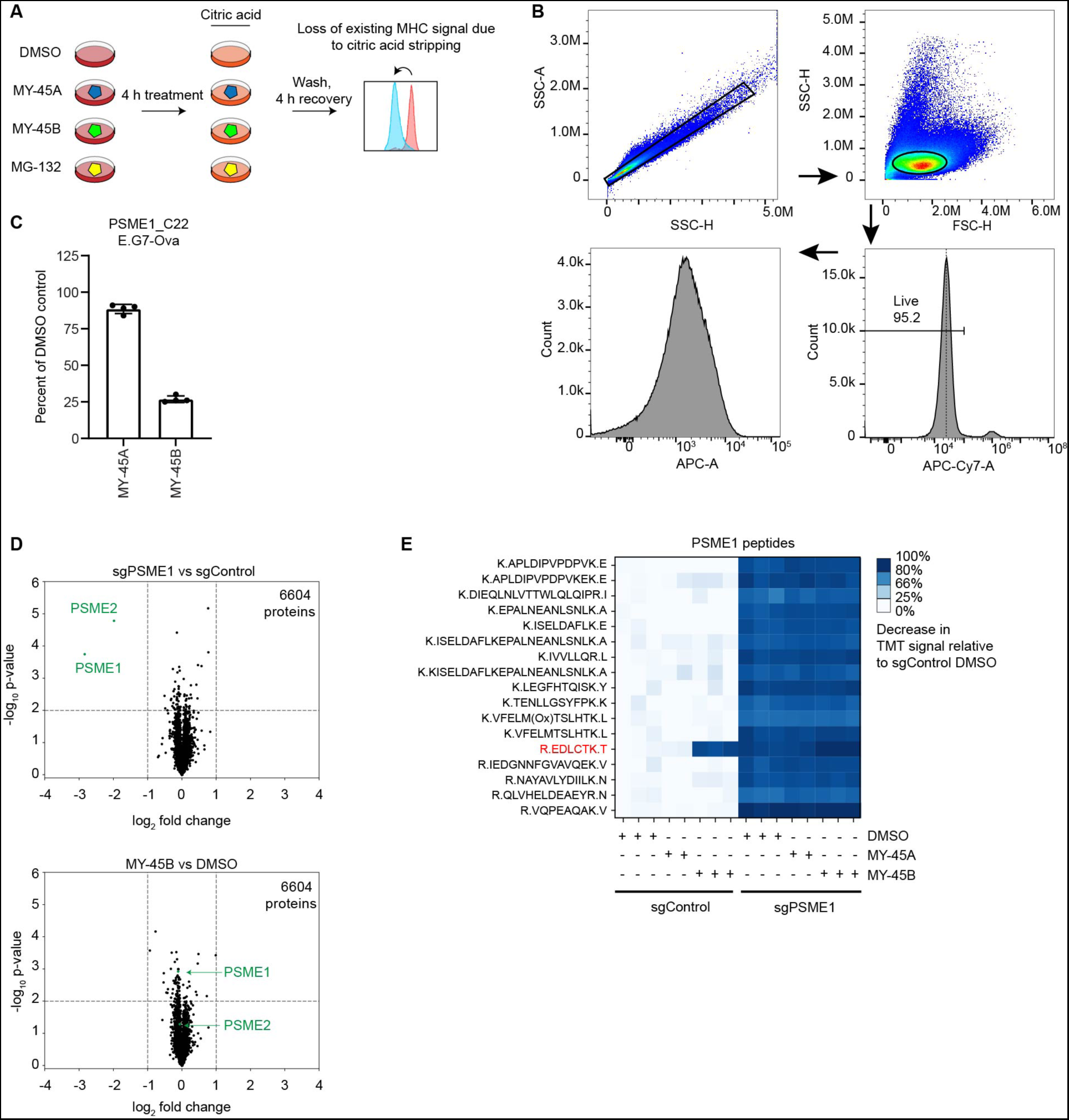
Chemical or genetic perturbation of PSME1 modulates MHC-I-immunopeptide interactions, related to Figures 3 and 4. A. Schematic of acid wash experiment to measure recovery of MHC-I antigen presentation: Cells were treated with compound for 4 h, washed with citric acid for 2 min, then allowed to recover for up to 4 h prior to analysis by flow cytometry to measure surface presentation of SIINFEKL antigen and MHC-I. B. Representative flow cytometry gating strategy for SIINFEKL (APC) signal quantification. C. Quantification of PSME1_C22 engagement measured by cysteine reactivity profiling experiments performed in E.G7-Ova cells (5 µM compound, 3 h). Data are average values ± SD relative to DMSO from n = 4 independent experiments. D. Volcano plots showing substantially (> two-fold increase or decrease) and significantly (*p* value < 0.01) changing proteins in MS-based proteomic experiments of sgPSME1 vs sgControl (sgAAVS1) cells (left) or sgControl cells treated with DMSO vs MY-45B (10 μM, 8 h). Data are average values from n = 3 independent experiments. E. Quantification of tryptic peptides from PSME1 in MS-based proteomic experiments of sgControl and sgPSME1 cells treated with the indicated compounds (10 µM, 8 h). Note that the tryptic peptide containing PSME1_is near-completely absent in cells treated with MY-45B, but unchanged in cells treated with MY-45A or DMSO. Data are individual values relative to DMSO (n = 2-3 per group).

**Figure S4.**
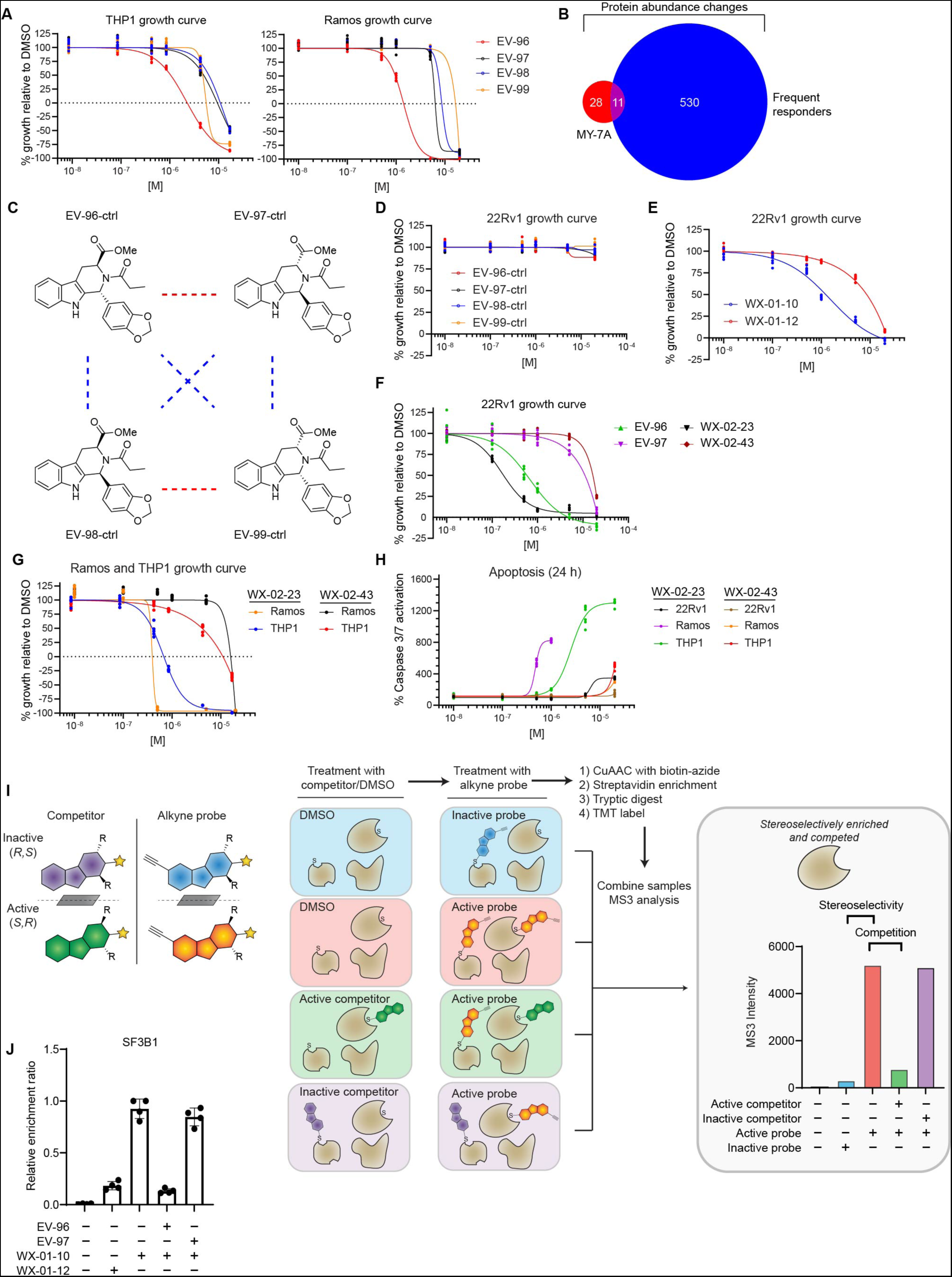
Tryptoline acrylamides that stereoselectively engage SF3B1 alter protein abundances and block the proliferation of cancer cells, related to Figure 5. A. EV-96 stereoselectively blocks the proliferation of THP1 and Ramos cells. Cells were treated with the indicated concentrations of tryptoline acrylamides for 72 h, after which cell growth was measured by CellTiter-Glo. Data are relative to DMSO control from n = 6 independent experiments. B. Venn diagram of proteins showing > 33% decreases in abundance in 22Rv1 cells treated with EV-96 (20 µM, 8 h; see Figure 6A) and proteins annotated as “frequent responders” by Rupert *et al*. Only proteins quantified in both proteomic studies were included in the Venn diagram. C. Structures of non-electrophilic analogues of tryptoline acrylamides EV-96-ctrl (**17**), EV-97-ctrl (**18**), EV-98-ctrl (**19**), and EV-99-ctrl (**20**). D. Non-electrophilic analogues do not affect the proliferation of 22Rv1 cells. Cells were treated with the indicated concentrations of compounds for 72 h, after which cell growth was measured by CellTiter-Glo. Data are relative to DMSO control from n = 6 independent experiments. E. Alkyne probe WX-01-10 stereoselectively blocks the proliferation of 22Rv1 cells. Cells treated with the indicated concentrations of WX-01-10 or WX-01-12 for 72 h, after which cell growth was measured by CellTiter-Glo, Data are average values ± SD relative to DMSO from n = 6 independent experiments. F. Enhanced anti-proliferative activity of WX-02-23 compared to EV-96. 22Rv1 cells were treated with the indicated concentrations of tryptoline acrylamides for 72 h, after which cell growth was measured by CellTiter-Glo. Data are relative to DMSO control from n = 6 independent experiments. G. Anti-proliferative activity of WX-02-23 in THP1 and Ramos cells. Cells were treated with the indicated concentrations of tryptoline acrylamides for 72 h, after which cell growth was measured by CellTiter-Glo. Data are relative to DMSO control from n = 6 independent experiments. H. Apoptotic activity of WX-02-23 in human cancer cell lines. The indicated cell lines were treated with the indicated concentrations of tryptoline acrylamides for 24 h, after which apoptosis was measured by Caspase-Glo 3/7. Data are relative to DMSO control from n = 6 independent experiments. I. Schematic for target identification strategy using alkyne probe enrichment and blockade of this enrichment by competitor compounds. 22Rv1 cells were treated with DMSO or active and inactive enantiomeric tryptoline acrylamides (“competitors”; 5 µM, 2 h), followed by DMSO or active and inactive enantiomeric alkyne probes (10 µM, 1 h). Cells were then harvested, lysed, and probe-labeled proteins biotinylated by CuAAC with biotin-azide, enriched with streptavidin, and analyzed by quantitative chemical proteomics. On the right, is a mock quantification profile for a candidate target responsible for stereoselective anti-proliferative activity, where such a target would show stereoselective enrichment by active alkyne probe and stereoselective blockade of this enrichment by active competitor. J. Quantification of stereoselective enrichment and competition of SF3B1 by active alkyne probe (WX-01-10) and competitor (EV-96) vs inactive alkyne enantiomer probe (WX-01-12) and inactive enantiomer competitor (WX-02-43). Data are average values ± SD normalized to WX-01-10 treatment group for n = 4 independent experiments.

**Figure S5.**
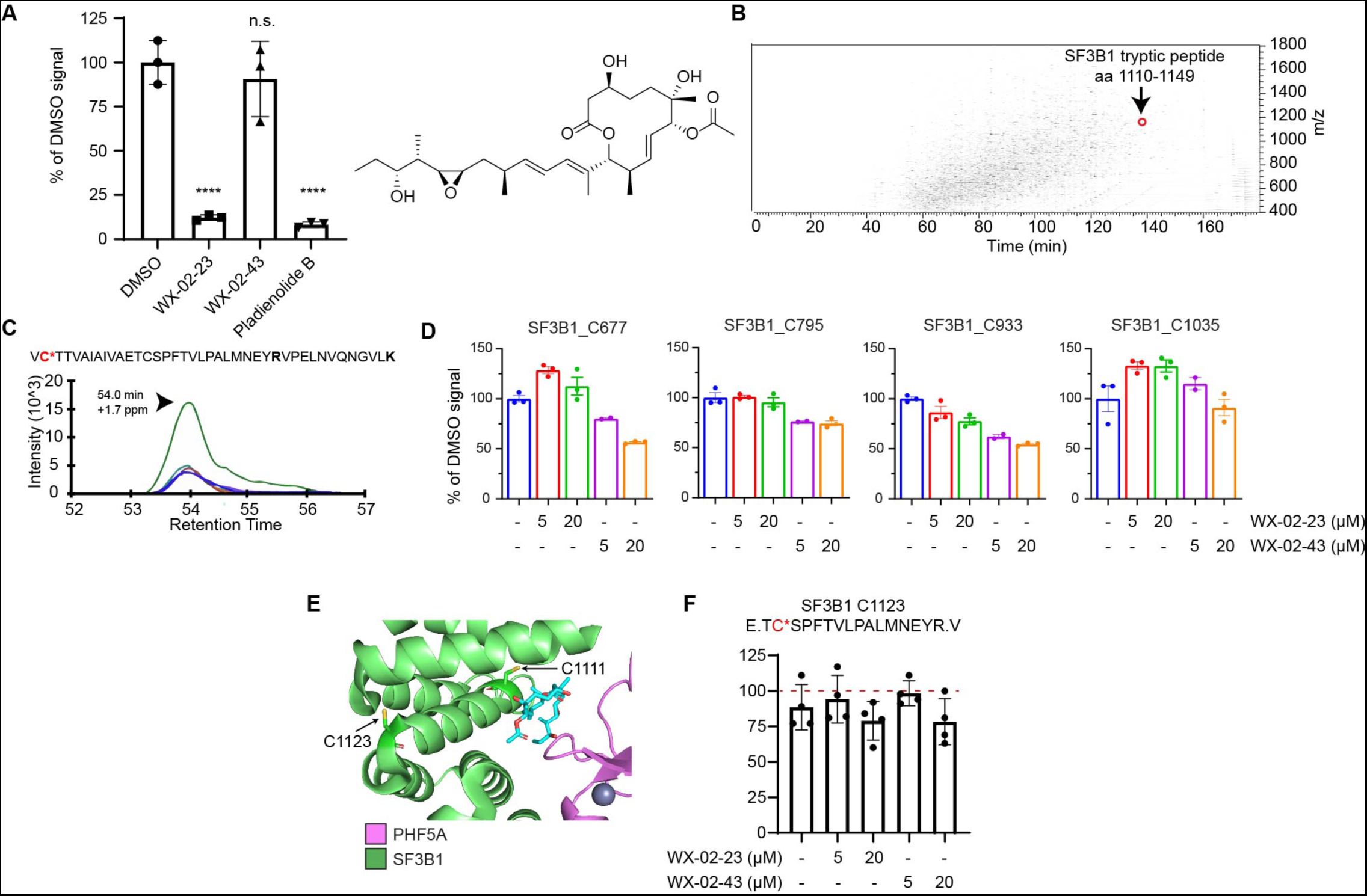
Tryptoline acrylamides engage C1111 of SF3B1, related to Figure 6. A. Quantification of fluorescent gel data showing WX-01-10 labelling of a 150 kDa protein consistent with SF3B1, as shown in Figure 6A. Data are average values ± SD relative to DMSO from n = 3 independent experiments. ****, p<0.0001, Significance determined by one-way ANOVA with Dunnett’s multiple comparisons test. B. LC chromatogram of IA-DTB-labeled SF3B1 peptide (aa 1110-1149) eluting late in the gradient at ∼38% acetonitrile. C. Representative chromatogram elution profile of fragment ions for SF3B1 peptide aa 1110-1149 harboring C1111 (labeled in red). D. Quantification of additional SF3B1 cysteines by targeted cysteine reactivity profiling, revealing that none of these residues are engaged by WX-02-23 (5 or 20 µM, 3 h) in 22Rv1 cells. Data are average values ± SD relative to DMSO from n = 2-3 independent experiments. E. Crystal structure of SF3B1-PHF5A complex bound to pladienolide B, highlighting the locations of C1111 and C1123 (PDB: 6EN4). F. Quantification of SF3B1_C1123 reactivity following trypsin and GluC digestion of proteomes from 22Rv1 cells treated with WX-02-23 or WX-02-43 (5 or 20 µM, 3 h). Data are average values ± SD relative to DMSO from n = 4 independent experiments.

**Figure S6.**
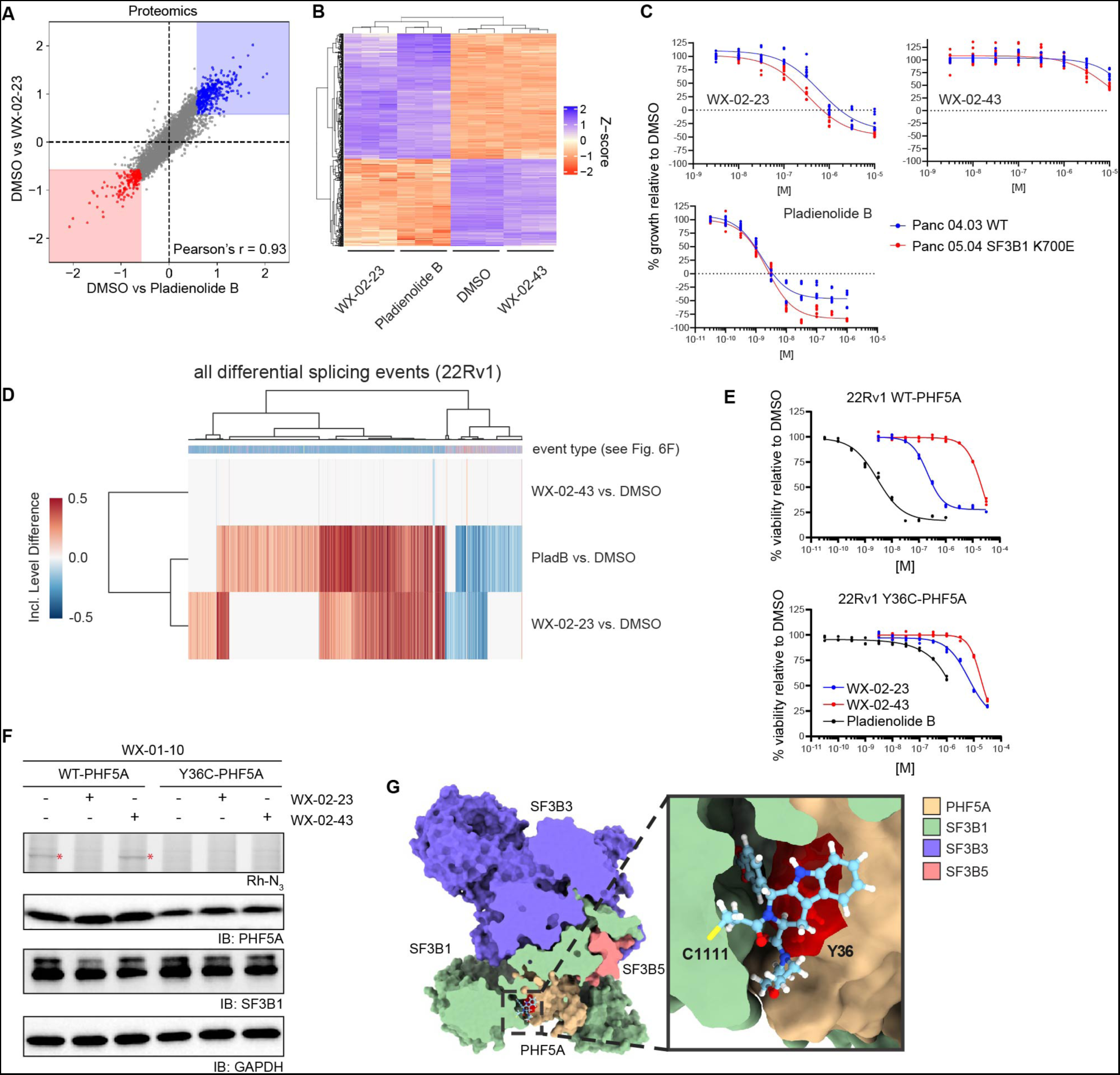
Tryptoline acrylamides engaging SF3B1 stereoselectively modulate spliceosome function, related to Figure 6. A. Scatter plot of protein abundance changes in 22Rv1 cells treated with WX-02-23 (1 µM) pladienolide B (10 nM), or DMSO for 24 h. Proteomics data are average values shown as log_2_ fold change relative to DMSO for n = 4 independent experiments. B. Hierarchical clustering of RNA-seq data showing similar changes in 22Rv1 cells treated with pladienolide B (10 nM, 8 h) or WX-02-23 (5 µM, 8 h) compared to WX-02-43 (5 µM 8 h) or DMSO-treated cells. Data are average values for n = 3 independent experiments. C. Cell growth effects of SF3B1 ligands WX-02-23 and pladienolide B (and control compound WX-02-43) on Panc 04.03 and Panc 05.04 cells, which express WT-SF3B1 and the cancer-related K700E-SF3B1 mutant, respectively. Cells were treated with the indicated concentrations of compounds for 72 h, after which cell growth was measured by CellTiter-Glo. Data are relative to DMSO control from n = 9 independent experiments. D. Clustered heatmap of inclusion level differences between 22Rv1 cells treated with the indicated compounds vs DMSO for all alternative splicing events from Figure 6F. Columns are annotated by the type of alternative splicing event in the color scheme of Figure 6F. E. Cell growth effects of WX-02-23 and pladienolide B in 22Rv1 cells recombinantly expressing WT-PHF5A (upper plot) or a Y36C-PHF5A mutant (lower plot). Cells were treated with the indicated concentrations of compounds for 72 h, after which cell growth was measured by CellTiter-Glo. Data are relative to DMSO control from n = 3 independent experiments. F. Assessment of reactivity of the WX-01-10 alkyne probe with SF3B1 in 22Rv1 cells expressing WT-PHF5A or Y36C-PHF5A by gel-ABPP. Top panel, gel-ABPP showing proteins labeled by WX-01-10 alkyne probe in the indicated cells. Cells were pre-treated with DMSO, WX-02-23 (1 µM) or WX-02-43 (1 µM) for 2 h followed by treatment with WX-01-10 (1 µM, 1 h), lysis, CuAAC with rhodamine azide, SDS-PAGE, and in-gel fluorescence scanning. Red asterisk marks a 150 kDa protein interpreted to be SF3B1 that is labeled by WX-01-10 and blocked in labeling by WX-02-23, but not inactive enantiomer WX-02-43. The labeling of SF3B1 by WX-01-10 is blocked in cells expressing the Y36C-PHF5A cells. Remaining panels represent western blots as loading controls. Results are from a single experiment representative of two independent experiments. G. Y36 of PHF5A is located in the predicted WX-02-23 binding cavity at the interface with SF3B1. Computational predictions of small-molecule binding to SF3B1 were performed on Schrodinger Maestro (V 12.3.012, MMshare V 4.9.012, Release 2020-1, Platform Windows-x64) using a structure of pladienolide B bound to SF3B1 (PDB: 6EN4). Covalent docking was performed using Glide. Images were generated using UCSF ChimeraX.

**Figure S7.**
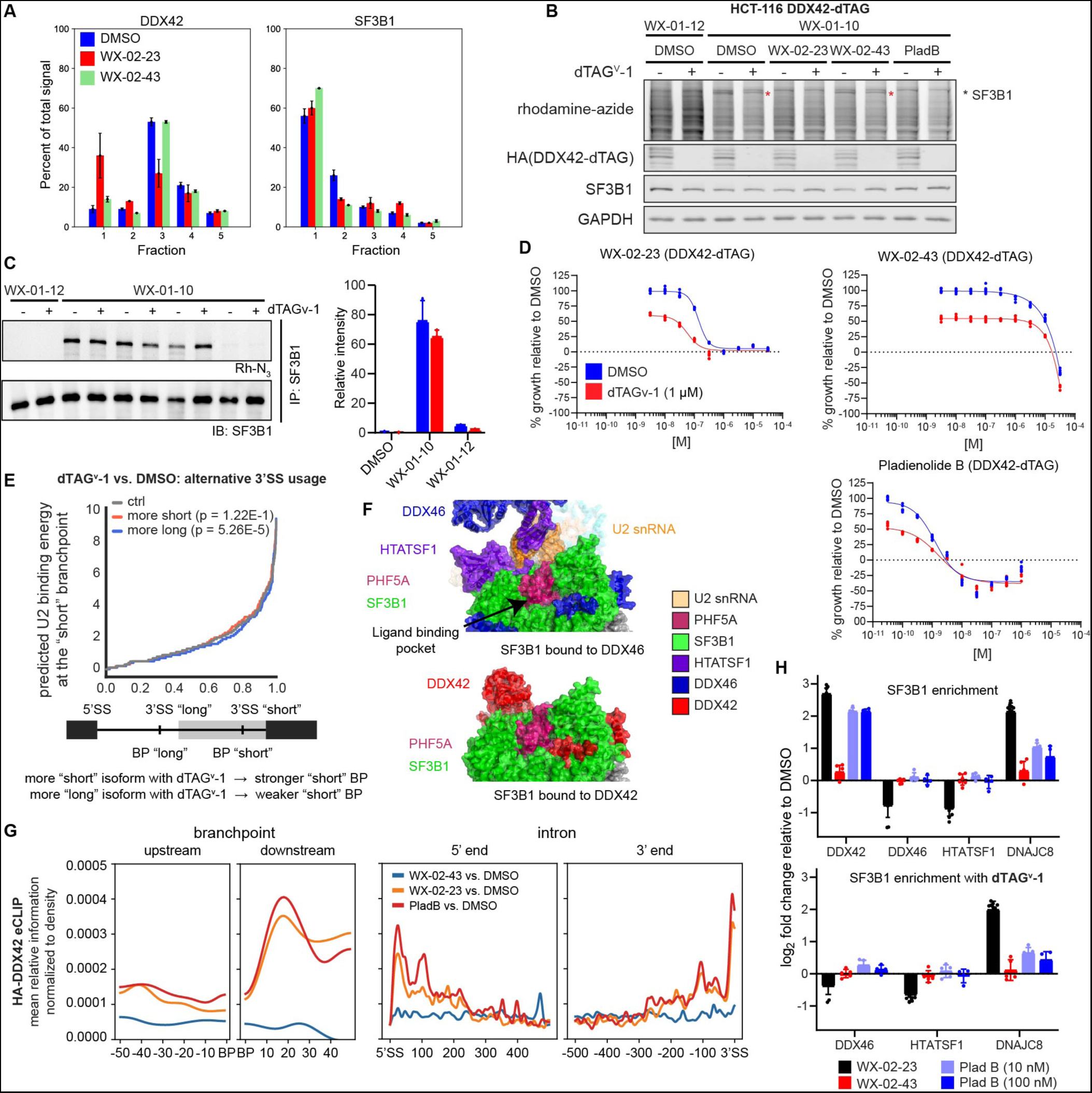
DDX42 facilitates branch point selection and associates more strongly with SF3B1 in WX-02-23 or pladienolide B-treated cells, related to Figure 7. A. SEC-MS elution profiles for DDX42 and SF3B1 in 22Rv1 cells treated with DMSO, WX-02-23, or WX-02-43 (20 μM, 3 h). Data are average values ± SEM of the fractional distribution of reporter ion intensities from n = 2-11 independent experiments. B. Impact of DDX42-dTAG pre-degradation on SF3B1 engagement by WX-01-10 alkyne probe as determined by gel-ABPP. DDX42-dTAG HCT-116 cells were pre-treated with dTAGv-1 (500 nM) or DMSO for 1 h and then treated with the DMSO, WX-02-23 (1 µM), WX-02-43 (1 µM), or pladienolide B (10 nM) for 2 h, followed by treatment with WX-01-10 or WX-01-12 (1 μM, 1 h) and processing for gel-ABPP as described in **Figure S6F**. Asterisk labels the 150 kDa band interpreted as SF3B1. Results are from a single experiment representative of two independent experiments. C. Impact of DDX42-dTAG pre-degradation on SF3B1 engagement by WX-01-10 alkyne probe as determined by gel-ABPP following immunoprecipitation of SF3B1. DDX42-dTAG HCT-116 cells were pre-treated with dTAGv-1 (500 nM) or DMSO for 1 h and then treated with WX-01-10 or WX-01-12 (2.5 µM) for 1 h. Immunoprecipitation was performed with an anti-SF3B1 antibody (CST #14434) followed by CuAAC of immunprecipitated proteins to an azide-rhodamine tag and analysis by SDS-PAGE and in-gel fluorescence scanning (top image) or western blotting (bottom image). Quantification normalized to SF3B1 enrichment from gel (left) is shown (right). D. Growth effects of WX-02-23 or pladienolide B in cells lacking DDX42. DDX42-dTAG HCT-116 cells were pre-treated with dTAGv-1 (1 µM) or DMSO for 1 h and then treated with the indicated concentrations of WX-02-23, pladienolide B (PladB), or the control compound WX-02-43 for 72 h, after which cell growth was measured by CellTiter-Glo. Data are relative to DMSO control from n = 6 independent experiments. E. DDX42 loss increased the usage of stronger branch point sequences and decreased the usage of weaker branch points at differentially included exons BP usage. DDX42-dTAG HCT-116 cells were pre-treated for 1 h with either 500 nM dTAGv-1 or DMSO, followed by addition of either DMSO or compounds to the same supernatant for 8 h. Data are from the union of n = 3 biological replicates. F. Cryo-EM structures of SF3B1 bound to DDX46/HTATSF1 (PDB: 7EVO) or DDX42 (PDB: 7EVN). Structures were overlayed and the same perspective is shown demonstrating the shared interaction sites of DDX42 and DDX46/HTATSF1 with SF3B1. G. Effects of SF3B1 ligands WX-02-23 and pladienolide B (PladB) on metadensity plots of HA-DDX42 eCLIP signal (relative information of IP of indicated treatment over IP of DMSO). DDX42-dTAG HCT-116 cells were treated with WX-02-23 (5 µM), WX-02-43 (5 µM), or pladienolide B (100 nM) for 3 h after which DDX42 eCLIP experiments were performed using the HA-tag of the dTAG fusion. Data are average values from the union of n = 2 independent experiments. H. Quantification of proteins showing differential co-immunoprecipitation with SF3B1 in cells treated with WX-02-23 or pladienolide B (PladB) and the effect of DDX42 disruption on these interactions. Cells were pre-treated with dTAGv-1 (500 nM) or DMSO for 1 h and then treated with WX-02-23 (5 µM), WX-02-43 (5 µM), pladienolide B (100 nM), or DMSO for 3 h. Co-immunoprecipitation analysis was performed with anti-SF3B1 antibody (CST #14434) followed by MS-based proteomics. Data are average values ± SD relative to DMSO from n = 4-10 independent experiments.

