## Supplementary material for "Proteomic discovery of chemical probes that perturb protein complexes in human cells": Chemistry Methods

### Methods S1

#### General considerations

All reagents and solvents were purchased and used as received from commercial vendors or synthesized according to cited procedures. Yields refer to chromatographically and spectroscopically pure compounds, unless otherwise stated. Flash chromatography was performed using 20-40 μm silica gel (60-Å mesh) on a Teledyne Isco Combiflash Rf or a Biotage Isolera Prime, alternatively in a glass column using SiliaFlash® F60 40-63 μm silica gel (60-Å mesh). Preparative high-pressure liquid chromatography (prep-HPLC) was performed on a Gilson GX-281 instrument equipped with a Phenomenex Gemini C18 column (150 mm × 25 mm × 10 μM) eluting with a mixture of acetonitrile and a buffered aqueous phase. Aqueous buffers are denoted as follows: BASE (0.05% ammonia v/v), TFA (0.075% trifluoroacetic acid v/v), FA (0.225% formic acid v/v), HCL (0.05% concentrated hydrochloric acid, v/v), NEU (10 mmol ammonium bicarbonate). Analytical thin layer chromatography (TLC) was performed on 0.2 mm or 0.25 mm silica gel 60-F plates and visualized by UV light (254 nm). Preparative thin layer chromatography (prep-TLC) was performed on GF254 plates (acrylic adhesive, 0.5×200×200 mm, 5–20 μM particle size, 250 μM thickness). NMR spectra were recorded on Bruker Avance III 400, Avance III HD 400, Avance Neo 400 spectrometers ($^1$H, 400 MHz) at 300 K unless otherwise noted. Data for $^1$H NMR are reported as follows: chemical shift (δ), multiplicity (s = singlet, d = doublet, t = triplet, m = multiplet; br = broad), coupling constants, and integration. Chemical shifts are reported in parts per million (ppm) using the appropriate solvent as reference. Analytical supercritical fluid chromatography (SFC) was performed on a Shimadzu LC system (flow rate: 3 mL/min, back pressure: 100 Bar, column temperature: 35 ºC) equipped with a polydiode array detector unless otherwise noted. Tandem liquid chromatography/mass spectrometry (LC-MS) was performed on an Agilent 1200 series LC/MSD system equipped with an Agilent G6110A mass detector, alternatively a Shimadzu LC-20AD or AB series LC-MS system equipped with Shimadzu SPD-M20A or SPDM40 mass detectors, alternatively a Waters H-Class LC with equipped with diode array and QDa mass detector.

#### Synthesis of azetidine probes

##### Synthesis of MY-1A

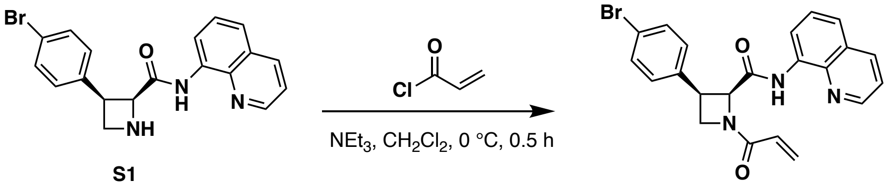

**S1** **MY-1A** (**1**)

##### (2*S*,3*R*)-1-acryloyl-3-(4-bromophenyl)-*N*-(quinolin-8-yl)azetidine-2-carboxamide (MY-1A) (1)

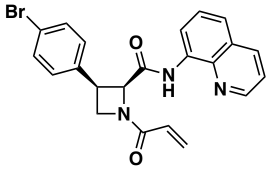

To a precooled (0 ºC) solution of **S1** (50.0 mg, 131 μmol) (Maetani et al., 2017) in dichloromethane (1 mL) were added triethylamine (26.5 mg, 262 μmol) and acryloyl chloride (14.2 mg, 157 μmol). The mixture was stirred at 0 °C for 0.5 hours. Upon completion, the reaction mixture was concentrated under reduced pressure to obtain a residue, which was purified by prep-TLC ($SiO_2$, petroleum ether/EtOAc = 2:1) to give **MY-1A** (51.0 mg, 89% yield) as a white solid.

**HRMS ESI-TOF** m/z calculated for $C_{22}H_{19}BrN_3O_2$ $[M+H]^+$ 436.0655. Found 436.0646.

**$^1$H NMR** (400 MHz, $CDCl_3$): δ 10.47 (br s, 1H), 8.81 (d, *J* = 3.6 Hz, 1H), 8.43 (d, *J* = 4.0 Hz, 1H), 8.16 (d, *J* = 8.0 Hz, 1H), 7.54-7.42 (m, 3H), 7.29-7.25 (m, 4H + $CHCl_3$), 6.53 (d, *J* = 16.3 Hz, 1H), 6.48-6.18 (m, 1H), 5.98-5.62 (m, 1H), 5.38 (d, *J* = 8.0 Hz, 1H), 4.64 (t, *J* = 9.3 Hz, 1H), 4.59-4.40 (m, 1H), 4.33-4.24 (m, 1H).

##### Synthesis of MY-1B

Prepared in analogous fashion from *ent*-**S1** (Maetani et al., 2017).

##### (2*R*,3*S*)-1-acryloyl-3-(4-bromophenyl)-*N*-(quinolin-8-yl)azetidine-2-carboxamide (MY-1B) (2)

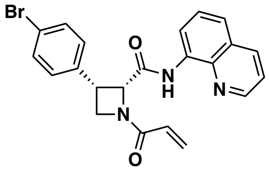

**HRMS ESI-TOF** m/z calculated for $C_{22}H_{19}BrN_3O_2$ $[M+H]^+$ 436.0655. Found 436.0649.

**$^1$H NMR** (400 MHz, $CDCl_3$): δ 10.47 (s, 1H), 8.80 (d, *J* = 4.0 Hz, 1H), 8.42 (d, *J* = 4.0 Hz, 1H), 8.15 (d, *J* = 8.0 Hz, 1H), 7.52-7.40 (m, 3H), 7.30-7.25 (m, 4H + $CHCl_3$), 6.64-6.15 (m, 2H), 5.90-5.64 (m, 1H), 5.37 (d, *J* = 8.0 Hz, 1H), 4.63 (t, *J* = 9.3 Hz, 1H), 4.57-4.38 (m, 1H), 4.33-4.22 (m, 1H).

#### Synthesis of MY-3A

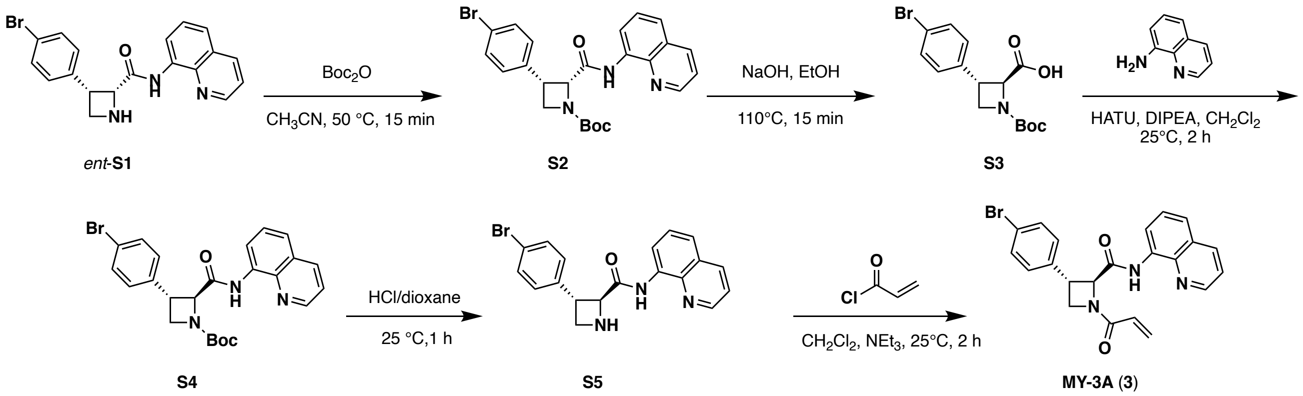

##### (2*S*,3*S*)-3-(4-bromophenyl)-1-(*tert*-butoxycarbonyl)azetidine-2-carboxylic acid (S3)

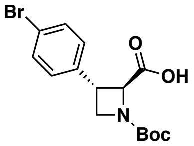

To a solution of *ent*-**S1** (450 mg, 1.18 mmol) in acetonitrile (3 mL) was added $Boc_2O$ (308 mg, 1.41 mmol), and the resulting mixture was stirred at 50 °C for 15 min. Upon completion, the reaction mixture was filtered over Celite and concentrated under reduced pressure to give **S2** (600 mg, crude) as a yellow oil. To a solution of **S2** (450 mg, 933 µmol) in ethanol (3 mL) was added sodium hydroxide (373 mg, 9.33 mmol), and the resulting mixture was stirred at 110 °C for 15 min. Upon completion, the mixture was diluted with water and washed with dichloromethane (2×100 mL). The resulting aqueous solution was acidified with HCl (1 M) to adjust pH to 5~6, exhaustively extracted with i-PrOH/$CHCl_3$ (3:7, 5×60 mL), dried over anhydrous sodium sulfate, filtered, and concentrated under reduced pressure to give **S3** (330 mg, 99% yield over two steps) as a white solid.

**LC-MS** m/z calculated for $C_{15}H_{19}BrNO_4$ $[M+H]^+$ 356.0. Found 356.0.

##### (2*S*,3*S*)-3-(4-bromophenyl)-*N*-(quinolin-8-yl)azetidine-2-carboxamide (S5)

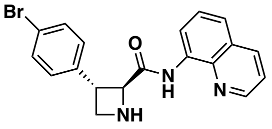

To a solution of **S3** (330 mg, 926 µmol) in dichloromethane (4 mL) were added diisopropylethylamine (239 mg, 1.85 mmol) and 8-aminoquinoline (23.7 mg, 262 µmol), followed by HATU (705 mg, 1.85 mmol). The resulting mixture was stirred at 25 °C for 2 hours. Upon completion, the reaction mixture was concentrated under reduced pressure to give a residue, which was purified by prep-TLC ($SiO_2$, petroleum ether/EtOAc = 2:1) to give **S4** (200 mg) as a yellow solid used directly in the next step. To a solution of **S4** (100 mg) in dichloromethane (1 mL) was added HCl/dioxane (4 M, 1 mL). and the resulting mixture was stirred at 25 °C for 1 hour. Upon completion, the reaction mixture was partitioned between ethyl acetate (40 mL) and brine (30 mL). The water layer was extracted with i-PrOH/$CHCl_3$ (3:7, 5×20 mL), then the organic phase was dried over sodium sulfate, filtered, and concentrated under reduced pressure to give **S5** (90.0 mg, 51% yield over two steps) as an off-white solid.

**LC-MS** m/z calculated for $C_{19}H_{17}BrN_3O$ $[M+H]^+$ 382.1. Found 382.1.

**(2*S*,3*S*)-1-acryloyl-3-(4-bromophenyl)-*N*-(quinolin-8-yl)azetidine-2-carboxamide (MY-3A) (3)**

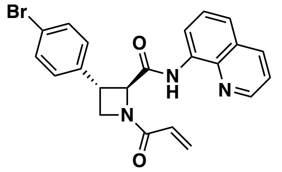

To a solution of **S5** (80.0 mg, 209 µmol) in dichloromethane (1 mL) were added triethylamine (42.4 mg, 419 µmol) and acryloyl chloride (37.9 mg, 419 µmol). The mixture was stirred at 25 °C for 2 hours. Upon completion, the reaction mixture was concentrated under reduced pressure. The residue was purified by preparative TLC ($SiO_2$, petroleum ether/EtOAc = 2:1) to give **MY-3A** (34.0 mg, 37% yield) as a white solid.

**HRMS ESI-TOF** m/z calculated for $C_{22}H_{19}BrN_3O_2$ $[M+H]^+$ 436.0655. Found 436.0649.

**$^1$H NMR** (400 MHz, $CDCl_3$): δ 11.13-10.67 (m, 1H), 8.95-8.84 (m, 1H), 8.83-8.75 (m, 1H), 8.16 (d, *J* = 8.0 Hz, 1H), 7.60-7.50 (m, 4H), 7.45 (dd, *J* = 8.2, 4.2 Hz, 1H), 7.33-7.28 (m, 2H), 6.54 (br d, *J* = 16.9 Hz, 1H), 6.42-6.25 (m, 1H), 5.91-5.75 (m, 1H), 5.18-4.95 (m, 1H), 4.76-4.55 (m, 1H), 4.43-4.14 (m, 2H).

**Synthesis of MY-3B**

Prepared in analogous fashion from **S1**.

**(2*S*,3*S*)-1-acryloyl-3-(4-bromophenyl)-*N*-(quinolin-8-yl)azetidine-2-carboxamide (MY-3B) (4)**

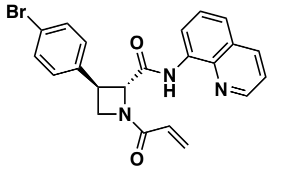

**HRMS ESI-TOF** m/z calculated for $C_{22}H_{19}BrN_3O_2$ $[M+H]^+$ 436.0655. Found 436.0643.

**$^1$H NMR** (400 MHz, $CDCl_3$): δ 10.95 (br s, 1H), 8.97-8.69 (m, 2H), 8.15 (d, *J* = 8.2 Hz, 1H), 7.55 (d, *J* = 4.7 Hz, 2H), 7.52 (d, *J* = 8.1 Hz, 2H), 7.45 (dd, *J* = 8.3, 4.2 Hz, 1H), 7.28 (d, *J* = 8.2 Hz, 2H), 6.53 (d, *J* = 16.9 Hz, 1H), 6.41-6.27 (m, 1H), 5.90-5.75 (m, 1H), 5.15-5.02 (m, 1H), 4.73-4.59 (m, 1H), 4.37-4.21 (m, 2H).

#### Synthesis of MY-11B

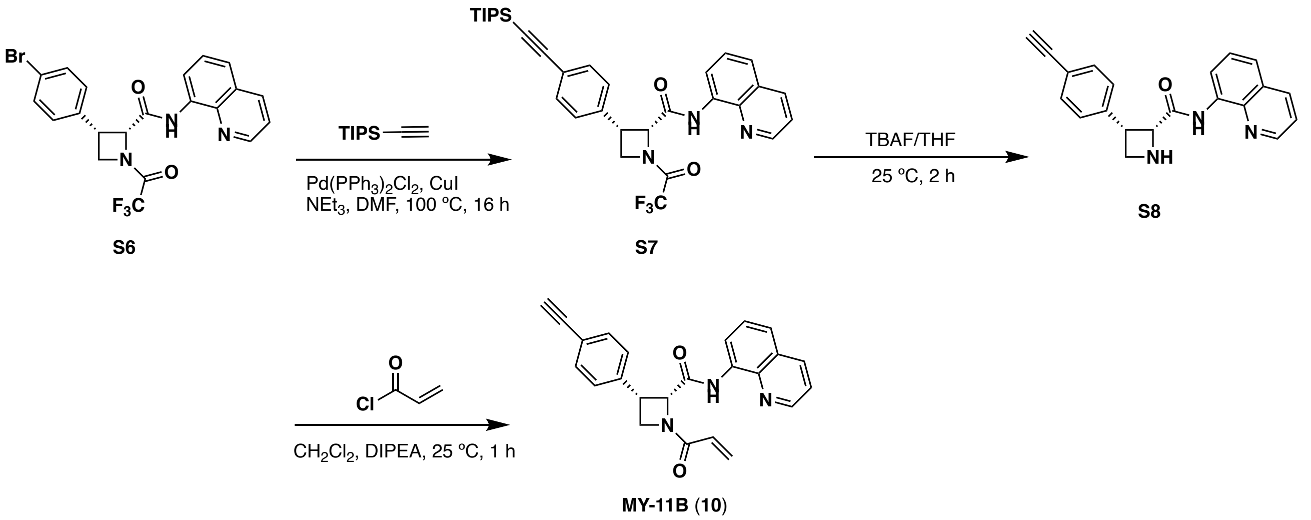

#### (2*R*,3*S*)-*N*-(quinolin-8-yl)-1-(2,2,2-trifluoroacetyl)-3-(4-((triisopropylsilyl)ethynyl)phenyl)azetidine-2-carboxamide (S7)

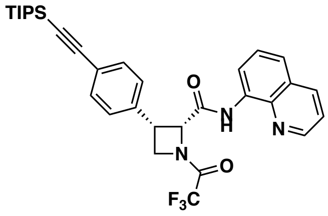

To a solution of **S6** (690 mg, 1.44 mmol) and (triisopropylsilyl)acetylene (789 mg, 4.33 mmol) in *N,N*-dimethyl formamide (2 mL) were added copper(I) iodide (27.5 mg, 144 μmol), $Pd(PPh_3)_2Cl_2$ (101 mg, 144 μmol) and triethylamine (292 mg, 2.89 mmol). The mixture was stirred at 100 °C for 16 hours under nitrogen atmosphere. Upon completion, the reaction mixture was partitioned between ethyl acetate (60 mL) and brine (40 mL). The water layer was extracted with ethyl acetate (40 mL×3). The organic layers were combined, dried over anhydrous sodium sulfate, filtered, and concentrated under reduced pressure to give a residue, which was purified by column chromatography ($SiO_2$, petroleum ether/EtOAc = 100:1 to 1:1) to give **S7** (500 mg, 59% yield) as a white solid.

**LC-MS** m/z calculated for $C_{32}H_{37}F_3N_3O_2Si$ $[M+H]^+$ 580.3. Found 580.4.

**$^1$H NMR** (400 MHz, $CD_3OD$, mixture of rotamers): δ 10.18-9.69 (m, 1H), 8.84-8.65 (m, 1H), 8.52-8.33 (m, 1H), 8.14 (dd, *J* = 8.3, 1.7 Hz, 1H), 7.53-7.39 (m, 3H), 7.34-7.22 (m, 3H + $CHCl_3$), 7.17 (d, *J* = 8.0 Hz, 1H), 5.66-5.38 (m, 1H), 4.91-4.39 (m, 3H), 1.11-0.91 (m, 21H).

**(2*R*,3*S*)-1-acryloyl-3-(4-ethynylphenyl)-*N*-(quinolin-8-yl)azetidine-2-carboxamide (MY-11B) (10)**

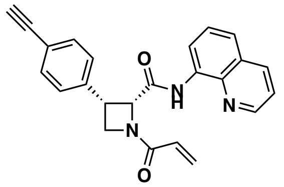

To a solution of **S7** (500 mg, 862 µmol) in tetrahydrofuran (8.6 mL) was added tetrabutylammonium fluoride (1 M in THF, 8.6 mL). The mixture was stirred at 25 °C for 2 hours. Upon completion, the reaction mixture was concentrated in vacuo to give the residue. The residue was purified by prep-TLC ($SiO_2$, petroleum ether/ethyl acetate = 0:1) to obtain **S8** (250 mg) as a white solid. To a solution of **S8** (100 mg, 305 µmol) in dichloromethane (2 mL) were added diisopropylethylamine (79.0 mg, 611 µmol) and acryloyl chloride (55.3 mg, 611 µmol). The mixture was stirred at 25 °C for 1 hour. Upon completion, the reaction mixture was concentrated under reduced pressure to give a residue. The residue was purified by preparative TLC ($SiO_2$, petroleum ether/EtOAc = 1:2) and prep-HPLC (column: Waters Xbridge 150mm×25mm×5µm; mobile phase: [water (10mM $NH_4HCO_3$)-$CH_3CN$]; B%: 35%-68%, 8min). It was further separated by SFC (Column: Chiralpak AD-3 50×4.6mm I.D., 3µm Mobile phase: Phase A: $CO_2$, and Phase B: i-PrOH (0.05% diethylamine); Gradient elution: 40% i-PrOH (0.05% diethylamine) in $CO_2$; Flow rate: 3mL/min; Column Temp: 35 ºC; Back Pressure: 100 Bar) to obtain **MY-11B** (52.0 mg, 53% yield over two steps) as a white solid.

**HRMS ESI-TOF** m/z calculated for $C_{24}H_{20}N_3O_2$ $[M+H]^+$ 382.1550. Found 382.1542.

**$^1$H NMR** (400 MHz, DMSO-*d*$_6$, mixture of rotamers): δ 10.4-10.2 (m, 1H), 8.90 (d, *J* = 4.0 Hz, 1H), 8.37 (d, *J* = 8.0 Hz, 1H), 8.21-8.12 (m, 1H), 7.65-7.56 (m, 2H), 7.45 (t, *J* = 8.0 Hz, 1H), 7.40-7.34 (m, 2H), 7.26-7.16 (m, 2H), 6.62-6.10 (m, 2H), 5.84 (d, *J* = 9.9 Hz, 1H), 5.70-5.40 (m, 1H), 4.65-4.55 (m, 1H), 4.50-4.22 (m, 2H), 4.10-4.00 (m, 1H).

**Synthesis of MY-11A**

Prepared in analogous fashion from *ent*-**S6**.

**(2*S*,3*R*)-1-acryloyl-3-(4-bromophenyl)-*N*-(quinolin-8-yl)azetidine-2-carboxamide (MY-11A) (9)**

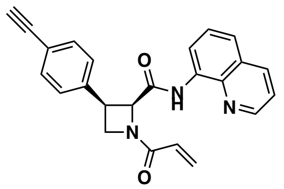

**HRMS ESI-TOF** m/z calculated for $C_{24}H_{20}N_3O_2$ $[M+H]^+$ 382.1550; Found 382.1540.

**$^1$H NMR** (400 MHz, $CD_3OD$): δ 8.85 (dd, *J* = 4.3, 1.7 Hz, 1H), 8.29 (dd, *J* = 8.3,

1.7 Hz, 1H), 8.26-8.11 (m, 1H), 7.60 (dd, *J* = 8.3, 1.3 Hz, 1H), 7.55 (dd, *J* = 8.3, 4.2 Hz, 1H), 7.44 (t, *J* = 8.0 Hz, 1H), 7.39 (d, *J* = 7.9 Hz, 2H), 7.21 (d, *J* = 7.8 Hz, 2H), 6.83-6.23 (m, 2H), 6.09-5.36 (m, 2H), 4.80-4.40 (m, 4H), 3.36 (s, 1H).

##### Synthesis of MY-45A and MY-45B

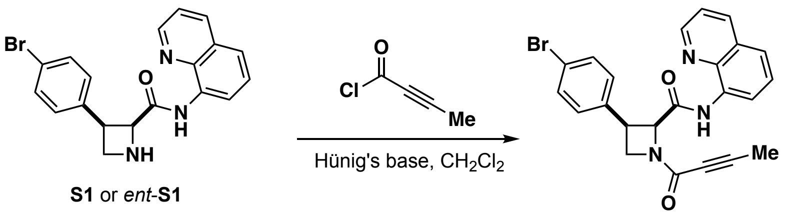

**S1** or *ent*-**S1**

**MY-45A** (**11**) or **MY-45B** (**12**)

**General procedure A: preparation of but-2-ynoyl chloride solution.** $PCl_5$ (115 mg, 0.55 mmol, 1.1 equiv) was added to an ice-cold suspension of but-2-ynoic acid (42.0 mg, 0.50 mmol, 1.0 equiv) in dichloromethane (1 mL). The ice bath was removed and the reaction mixture was stirred at room temperature until it turned into a clear solution (typically 30 minutes to 1 hour).

**General procedure B: butynamide formation.** But-2-ynoyl chloride (0.5 M solution in dichloromethane, 1.5 equiv), prepared according to general procedure A, was slowly added to a solution of the corresponding amine (1.0 equiv) and Hünig's base (3.0 equiv) in dichloromethane (0.05 M) at 0 °C. The reaction was allowed to warm to room temperature and stirred until complete consumption of starting material (as monitored by TLC). The reaction was quenched by addition of sat. aq. $NaHCO_3$ and was extracted with EtOAc (3×). The combined organic layers were dried over anhydrous sodium sulfate, filtered, and concentrated under reduced pressure. The residue was purified by flash chromatography and preparative TLC as indicated.

##### (2*S*,3*R*)-3-(4-bromophenyl)-1-(but-2-ynoyl)-*N*-(quinolin-8-yl)azetidine-2-carboxamide (MY-45A) (11)

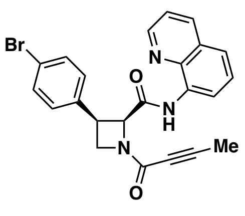

Following general procedure B using **S1** (16.9 mg, 0.044 mmol, 1.0 equiv).(Maetani et al., 2017) Purification by flash column chromatography ($SiO_2$, dichloromethane/acetone = 100:0 to 10:1) followed by preparative TLC ($SiO_2$, $CHCl_3$/acetone = 9:1) provided **MY-45A** as a white foam (9.9 mg, 50% yield).

**HRMS ESI-TOF** m/z calculated for $C_{23}H_{19}BrN_3O_2$ $[M+H]^+$ 448.0655. Found 448.0648.

**$^1$H NMR** (400 MHz, $CDCl_3$, mixture of rotamers): δ 10.54 (s, 0.6H), 10.35 (s, 0.4H), 8.95-8.77 (m, 1H), 8.40 (d, *J* = 7.4 Hz, 1H), 8.21-8.09 (m, 1H), 7.57-7.38 (m, 3H), 7.32-7.20 (m, 4H), 5.40 (d, *J* = 9.7 Hz, 0.6H), 5.29 (d, *J* = 9.7 Hz, 0.4H), 4.69-4.53 (m, 1H), 4.53-4.32 (m, 1H), 4.32-4.15 (m, 1H), 2.10 (s, 1H), 1.79 (s, 2H).

**(2*R*,3*S*)-3-(4-bromophenyl)-1-(but-2-ynoyl)-*N*-(quinolin-8-yl)azetidine-2-carboxamide (MY-45B) (12)**

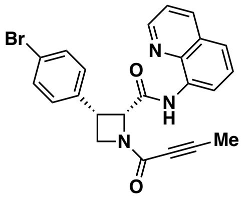

Following general procedure B using *ent*-**S1** (18.7 mg, 0.049 mmol, 1.0 equiv).(Maetani et al., 2017) Purification by flash column chromatography ($SiO_2$, dichloromethane/acetone = 100:0 to 10:1) followed by preparative TLC ($SiO_2$, $CHCl_3$/acetone = 9:1) provided **MY-45B** as a white foam (10.3 mg, 47% yield).

**HRMS ESI-TOF** m/z calculated for $C_{23}H_{19}BrN_3O_2$ $[M+H]^+$ 448.0655. Found 448.0644.

**$^1$H NMR** (400 MHz, $CDCl_3$, mixture of rotamers): δ 10.55 (s, 0.6H), 10.35 (s, 0.4 H), 8.98-8.74 (m, 1H), 8.40 (d, *J* = 7.4 Hz, 1H), 8.22-7.98 (m, 1H), 7.73-7.40 (m, 3H), 7.30-7.19 (m, 4H), 5.40 (d, *J* = 9.7 Hz, 0.6H), 5.29 (d, *J* = 9.7 Hz, 0.4H), 4.72-4.50 (m, 1H), 4.52-4.32 (m, 1H), 4.32-4.08 (m, 1H), 2.10 (s, 1H), 1.80 (s, 2H).

#### Synthesis of tryptoline probes

**EV-96, EV-97, EV-98, EV-99** were prepared as reported previously (Vinogradova et al., 2020).

##### methyl (1*R*,3*S*)-2-acryloyl-1-(benzo[*d*][1,3]dioxol-5-yl)-2,3,4,9-tetrahydro-1*H*-pyrido[3,4-*b*] indole-3-carboxylate (EV-96) (5)

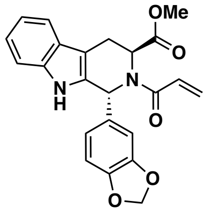

**HRMS ESI-TOF** m/z calculated for $C_{23}H_{21}N_2O_5$ $[M+H]^+$ 405.1445. Found 405.1441.

**$^1$H NMR** (400 MHz, $CD_3OD$): δ 7.44 (d, *J* = 7.8 Hz, 1H), 7.33-7.17 (m, 1H), 7.12-7.03 (m, 1H), 7.00 (t, *J* = 7.4 Hz, 1H), 6.95-6.85 (m, 2H), 6.84-6.65 (m, 2H), 6.30-6.04 (m, 2H), 6.00-5.79 (m, 2H), 5.70 (dd, *J* = 10.6, 1.8 Hz, 1H), 5.50-4.95 (m, 1H), 3.73-3.51 (m, 3H), 3.50-3.39 (m, 1H), 3.26-3.14 (m, 1H), 1 exchangeable proton not observed.

##### methyl (1*S*,3*R*)-2-acryloyl-1-(benzo[*d*][1,3]dioxol-5-yl)-2,3,4,9-tetrahydro-1*H*-pyrido[3,4-*b*] indole-3-carboxylate (EV-97) (6)

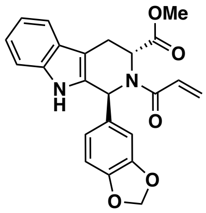

**HRMS ESI-TOF** m/z calculated for $C_{23}H_{21}N_2O_5$ $[M+H]^+$ 405.1445. Found 405.1450.

**$^1$H NMR** (400 MHz, $CD_3OD$): δ 7.44 (d, *J* = 8.0 Hz, 1H), 7.32-7.15 (m, 1H), 7.11-7.04 (m, 1H), 7.00 (t, *J* = 7.4 Hz, 1H), 6.95-6.86 (m, 2H), 6.85-6.68 (m, 2H), 6.30-6.05 (m, 2H), 6.00-5.79 (m, 2H), 5.70 (dd, *J* = 10.6, 1.8 Hz, 1H), 5.55-4.95 (m, 1H), 3.70-3.51 (m, 3H), 3.50-3.37 (m, 1H), 3.28-3.10 (m, 1H), 1 exchangeable proton not observed.

##### methyl (1*S*,3*S*)-2-acryloyl-1-(benzo[*d*][1,3]dioxol-5-yl)-2,3,4,9-tetrahydro-1*H*-pyrido[3,4-*b*] indole-3-carboxylate (EV-98) (7)

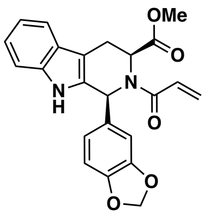

**HRMS ESI-TOF** m/z calculated for $C_{23}H_{21}N_2O_5$ $[M+H]^+$ 405.1445. Found 405.1444.

**$^1$H NMR** (400 MHz, $CD_3OD$): δ 7.52 (d, *J* = 7.8 Hz, 1H), 7.28 (d, *J* = 8.0 Hz, 1H), 7.15-7.09 (m, 1H), 7.08-7.03 (m, 1H), 7.02-6.81 (m, 2H), 6.80 (s, 1H), 6.73-6.39 (m,

2H), 6.49-6.20 (m, 1H), 5.90 (s, 2H), 5.82 (d, *J* = 8.0 Hz, 1H), 5.69-5.23 (m, 1H), 3.66-3.50 (m, 1H), 3.13 (s, 3H), 3.07-2.96 (m, 1H), 1 exchangeable proton not observed.

**methyl (1*R*,3*R*)-2-acryloyl-1-(benzo[*d*][1,3]dioxol-5-yl)-2,3,4,9-tetrahydro-1*H*-pyrido[3,4-*b*] indole-3-carboxylate (EV-99) (8)**

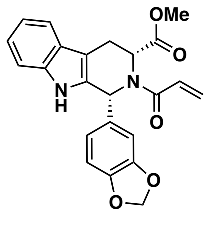

**HRMS ESI-TOF** m/z calculated for $C_{23}H_{21}N_2O_5$ $[M+H]^+$ 405.1445. Found 405.1442.

**$^1$H NMR** (400 MHz, $CD_3OD$): δ 7.52 (d, *J* = 8.0 Hz, 1H), 7.28 (d, *J* = 8.0 Hz, 1H), 7.15-7.09 (m, 1H), 7.08-7.02 (m, 1H), 7.02-6.84 (m, 2H), 6.80 (s, 1H), 6.73-6.65 (m, 1H), 6.62-6.53 (m, 1H), 6.38-6.21 (m, 1H), 5.90 (s, 2H), 5.87-5.78 (m, 1H), 5.68-5.25 (m, 1H), 3.66-3.56 (m, 1H), 3.13 (s, 3H), 3.09-2.97 (m, 1H), 1 exchangeable proton not observed.

#### Synthesis of WX-01-10

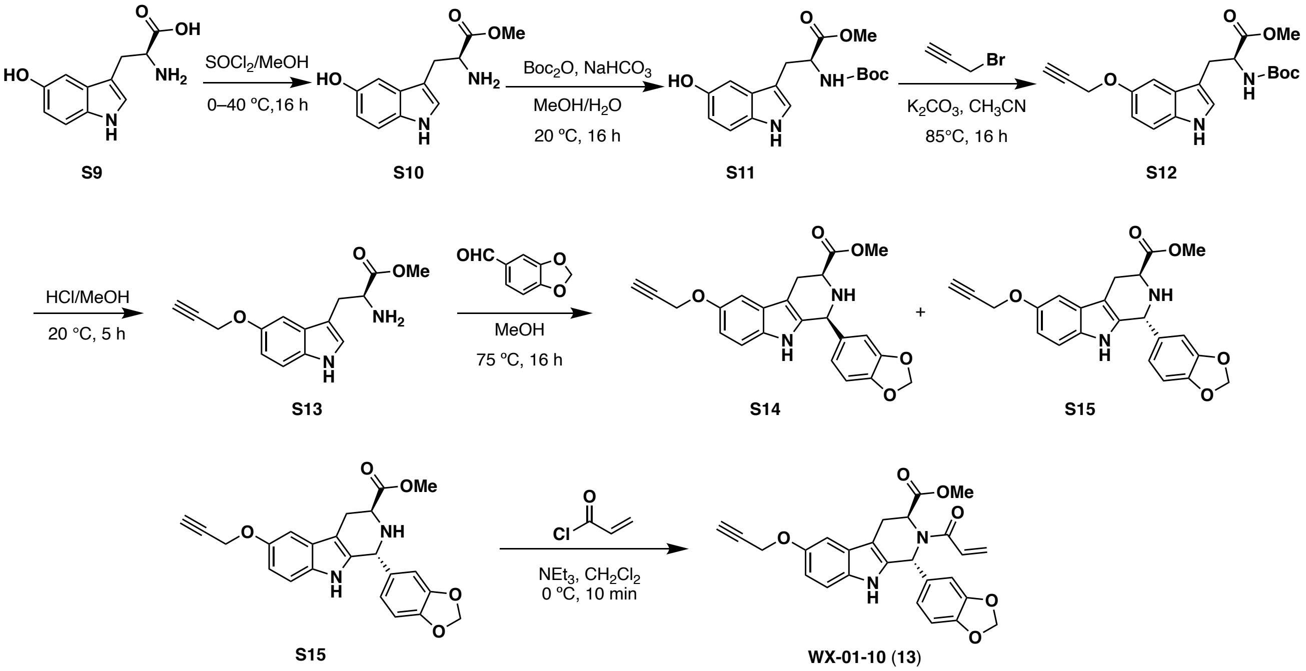

#### methyl (*S*)-2-((*tert*-butoxycarbonyl)amino)-3-(5-hydroxy-1*H*-indol-3-yl)propanoate (S11)

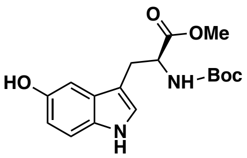

To a solution of **S9** (2.00 g, 9.08 mmol) in methanol (20 mL) was added dropwise thionyl chloride (2.00 g, 16.8 mmol, 1.22 mL) at 0 °C. The mixture was warmed and stirred at 40 °C for 16 hours. The reaction was monitored by LC-MS. The reaction mixture was concentrated under reduced pressure to give **S10** (1.90 g, crude) as a yellow oil, which was used for next step without purification. To a solution of **S10** (1.90 g, crude) in methanol (20 mL) and water (5 mL) were added $Boc_2O$ (3.54 g, 16.2 mmol, 3.73 mL) and sodium bicarbonate (2.04 g, 24.3 mmol). The mixture was stirred at 20 °C for 16 hours and the reaction was monitored by LC-MS. Upon completion, the reaction mixture was diluted with water (50 mL) and extracted with ethyl acetate (3×40 mL). The organic layers were combined, dried over anhydrous sodium sulfate, filtered, and concentrated under reduced pressure to give **S11** (3.00 g, crude) as a yellow solid, which was used in the next step without further purification.

**LC-MS** m/z calculated for $C_{17}H_{23}N_2O_5$ $[M+H]^+$ 335.2. Found 335.2.

**$^1$H NMR** (400 MHz, $CDCl_3$): δ 7.93 (br s, 1H), 7.21 (d, *J* = 8.7 Hz, 1H), 6.97 (dd, *J* = 7.9, 2.4 Hz, 2H), 6.77 (dd, *J* = 8.7, 2.4 Hz, 1H), 5.13-5.00 (m, 1H), 4.69-4.55 (m, 1H), 3.68 (s, 3H), 3.49 (d, *J* = 4.5 Hz, 1H), 3.25-3.17 (m, 2H), 1.57 (s, 9H).

**methyl (*S*)-2-((*tert*-butoxycarbonyl)amino)-3-(5-(prop-2-yn-1-yloxy)-1*H*-indol-3-yl)propanoate (S12)**

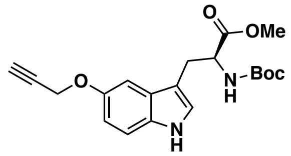

To a solution of **S11** (2.00 g, crude) in acetonitrile (30 mL) were added potassium carbonate (2.48 g, 17.9 mmol) and propargyl bromide (934 mg, 6.28 mmol, 677 µL, 80% w/w in toluene). The mixture was stirred at 85 °C for 16 hours. The reaction was monitored by LC-MS. The reaction mixture was extracted with EtOAc (3×40 mL). The organic layers were combined, dried over anhydrous sodium sulfate, filtered, and concentrated under reduced pressure to give a residue, which was purified by column chromatography ($SiO_2$, petroleum ether/EtOAc = 10:1 to 5:1) to give **S12** (2.30 g, quant.) as a white solid.

**LC-MS** m/z calculated for $C_{15}H_{17}N_2O_3$ [M–Boc+H]$^+$ 273.1. Found 273.1.

**$^1$H NMR** (400 MHz, $CDCl_3$): δ 8.00 (br s, 1H), 7.29-7.24 (m, 1H + $CHCl_3$), 7.11 (d, *J* = 2.4 Hz, 1H), 7.01 (d, *J* = 2.5 Hz, 1H), 6.92 (dd, *J* = 8.8, 2.4 Hz, 1H), 5.13-5.05 (m, 1H), 4.73 (d, *J* = 2.4 Hz, 2H), 4.68-4.61 (m, 1H), 3.68 (s, 3H), 3.24 (d, *J* = 5.6 Hz, 2H), 2.52 (t, *J* = 2.4 Hz, 1H), 1.42 (s, 9H).

**Compounds S14 and S15**

To a solution of **S12** (1.30 g, 3.49 mmol) in methanol (13 mL) was added HCl (4 M in MeOH, 9 mL). The mixture was stirred at 20 °C for 5 hours. The reaction was monitored by LC-MS. Upon completion, the reaction mixture was concentrated under reduced pressure to give **S13** (1.00 g, crude) as a yellow oil, which was used in the next step without purification. To a solution of **S13** (900 mg, 3.31 mmol) in methanol (15 mL) was added piperonal (595 mg, 3.97 mmol). The mixture was stirred at 75 °C for 16 hours. The reaction was monitored by LC-MS. Upon completion, the reaction mixture was diluted with water (20 mL) to adjust the pH to 8-9, then extracted with EtOAc (3×20 mL). The organic layers were combined, dried over anhydrous sodium sulfate, filtered, and concentrated under reduced pressure to give a residue. The residue was purified by column chromatography ($SiO_2$, petroleum ether/EtOAc = 1:0 to 2:1) to give **S14** (130 mg, 7.7% yield) and **S15** (130 mg, 7.2% yield) as yellow solids.

**methyl (1*S*,3*S*)-1-(benzo[*d*][1,3]dioxol-5-yl)-6-(prop-2-yn-1-yloxy)-2,3,4,9-tetrahydro-1*H*-pyrido [3,4-*b*]indole-3-carboxylate (S14)**

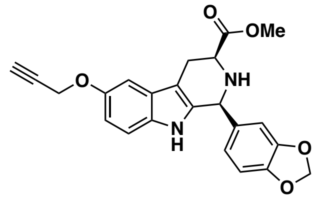

**LC-MS** m/z calculated for $C_{23}H_{21}N_2O_5$ $[M+H]^+$ 405.1. Found 405.2.

**$^1$H NMR** (400 MHz, $CDCl_3$): δ 7.34 (br s, 1H), 7.13 (d, *J* = 8.7 Hz, 1H), 7.09 (d, *J* = 2.5 Hz, 1H), 6.87 (ddd, *J* = 8.4, 5.6, 2.1 Hz, 2H), 6.84-6.75 (m, 2H), 5.95 (s, 2H), 5.21-5.10 (m, 1H), 4.79-4.68 (m, 2H), 3.95 (dd, *J* = 11.1, 4.3 Hz, 1H), 3.82 (s, 3H), 3.17 (ddd, *J* = 15.0, 4.2, 1.8 Hz, 1H), 2.97 (ddd, *J* = 15.0, 11.1, 2.5 Hz, 1H), 2.52 (t, *J* = 2.4 Hz, 1H), 1 exchangeable proton not observed.

**methyl (1*R*,3*S*)-1-(benzo[*d*][1,3]dioxol-5-yl)-6-(prop-2-yn-1-yloxy)-2,3,4,9-tetrahydro-1*H*-pyrido [3,4-*b*]indole-3-carboxylate (S15)**

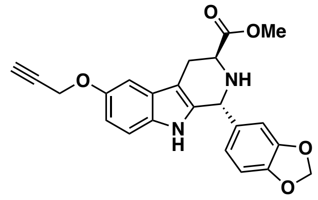

**LC-MS** m/z calculated for $C_{23}H_{21}N_2O_5$ $[M+H]^+$ 405.1. Found 405.2.

**$^1$H NMR** (400 MHz, $CDCl_3$): δ 7.46 (br s, 1H), 7.15 (d, *J* = 8.7 Hz, 1H), 7.14-7.08 (m, 1H), 6.93-6.78 (m, 1H), 6.75 (s, 3H), 5.99-5.89 (m, 2H), 5.35-5.29 (m, 1H), 4.74 (dd, *J* = 3.5, 2.4 Hz, 2H), 3.98 (t, *J* = 6.0 Hz, 1H), 3.72 (s, 3H), 3.22 (ddd, *J* = 15.3, 5.5, 1.3 Hz, 1H), 3.09 (ddd, *J* = 15.4, 6.6, 1.5 Hz, 1H), 2.52 (t, *J* = 2.4 Hz, 1H), 1 exchangeable proton not observed.

**methyl (1*R*,3*S*)-2-acryloyl-1-(benzo[*d*][1,3]dioxol-5-yl)-6-(prop-2-yn-1-yloxy)-2,3,4,9-tetra-hydro-1*H*-pyrido[3,4-*b*]indole-3-carboxylate (WX-01-10) (13)**

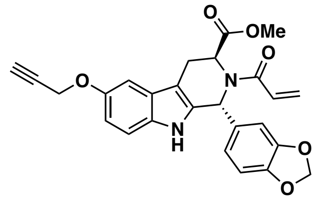

To a solution of **S15** (80.00 mg, 198 µmol) in dichloromethane (2 mL) were added triethylamine (40.0 mg, 396 µmol, 55.1 uL) and acryloyl chloride (17.9 mg, 198 µmol, 16.1 uL). The mixture was stirred at 0 °C for 10 min and the reaction was monitored by LC-MS. Upon completion, the reaction mixture was concentrated under reduced pressure to give a residue, which was purified by prep-HPLC (column: Phenomenex Luna C18 150*25mm*10um; mobile phase: [water (0.225%FA)-ACN]; B%: 39%-69%, 10min) and preparatory TLC ($SiO_2$, petroleum ether/EtOAc = 1:1) to give **WX-01-10** (18.0 mg, 20% yield) as a white solid.

**HRMS ESI-TOF** m/z calculated for $C_{26}H_{23}N_2O_6$ $[M+H]^+$ 459.1551. Found 459.1551.

**$^1$H NMR** (400 MHz, $CDCl_3$): δ 7.64 (br s, 1H), 7.17 (d, *J* = 8.8 Hz, 1H), 7.08 (d, *J* = 2.4 Hz, 1H), 6.89 (dd, *J* = 8.8, 2.4 Hz, 1H), 6.84 (br s, 1H), 6.82-6.74 (m, 2H), 6.57 (br dd, *J* = 16.6, 10.8 Hz, 1H), 6.30 (dd, *J* =1 6.6, 1.6 Hz, 1H), 6.10 (s, 1H), 5.93 (br d, *J* = 6.4 Hz, 2H), 5.66 (br d, *J* = 10.6 Hz, 1H), 5.10 (br s, 1H), 4.73 (d, *J* = 2.4 Hz, 2H), 3.66 (s, 3H), 3.54 (br d, *J* = 15.2 Hz, 1H), 3.25 (br s, 1H), 2.52 (t, *J* = 2.4 Hz, 1H).

##### Synthesis of WX-01-12

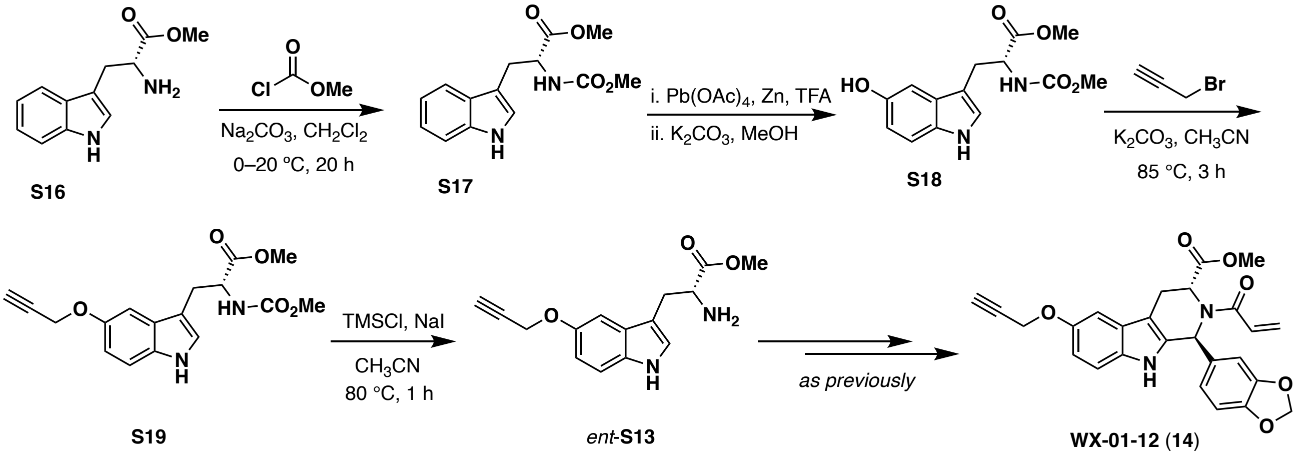

##### methyl (methoxycarbonyl)-*D*-tryptophanate (S17)

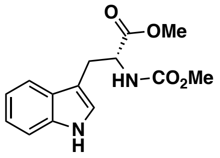

To a precooled (0 ºC) solution of **S16** (1.05 g, 4.12 mmol, HCl) in dichloromethane (20 mL) were added sodium carbonate (0.584 g, 6.18 mmol) and methyl chloroformate (0.655 g, 6.18 mmol, 0.48 mL). The mixture was allowed to gradually warm to 20 °C overnight (20 h). The reaction mixture was diluted with water (100 mL) and dichloromethane (50 mL). The organic layer was washed sequentially with water, sat. aq. sodium bicarbonate, and brine. The organic layer was dried over anhydrous sodium sulfate, filtered, and concentrated under reduced pressure to give **S17** (1.14 g, 4.12 mmol, quant.), which was used in the next step without further purification.

**LC-MS** m/z calculated for $C_{14}H_{17}N_2O_4$ $[M+H]^+$ 277.1. Found 277.1.

**$^1$H NMR** (400 MHz, $CDCl_3$) δ 8.10 (br s, 1H), 7.54 (d, *J* = 7.9 Hz, 1H), 7.36 (dt, *J* = 8.1, 0.9 Hz, 1H), 7.20 (ddd, *J* = 8.2, 7.0, 1.3 Hz, 1H), 7.13 (ddd, *J* = 8.0, 7.0, 1.1 Hz, 1H), 7.01 (d, *J* = 2.2 Hz, 1H), 5.23

(d, $J$ = 8.3 Hz, 1H), 4.71 (td, $J$ = 5.8, 5.8 Hz, 1H), 3.68 (s, 3H), 3.67 (s, 3H), 3.31 (d, $J$ = 5.5 Hz, 2H).

**methyl (*R*)-3-(5-hydroxy-1*H*-indol-3-yl)-2-((methoxycarbonyl)amino)propanoate (S18)**

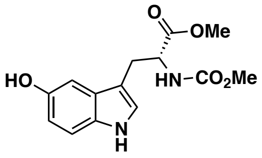

**S17** (1.14 g, 4.12 mmol) was dissolved in trifluoroacetic acid (12 mL) and the solution was stirred at 20 °C for 2.5 h. Next, the reaction mixture was cooled to 12 ºC (1,4 dioxane dry ice bath) and a solution of lead tetraacetate (4.02 g, 9.08 mmol; best results were obtained using fresh Strem Chemicals batch) in dichloromethane (80 mL) was added over 10 min. The brown mixture was stirred for 1.5 h at the same temperature before adding zinc (1.35 g, 20.6 mmol) and warming the reaction to 20 °C over 45 min (the reaction becomes amber in color). The reaction was diluted with water (100 mL) and stirred vigorously over 30 min before extracting with dichloromethane (3×50 mL). The combined organic layers were filtered through a silica plug and concentrated under reduced pressure. The resulting brown oil was dissolved in methanol (20 mL) and treated with potassium carbonate (0.375 g, 2.71 mmol) overnight (18 h) at 20 °C to solvolyze any trifluoroacetate ester formed over previous steps. The resulting solution was diluted in 50% sat. aq. NaCl and extracted with dichloromethane (3×50 mL). The combined organic layers were dried over anhydrous sodium sulfate, filtered, and concentrated under reduced pressure. The residue was purified by column chromatography ($SiO_2$, hexanes/EtOAc = 1:1 with 0.1% acetic acid) to give **S18** as a tan oil/foam (0.605 g, 4.13 mmol, 50%).

**LC-MS** m/z calculated for $C_{14}H_{17}N_2O_5$ $[M+H]^+$ 293.3. Found 293.0.

**$^1$H NMR** (400 MHz, $CDCl_3$) δ 8.10 (br s, 1H), 7.17 (dd, $J$ = 8.8, 2.6 Hz, 1H), 6.99-6.92 (m, 2H), 6.77 (dd, $J$ = 8.6, 2.4 Hz, 1H), 5.43-5.31 (m, 1H), 4.67 (ddd, $J$ = 6.4, 6.3, 6.3 Hz, 1H), 3.67 (s, 3H), 3.65 (s, 3H), 3.20 (d, $J$ = 5.7 Hz, 2H), 1 exchangeable proton not observed.

**methyl (*R*)-2-((methoxycarbonyl)amino)-3-(5-(prop-2-yn-1-yloxy)-1*H*-indol-3-yl)propanoate (S19)**

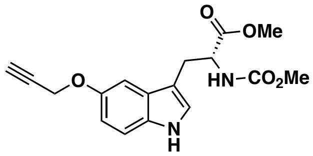

To a solution of **S18** (690 mg, 2.36 mmol) in acetonitrile (20 mL) were added potassium carbonate (979 mg, 7.08 mmol) and propargyl bromide (351 mg, 2.36 mmol, 254 µL, 80% w/w in toluene), and the mixture was stirred at 85 °C for 3 hours. The reaction was monitored by TLC and LC-MS. The reaction mixture was diluted with water (100 mL) and extracted with ethyl acetate (50 mL × 3). The combined organic layers were dried over anhydrous sodium sulfate, filtered, and concentrated under reduced pressure. The resulting residue was purified by column chromatography ($SiO_2$, petroleum ether/EtOAc = 1 : 1 to 1 : 1) to give **S19** (550 mg, 1.66 mmol, 71% yield) as a yellow oil.

**LC-MS** m/z calculated for $C_{17}H_{19}N_2O_5$ $[M+H]^+$ 331.1. Found 331.1.

**methyl (*R*)-2-amino-3-(5-(prop-2-yn-1-yloxy)-1*H*-indol-3-yl)propanoate (*ent*-S13)**

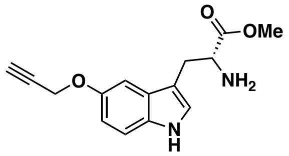

To a solution of **S19** (400 mg, 1.21 mmol) in acetonitrile (4 mL) was added trimethylchlorosilane (263 mg, 2.42 mmol, 307 μL) and sodium iodide (363 mg, 2.42 mmol). The mixture was stirred at 80 °C for 1 hour. This reaction was monitored by LC-MS. The reaction mixture was diluted with water (100 mL) and extracted with EtOAc (3×30 mL). The combined organic layers were dried over anhydrous sodium sulfate, filtered, and concentrated under reduced pressure to give *ent*-**S13** (510 mg, crude) as a yellow oil. It was used in next step directly without purification.

The remaining transformations were performed as described previously for **WX-01-10**.

**methyl (1*S*,3*R*)-2-acryloyl-1-(benzo[*d*][1,3]dioxol-5-yl)-6-(prop-2-yn-1-yloxy)-2,3,4,9-tetra-hydro-1*H*-pyrido[3,4-*b*]indole-3-carboxylate (WX-01-12) (14)**

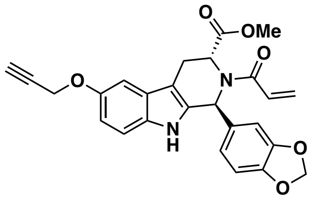

**HRMS ESI-TOF** m/z calculated for $C_{26}H_{23}N_2O_6$ $[M+H]^+$ 459.1551. Found 459.1552.

**$^1$H NMR** (400 MHz, $CDCl_3$): δ 7.74 (br s, 1H), 7.15 (d, *J* = 8.8 Hz, 1H), 7.07 (d, *J* = 2.4 Hz, 1H), 6.87 (dd, *J* = 8.8, 2.5 Hz, 1H), 6.85-6.70 (m, 3H), 6.56 (dd, *J* = 16.7, 10.5 Hz, 1H), 6.29 (dd, *J* = 16.7, 1.8 Hz, 1H), 6.09 (s, 1H), 5.99-5.86 (m, 2H), 5.64 (d, *J* = 10.5 Hz, 1H), 5.17-5.00 (m, 1H), 4.72 (d, *J* = 2.4 Hz, 2H), 3.64 (s, 3H), 3.59-3.45 (m, 1H), 3.39-3.03 (m, 1H), 2.51 (t, *J* = 2.4 Hz, 1H).

#### Synthesis of WX-02-23

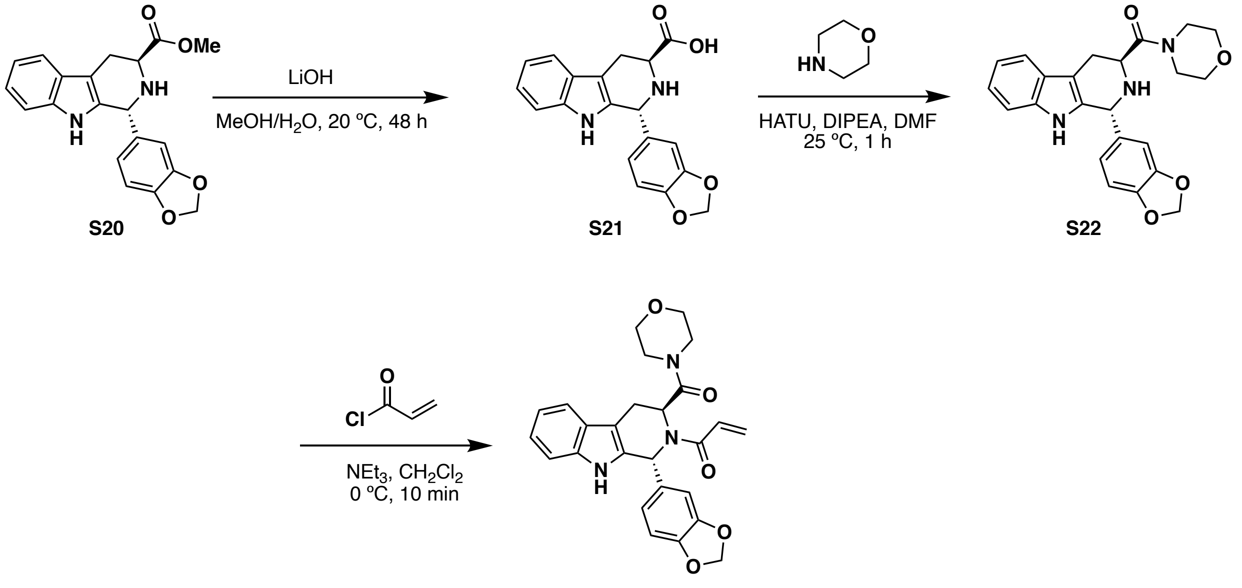

**WX-02-23** (**15**)

#### (1*R*,3*S*)-1-(benzo[*d*][1,3]dioxol-5-yl)-2,3,4,9-tetrahydro-1*H*-pyrido[3,4-*b*]indole-3-carboxylic acid (S21)

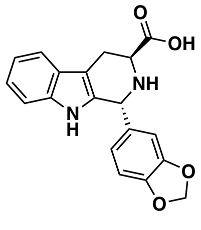

To a solution of **S20** (500 mg, 1.43 mmol) (Vinogradova et al., 2020) in methanol (10 mL) were added lithium hydroxide monohydrate (71.9 mg, 1.71 mmol) and water (257 mg, 14.3 mmol, 257 µL). The mixture was stirred at 20 °C for 48 hours. The reaction was monitored by LC-MS. The reaction mixture was concentrated under reduced pressure to give a residue, which was suspended in toluene (10 mL) and concentrated under reduced pressure to remove residual water. **S21** (500 mg, 97% yield), obtained as a white solid was used in the next step without further purification.

**LC-MS** m/z calculated for $C_{19}H_{17}N_2O_4$ $[M+H]^+$ 337.1. Found 337.1.

**$^1$H NMR** (400 MHz, DMSO-*d*$_6$): δ 7.40 (d, *J* = 8.0 Hz, 1H), 7.30-7.10 (m, 3H), 7.05-6.89 (m, 2H), 6.80 (d, *J* = 8.0 Hz, 1H), 6.74 (m, 1H), 6.60-6.54 (m, 1H), 5.96 (s, 2H), 5.15 (m, 1H), 3.15-3.06 (m, 1H), 2.95-2.85 (m, 1H), 2.65-2.54 (m, 2H).

**((1*R*,3*S*)-1-(benzo[*d*][1,3]dioxol-5-yl)-2,3,4,9-tetrahydro-1*H*-pyrido[3,4-*b*]indol-3-yl)(morpholino)methanone (S22)**

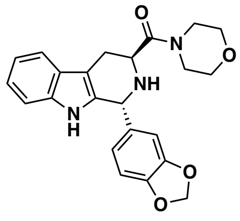

To a solution of **S21** (350 mg, 1.04 mmol) in *N,N*-dimethyl formamide (2 mL) were added HATU (594 mg, 1.56 mmol), morpholine (1.81 g, 20.8 mmol, 1.83 mL), and diisopropylethylamine (269 mg, 2.08 mmol, 363 μL). The mixture was stirred at 25 °C for 1 hour and the reaction was monitored by LC-MS. Upon completion, the reaction mixture was diluted with water (20 mL) and extracted with ethyl acetate (3× 20 mL). The organic layers were combined, dried over anhydrous sodium sulfate, filtered, and concentrated under reduced pressure to give a residue, which was purified by prep-HPLC (column: Waters Xbridge 150mm×25mm×5μm; mobile phase: [water (10mM $NH_4HCO_3$)-$CH_3CN$]; B%: 26%-59%,10min) to give **S22** (300 mg, 70% yield) as a white solid.

**LC-MS** m/z calculated for $C_{23}H_{24}N_3O_4$ $[M+H]^+$ 406.2. Found 406.2.

**$^1$H NMR** (400 MHz, $CD_3OD$): δ 7.50 (d, *J* = 8.0 Hz, 1H), 7.29 (d, *J* = 8.0 Hz, 1H), 7.13-7.06 (m, 1H), 7.06-7.00 (m, 1H), 6.80-6.72 (m, 2H), 6.68-6.62 (m, 1H), 5.94 (s, 2H), 5.27 (s, 1H), 3.97 (dd, *J* = 10.0, 5.0 Hz, 1 H), 3.78-3.47 (m, 8 H), 3.29-3.13 (m, 1 H), 3.05-2.90 (m, 1 H), 2 exchangeable protons not observed.

**1-((1*R*,3*S*)-1-(benzo[*d*][1,3]dioxol-5-yl)-3-(morpholine-4-carbonyl)-1,3,4,9-tetrahydro-2*H*-pyrido[3,4-*b*]indol-2-yl)prop-2-en-1-one (WX-02-23) (15)**

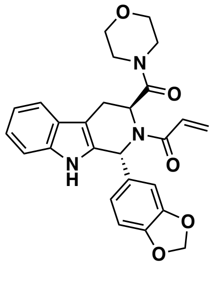

To a solution of **S22** (70.0 mg, 173 μmol) in dichloromethane (2 mL) were added triethylamine (34.9 mg, 345 μmol, 48.1 μL) and acryloyl chloride (15.6 mg, 173 μmol, 14.1 μL). The mixture was stirred at 0 °C for 10 min and the reaction was monitored by LC-MS. Upon completion, the reaction mixture was concentrated under reduced pressure to give a residue, which was purified by prep-HPLC (column: Waters Xbridge 150mm×25mm×5μm; mobile phase: [water (10mM $NH_4HCO_3$)-$CH_3CN$]; B%: 28%-58%,10min) to give **WX-02-23** (23.1 mg, 28% yield) as a white solid.

**HRMS ESI-TOF** m/z calculated for $C_{26}H_{26}N_3O_5$ $[M+H]^+$ 460.1867. Found 460.1867.

**$^1$H NMR** (400 MHz, $CD_3OD$): δ 7.47 (d, *J* = 7.8 Hz, 1H), 7.27 (d, *J* = 7.9 Hz, 1H), 7.08 (ddd, *J* = 8.2, 7.1, 1.3 Hz, 1H), 7.01 (ddd, *J* = 8.1, 7.1, 1.1 Hz, 1H), 6.98-6.72 (m, 4H), 6.42-6.32 (m, 1H), 6.24 (dd,

*J* = 16.6, 1.8 Hz, 1H), 5.96-5.88 (m, 2H), 5.75 (dd, *J* = 10.6, 1.9 Hz, 1H), 5.14-5.01 (m, 1H), 3.68-3.44 (m, 5H), 3.44-3.12 (m, 5H + solvent residual peak), 1 exchangeable proton not observed.

**Synthesis of WX-02-43**

Prepared in analogous fashion from *ent*-**S20** (Vinogradova et al., 2020).

**1-((1*S*,3*R*)-1-(benzo[*d*][1,3]dioxol-5-yl)-3-(morpholine-4-carbonyl)-1,3,4,9-tetrahydro-2*H*-pyrido [3,4-*b*]indol-2-yl)prop-2-en-1-one (WX-02-43) (16)**

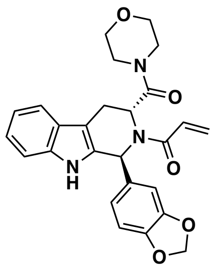

**HRMS ESI-TOF** m/z calculated for $C_{26}H_{26}N_3O_5$ $[M+H]^+$ 460.1867. Found 460.1870.

**$^1$H NMR** (400 MHz, $CD_3OD$): δ 7.47 (d, J = 7.8 Hz, 1H), 7.27 (d, J = 8.0 Hz, 1H), 7.08 (t, J = 7.5 Hz, 1H), 7.01 (t, J = 7.4 Hz, 1H), 6.98-6.66 (m, 4H), 6.42-6.34 (m, 1H), 6.24 (d, J = 16.6 Hz, 1H), 5.93 (s, 2H), 5.76 (dd, J = 10.6, 1.4 Hz, 1H), 5.13-5.02 (m, 1H), 3.72-3.44 (m, 5H), 3.44-3.11 (m, 5H + solvent residual peak), 1 exchangeable proton not observed.

#### Synthesis of EV-97-ctrl

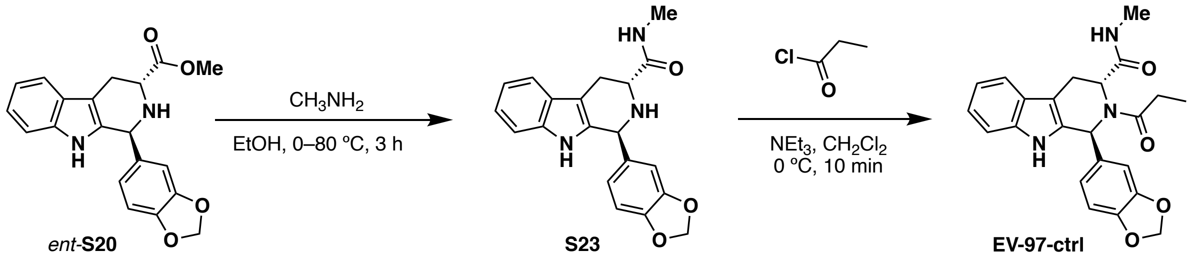

**(1*S*,3*R*)-1-(benzo[*d*][1,3]dioxol-5-yl)-*N*-methyl-2,3,4,9-tetrahydro-1*H*-pyrido[3,4-*b*]indole-3-carboxamide (S23)**

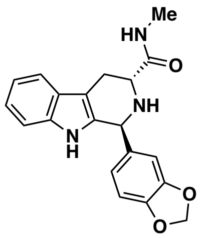

A solution of *ent*-**S20** (1.50 g, 4.28 mmol) (Vinogradova et al., 2020) in ethanol (10 mL) was degassed by purging with nitrogen 3 times, and methylamine (20.3 g, 40% w/w in water) was added. The mixture was stirred at 0 °C for 1 hour, then at 80 °C for 2 hours under nitrogen atmosphere. The reaction was monitored by LC-MS. The reaction mixture was concentrated under reduced pressure to give a residue, which was purified by column chromatography ($SiO_2$, petroleum ether/EtOAc = 1:0 to 1:1) to give **S23** (1.25 g, 3.28 mmol, 77% yield) as a yellow solid.

**LC-MS** m/z calculated for $C_{20}H_{20}N_3O_3$ $[M+H]^+$ 350.1. Found 350.1.

**$^1$H NMR** (400 MHz, $CD_3OD$): δ 7.47 (d, *J* = 8.0 Hz, 1H), 7.26 (d, *J* = 8.0 Hz, 1H), 7.10-7.04 (m, 1H), 7.03-6.97 (m, 1H), 6.77-6.70 (m, 2H), 6.68-6.60 (m, 1H), 5.89 (s, 2H), 5.24 (s, 1H), 3.65-3.55 (m, 1H), 3.16-3.07 (m, 1H), 2.89-2.78 (m, 1H), 2.75 (s, 3H), 3 exchangeable protons not observed.

**(1*S*,3*R*)-1-(benzo[*d*][1,3]dioxol-5-yl)-*N*-methyl-2-propionyl-2,3,4,9-tetrahydro-1*H*-pyrido[3,4-*b*]indole-3-carboxamide (EV-97-ctrl) (18)**

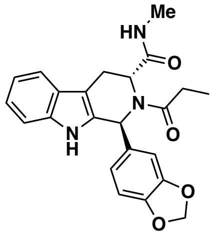

To a solution of **S23** (50.0 mg, 143 µmol) in dichloromethane (2 mL) were added triethylamine (21.7 mg, 215 µmol, 29.9 µL) and propionyl chloride (13.2 mg, 143 µmol, 13.2 µL). The mixture was stirred at 0 °C for 10 min and the reaction was monitored by LC-MS. The reaction mixture was concentrated under reduced pressure to give a residue, which was purified by prep-HPLC (column: Waters Xbridge 150mm×25mm×5µm; mobile phase: [water (10mM $NH_4HCO_3$)-$CH_3CN$]; B%: 25%-55%, 10min) to give **EV-97-ctrl** (37.6 mg, 92.8 µmol, 65% yield) as an off-white solid.

**HRMS ESI-TOF** m/z calculated for $C_{23}H_{24}N_3O_4$ $[M+H]^+$ 406.1761. Found 406.1760.

**$^1$H NMR** (400 MHz, $CD_3OD$): δ 7.42 (d, *J* = 8.0 Hz, 1H), 7.27 (d, *J* = 8.0 Hz, 1H), 7.03 (t, *J* = 7.6 Hz, 1H), 6.97 (t, *J* = 7.3 Hz, 1H), 6.94-6.87 (m, 2H), 6.79-6.67 (m, 1H), 6.22 (s, 1H), 5.88 (s, 2H), 5.18 (m, 1H), 3.53-3.30 (m, 2H + solvent residual peak), 2.61 (s, 3H), 2.58-2.51 (m, 1H), 2.37-2.20 (m, 1H), 1.10-0.94 (m, 3H), 2 exchangeable protons not observed.

**Synthesis of EV-96-ctrl**

Prepared in analogous fashion from **S20** (Vinogradova et al., 2020).

**(1*S*,3*R*)-1-(benzo[*d*][1,3]dioxol-5-yl)-*N*-methyl-2-propionyl-2,3,4,9-tetrahydro-1*H*-pyrido[3,4-*b*]indole-3-carboxamide (EV-96-ctrl) (17)**

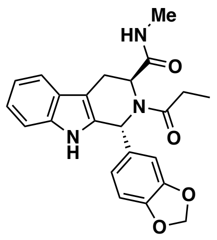

**HRMS ESI-TOF** m/z calculated for $C_{23}H_{24}N_3O_4$ $[M+H]^+$ 406.1761. Found 406.1761.

**$^1$H NMR** (400 MHz, $CD_3OD$): δ 7.42 (d, *J* = 8.0 Hz, 1H), 7.27 (d, *J* = 8.0 Hz, 1H), 7.09-7.02 (m, 1H), 7.01-6.95 (m, 1H), 6.93-6.87 (m, 2H), 6.80-6.68 (m, 1H), 6.23 (s, 1H), 5.91-5.80 (m, 2H), 5.16 (s, 1H), 3.56-3.30 (m, 2H + solvent residual peak), 2.61 (s, 3H), 2.59-2.48 (m, 1H), 2.36-2.25 (m, 1H), 1.03 (m, 3H), 2 exchangeable protons not observed.

**Synthesis of EV-99-ctrl**

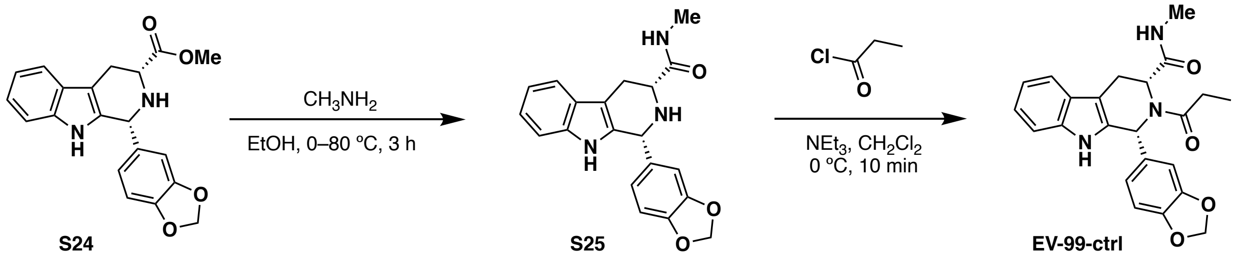

**(1*R*,3*R*)-1-(benzo[*d*][1,3]dioxol-5-yl)-*N*-methyl-2,3,4,9-tetrahydro-1*H*-pyrido[3,4-*b*]indole-3-carboxamide (S25)**

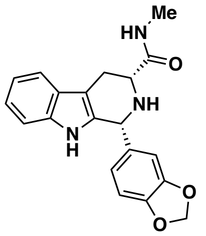

A solution of **S24** (1.50 g, 4.28 mmol) (Vinogradova et al., 2020) in ethanol (10 mL) was degassed by purging with nitrogen 3 times, and methylamine (20.3 g, 40% w/w in water) was added. The mixture was stirred at 0 °C for 1 hour, then 80 °C for 2 hours under nitrogen atmosphere. The reaction was monitored by LC-MS. Upon completion, the reaction mixture was concentrated under reduced pressure to give a residue, which was purified by column chromatography ($SiO_2$, petroleum ether/EtOAc = 1:0 to 1:1) to give **S25** (1.45 g, 4.03 mmol, 94% yield) as a yellow solid.

**LC-MS** m/z calculated for $C_{20}H_{20}N_3O_3$ $[M+H]^+$ 350.1. Found 350.1.

**$^1$H NMR** (400 MHz, $CD_3OD$): δ 7.44 (d, *J* = 8.0 Hz, 1H), 7.22 (d, *J* = 8.0 Hz, 1H), 7.08-6.95 (m, 2H), 6.90-6.75 (m, 3H), 5.91 (s, 2H), 5.10 (m, 1H), 3.75-3.65 (m, 1H), 3.15-3.05 (m, 1H), 2.90-2.80 (m, 1H), 2.79 (s, 3H), 3 exchangeable protons not observed.

**(1*R*,3*R*)-1-(benzo[*d*][1,3]dioxol-5-yl)-*N*-methyl-2-propionyl-2,3,4,9-tetrahydro-1*H*-pyrido[3,4-*b*]indole-3-carboxamide (EV-99-ctrl) (20)**

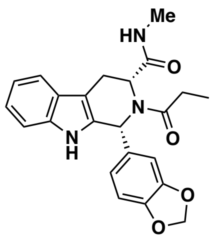

To a solution of **S25** (50.0 mg, 143 μmol) in dichloromethane (2 mL) were added triethylamine (21.7 mg, 215 μmol, 29.9 μL) and propionyl chloride (13.2 mg, 143 μmol, 13.2 μL). The mixture was stirred at 0 °C for 10 min and the reaction was monitored by LC-MS. Upon completion, the reaction mixture was concentrated under reduced pressure. The resulting residue was purified by prep-HPLC (column: Waters Xbridge 150mm×25mm×5μm; mobile phase: [water (10mM $NH_4HCO_3$)-$CH_3CN$]; B%: 29%-59%, 10 min) to give **EV-99-ctrl** (36.9 mg, 90.9 μmol, 64% yield) as a white solid.

**HRMS ESI-TOF** m/z calculated for $C_{23}H_{24}N_3O_4$ $[M+H]^+$ 406.1761. Found 406.1756.

**$^1$H NMR** (400 MHz, $CD_3OD$): δ 7.53 (d, *J* = 8.0 Hz, 1H), 7.27 (d, *J* = 8.0 Hz, 1H), 7.12-6.95 (m, 3H), 6.86-6.60 (m, 3H), 5.89 (s, 2H), 5.20-5.00 (m, 1H), 3.75-3.55 (m, 1H), 2.97 (dd, *J* = 15.7, 6.8 Hz, 1H), 2.80-2.55 (m, 2H), 2.21 (s, 3H), 1.21 (t, *J* = 8.0 Hz, 3H), 2 exchangeable protons not observed.

**Synthesis of EV-98-ctrl**

Prepared in analogous fashion from *ent*-**S24** (Vinogradova et al., 2020).

**(1*S*,3*S*)-1-(benzo[*d*][1,3]dioxol-5-yl)-*N*-methyl-2-propionyl-2,3,4,9-tetrahydro-1*H*-pyrido[3,4-*b*]indole-3-carboxamide (EV-98-ctrl) (19)**

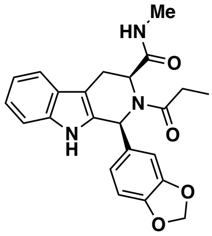

**HRMS ESI-TOF** m/z calculated for $C_{23}H_{24}N_3O_4$ $[M+H]^+$ 406.1761. Found 406.1762.

**$^1$H NMR** (400 MHz, $CD_3OD$): δ 7.54 (d, *J* = 8.0 Hz, 1H), 7.28 (d, *J* = 8.0 Hz, 1H), 7.14-7.09 (m, 1H), 7.07-6.90 (m, 2H), 6.84 (m, 1H), 6.80-6.62 (m, 2H), 5.90 (s, 2H), 5.25-5.00 (m, 1H), 3.75-3.55 (m, 1H), 2.96 (dd, *J* = 15.9, 6.8 Hz, 1H), 2.75-2.55 (m, 2H), 2.22 (s, 3H), 1.22 (t, *J* = 8.0 Hz, 3H), 2 exchangeable protons not observed.

#### Spectroscopic and chromatographic data

##### $^1H$ NMR of MY-1A (1) in $CDCl_3$

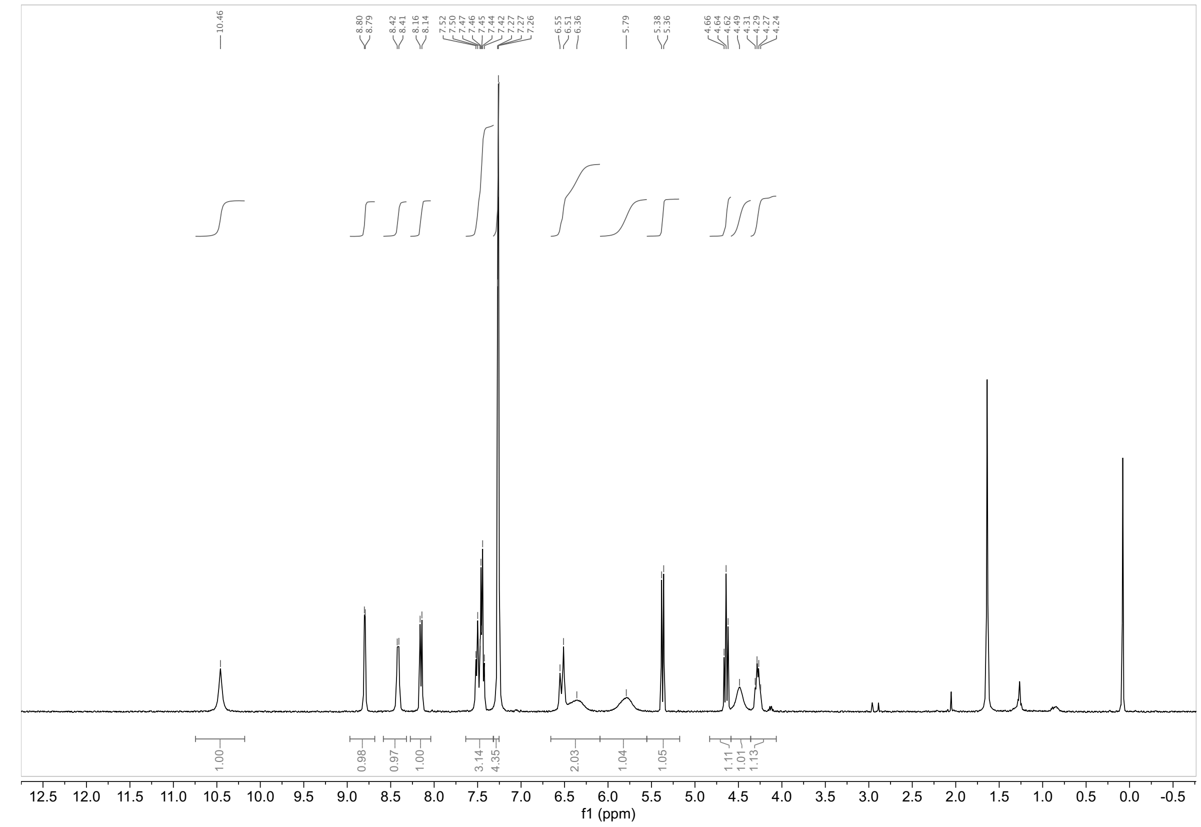

##### $^1H$ NMR of MY-1B (2) in $CDCl_3$

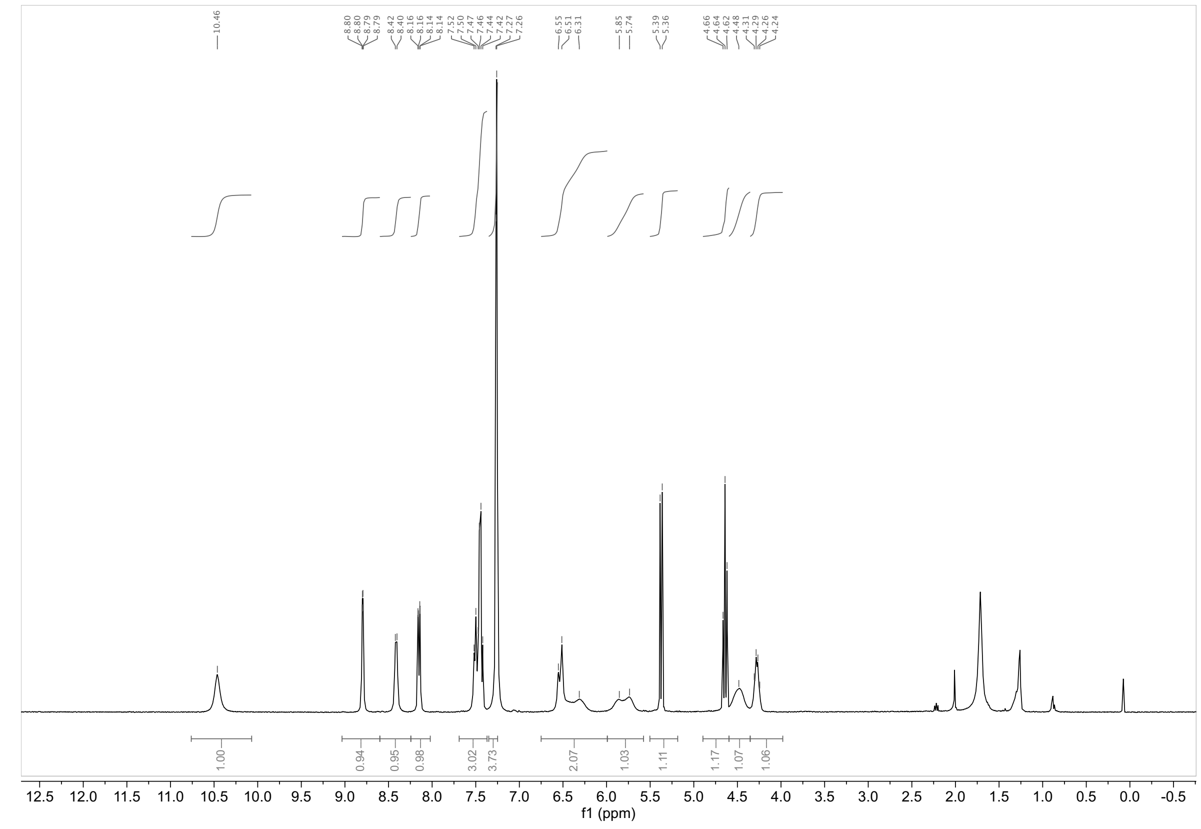

#### $^1$H NMR of MY-3A (3) in $CDCl_3$

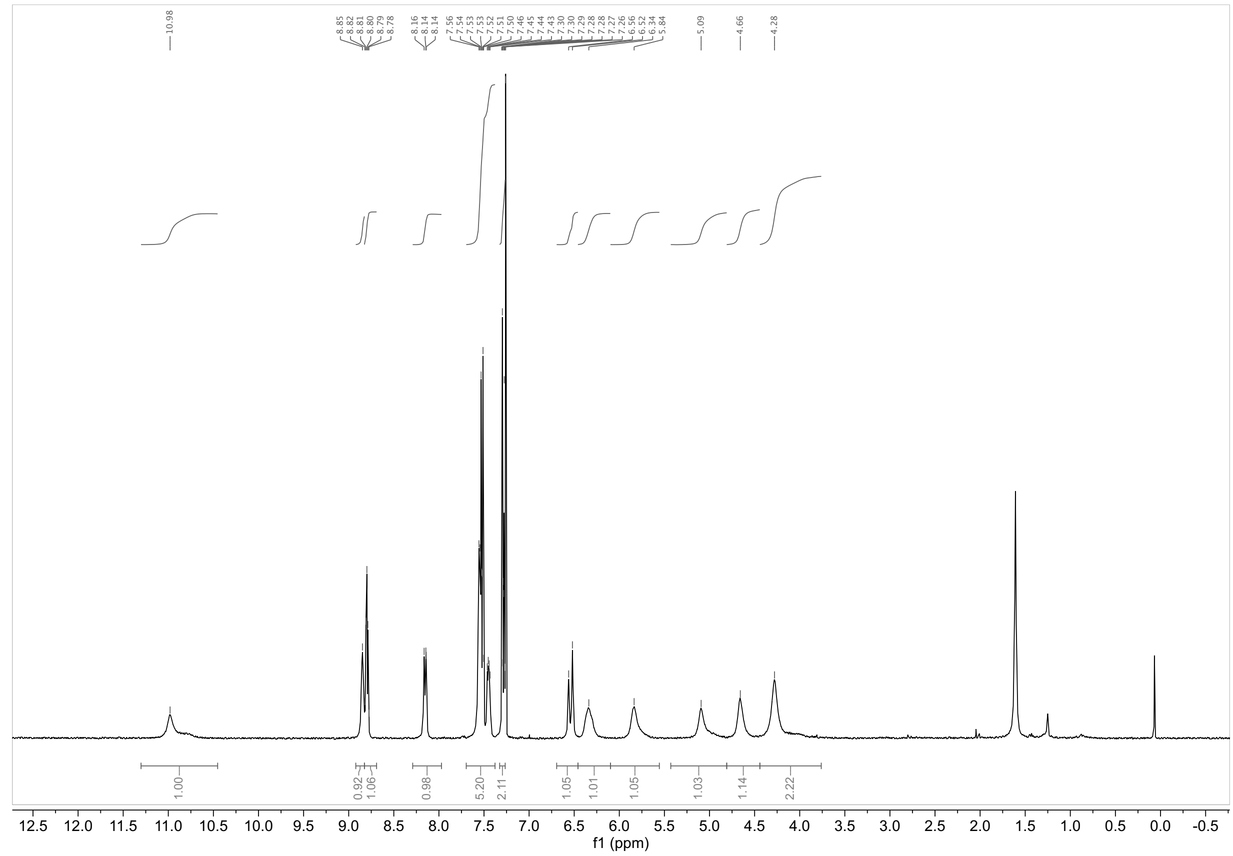

#### $^1$H NMR of MY-3B (4) in $CDCl_3$

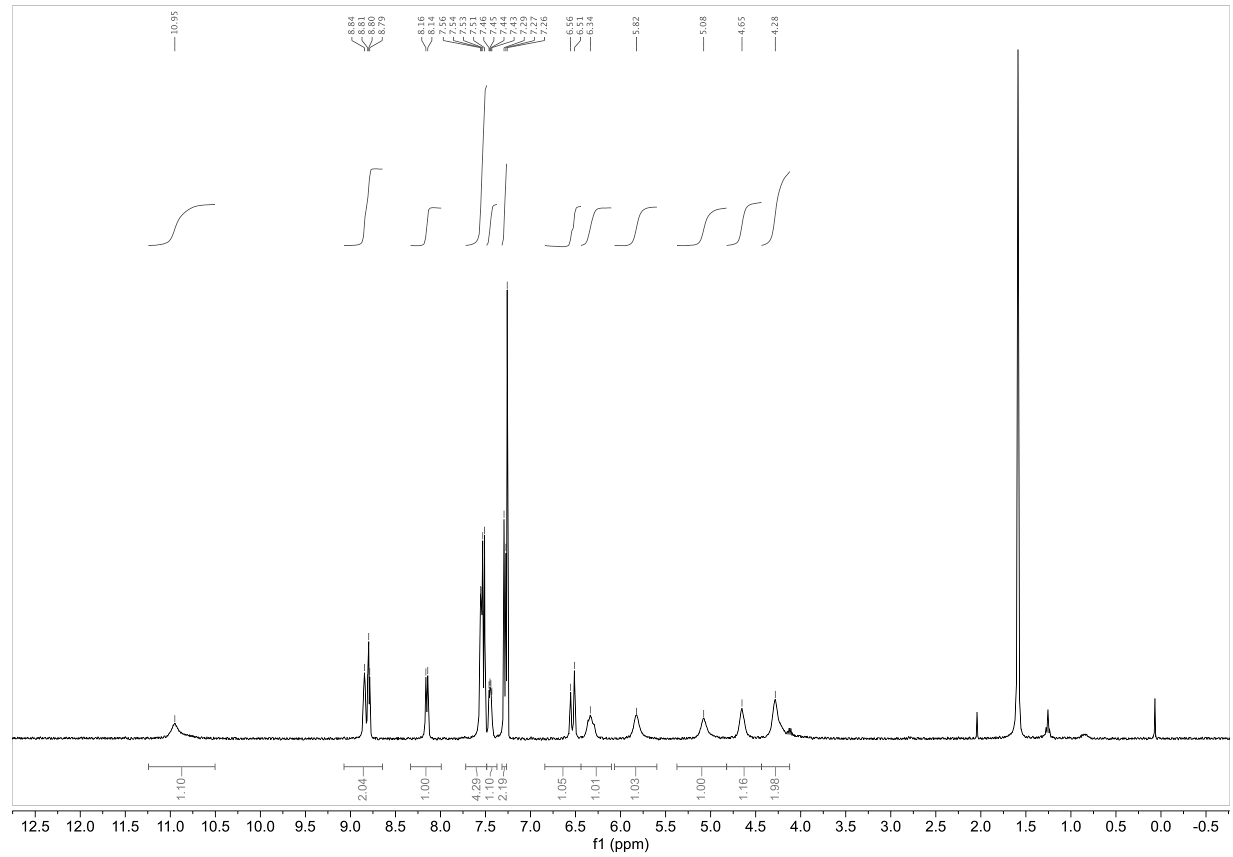

**[1]H NMR of EV-96 (5) in $CD_3OD$**

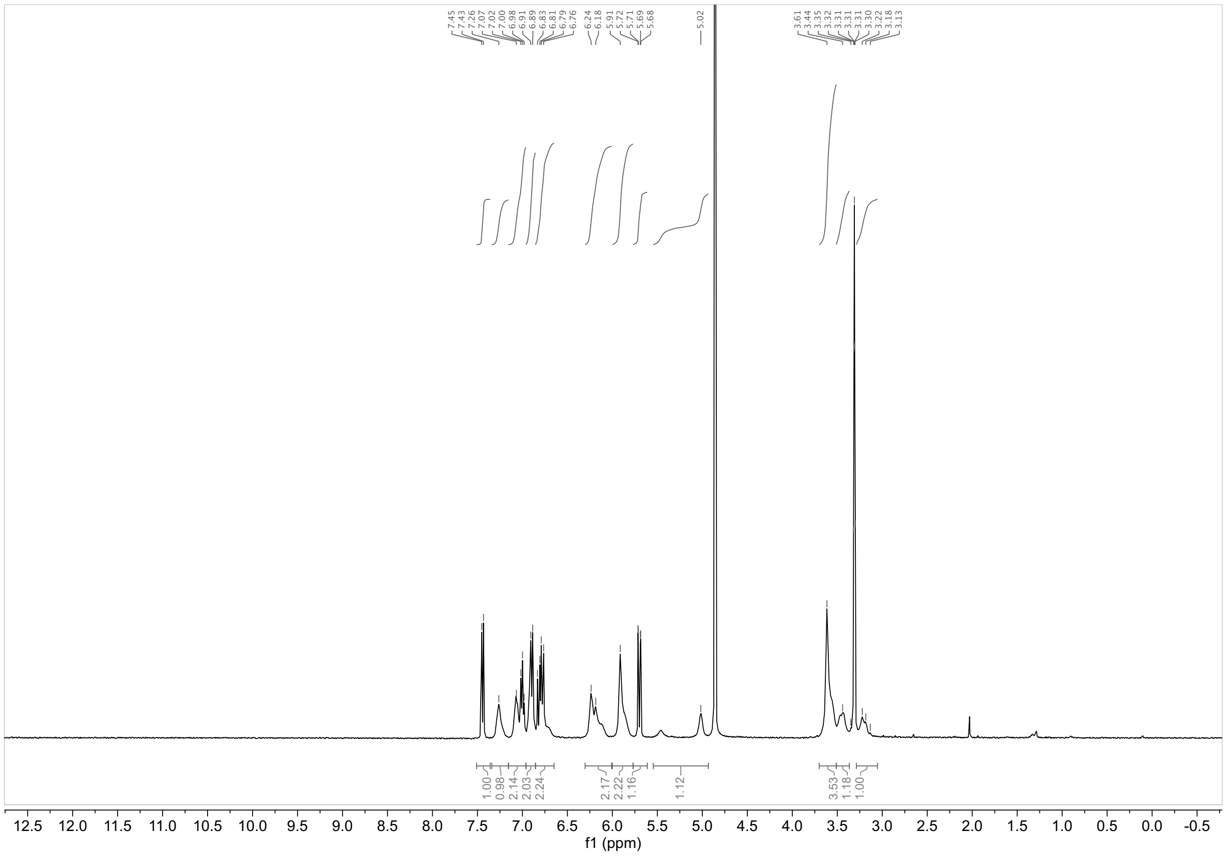

**[1]H NMR of EV-97 (6) in $CD_3OD$**

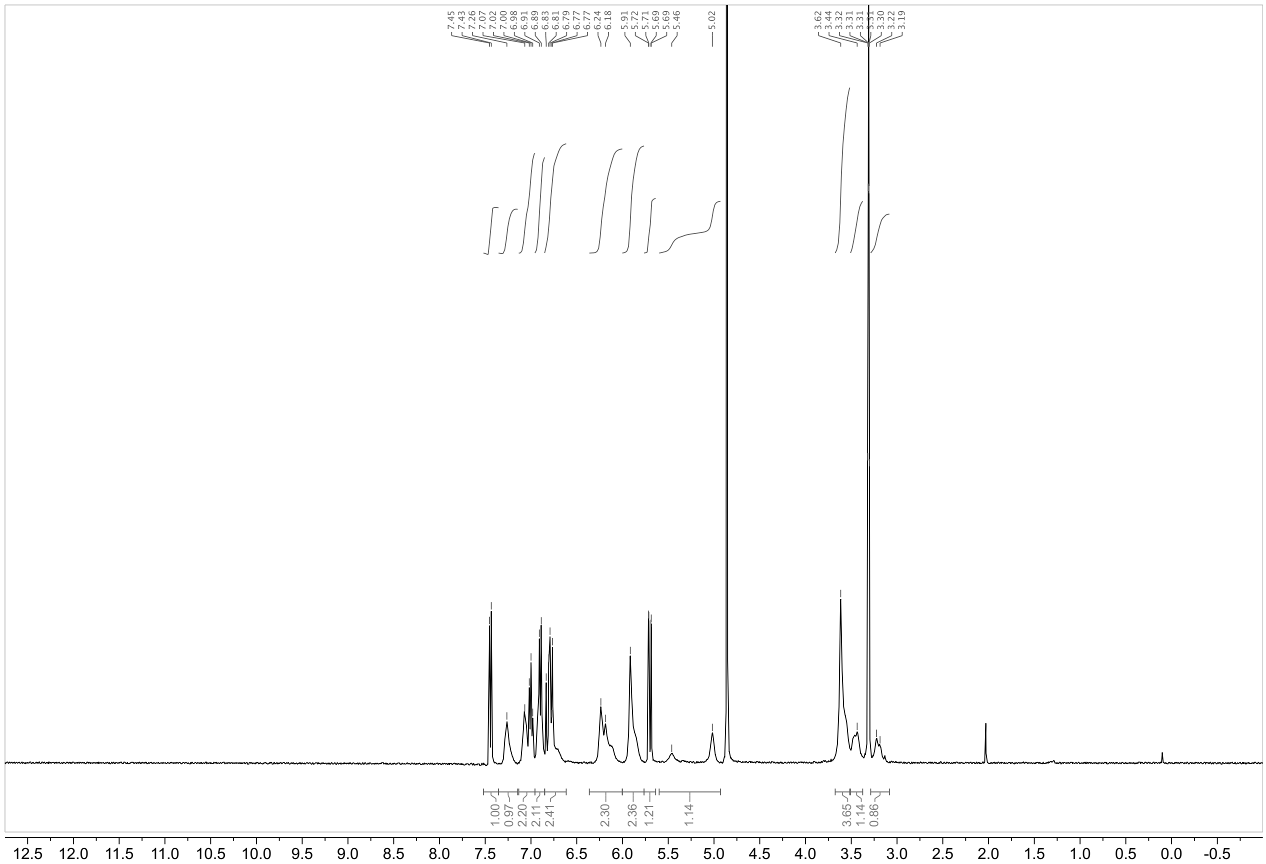

**$^{1}H$ NMR of EV-98 (7) in $CD_3OD$**

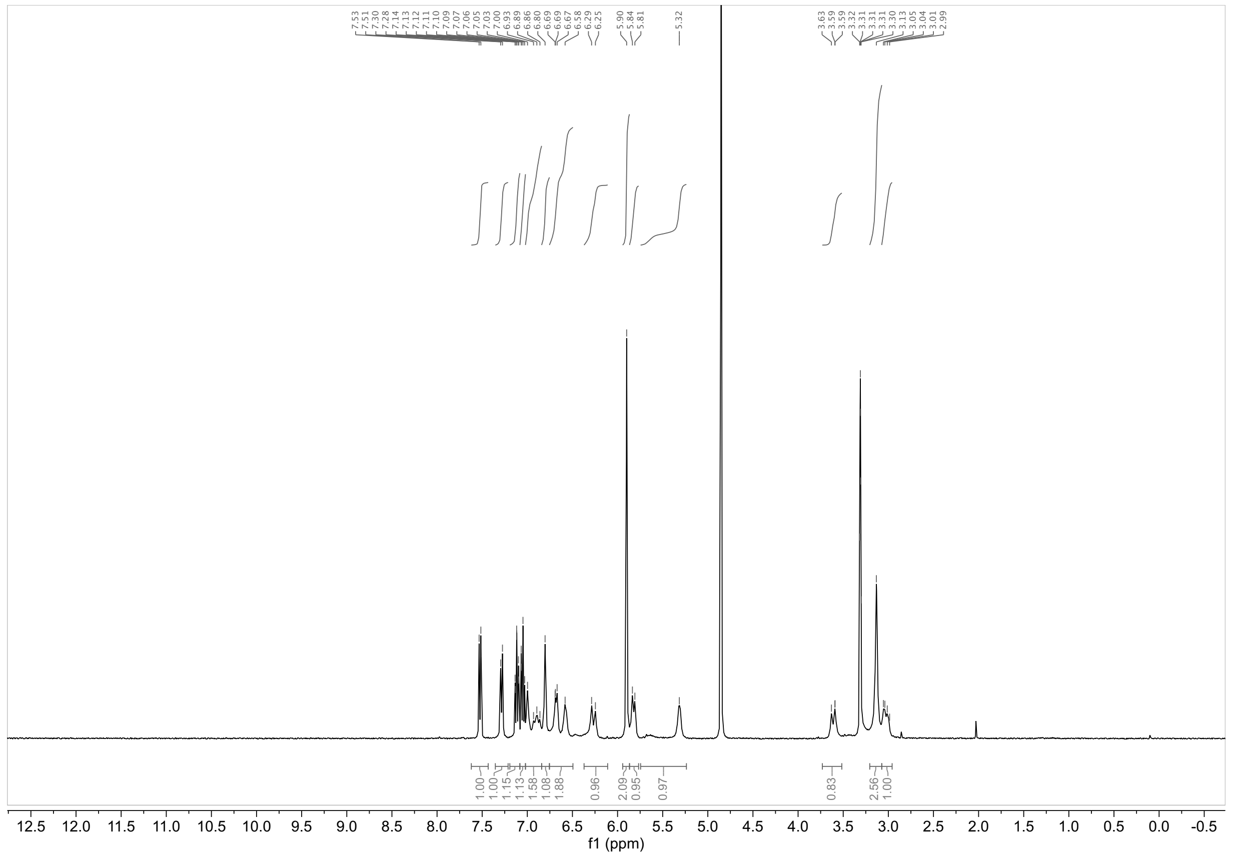

**$^{1}H$ NMR of EV-99 (8) in $CD_3OD$**

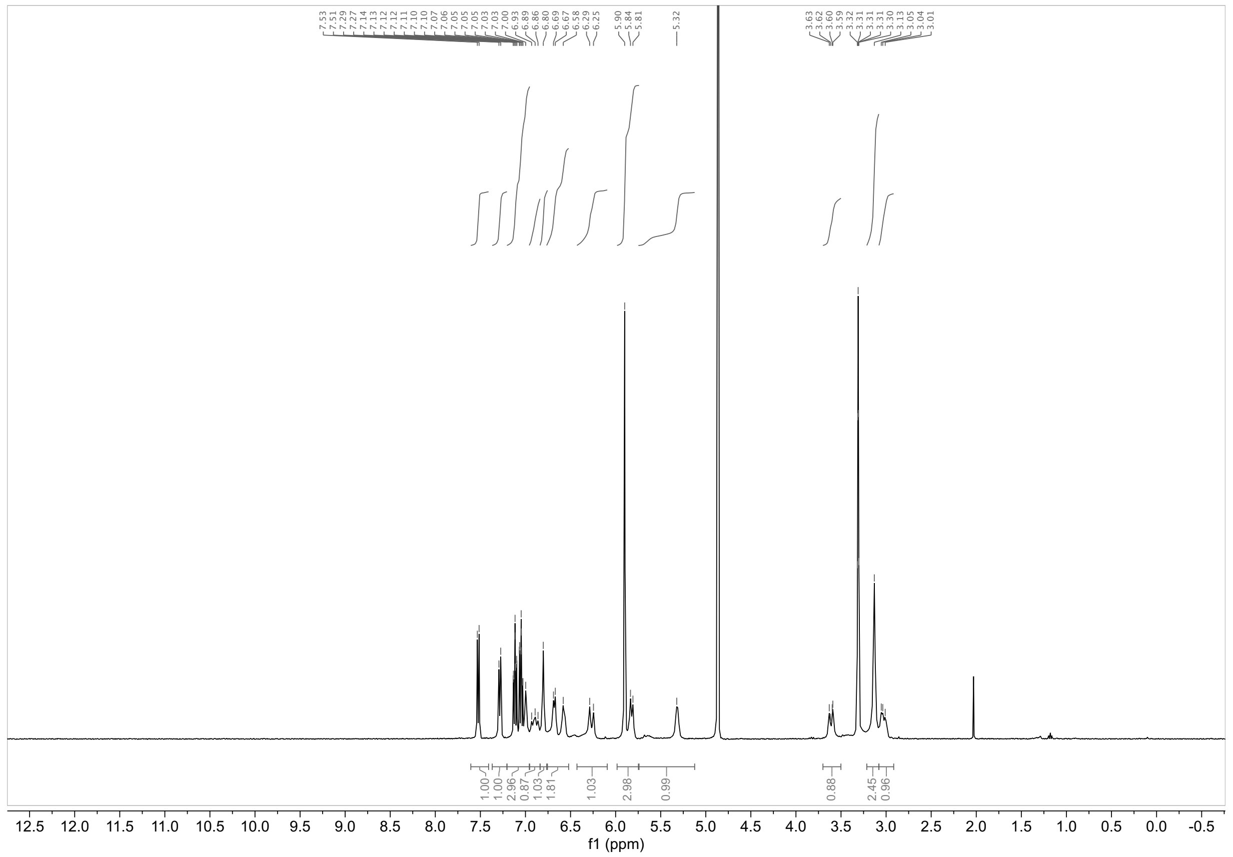

#### $^1$H NMR of MY-11A (9) in $CD_3OD$

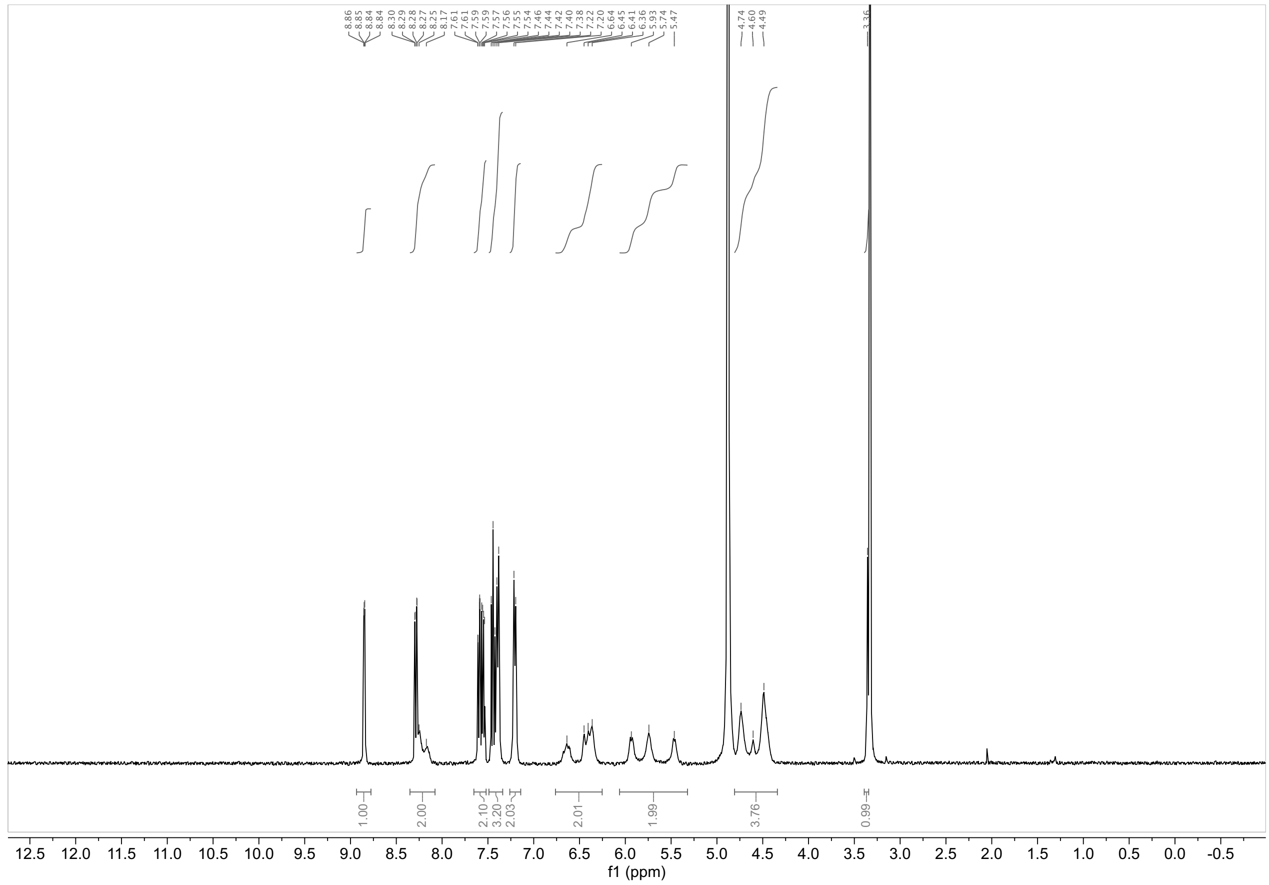

#### $^1$H NMR of MY-11B (10) in DMSO-$d_6$

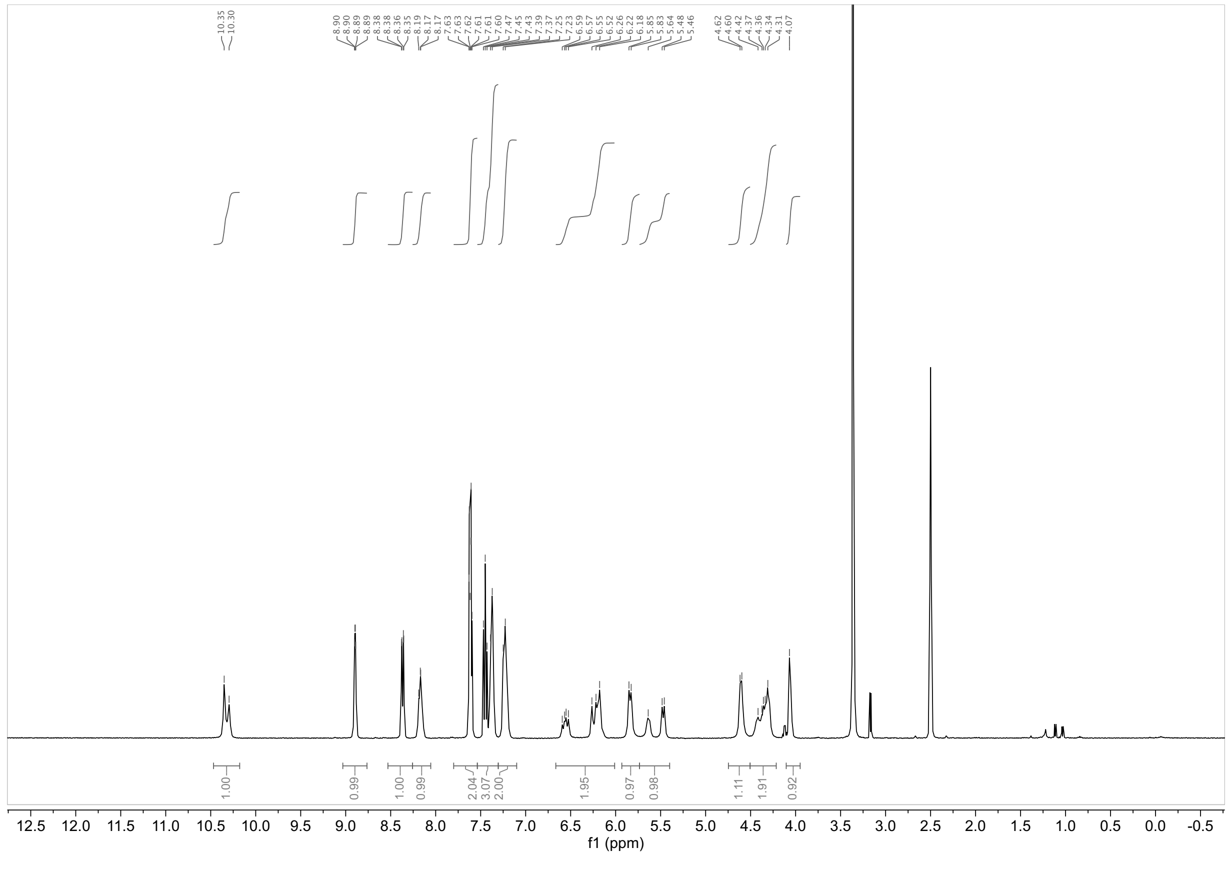

**$^1$H NMR of MY-45A (11) in $CDCl_3$**

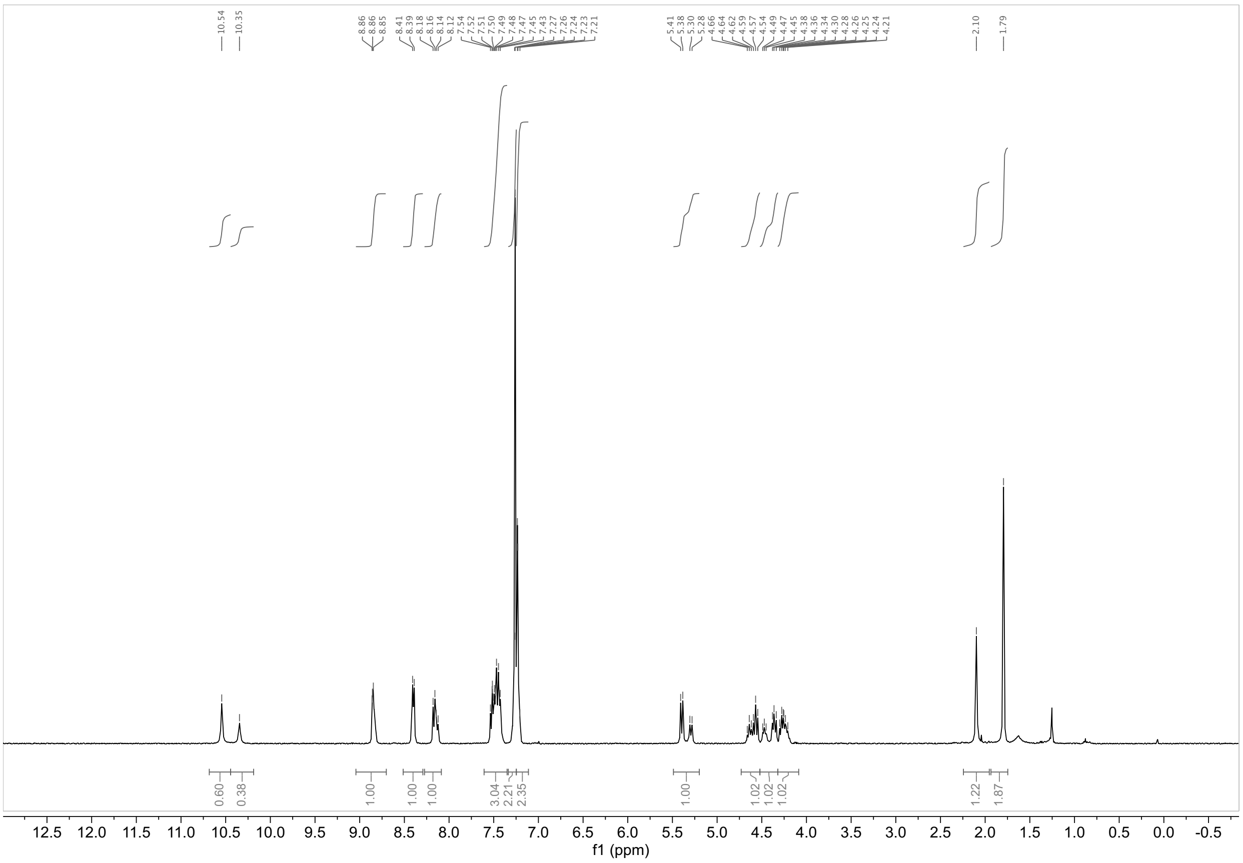

**$^1$H NMR of MY-45B (12) in $CDCl_3$**

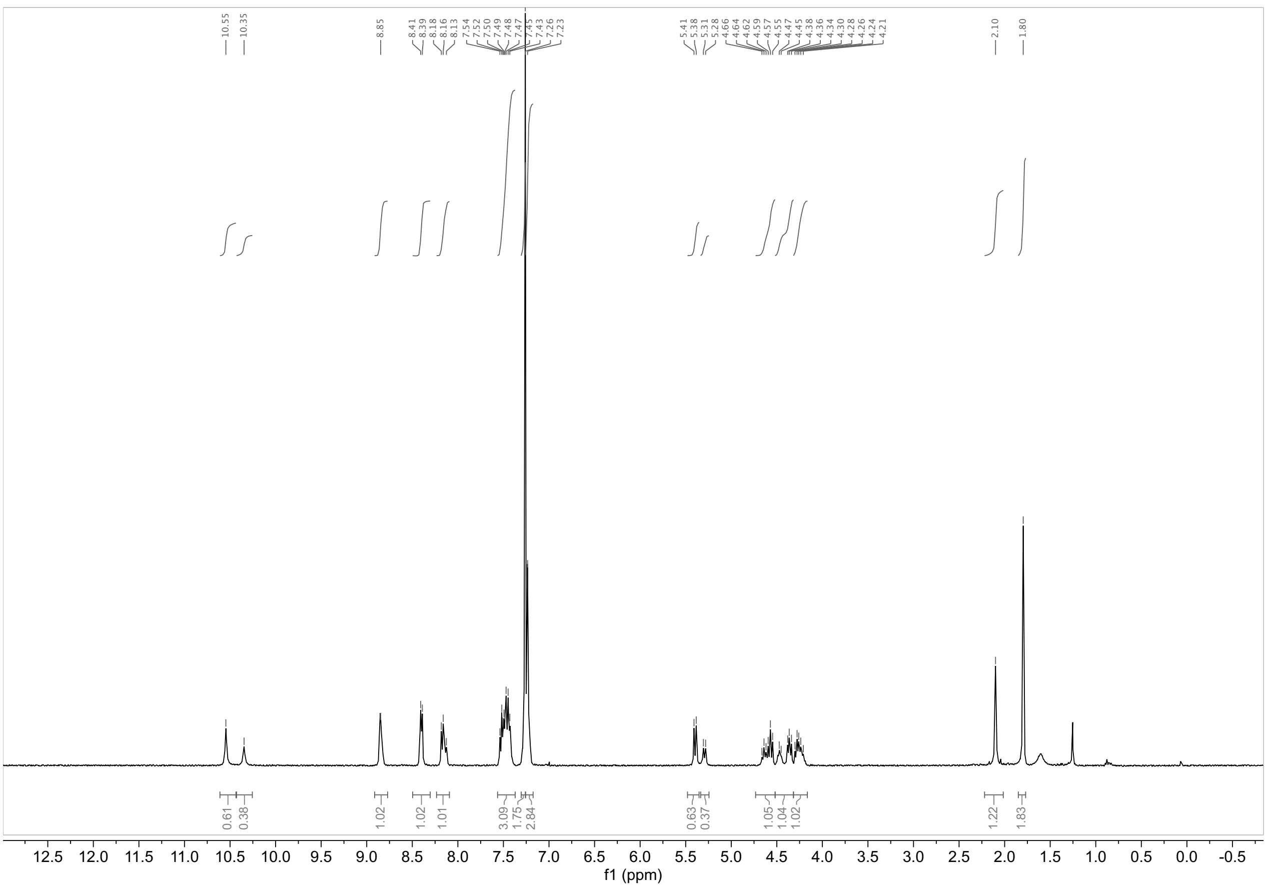

##### $^{1}H$ NMR of WX-01-10 (13) in $CDCl_3$

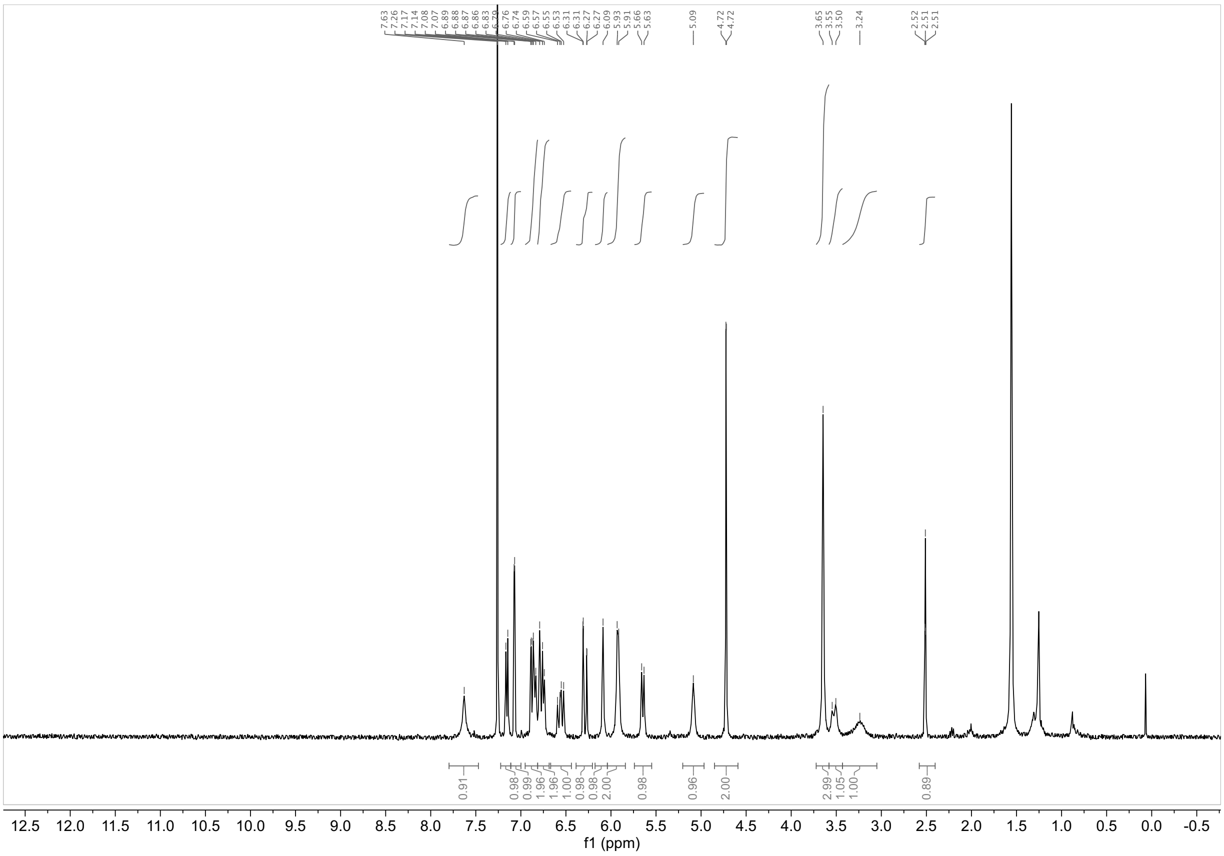

##### $^{1}H$ NMR of WX-01-12 (14) in $CDCl_3$

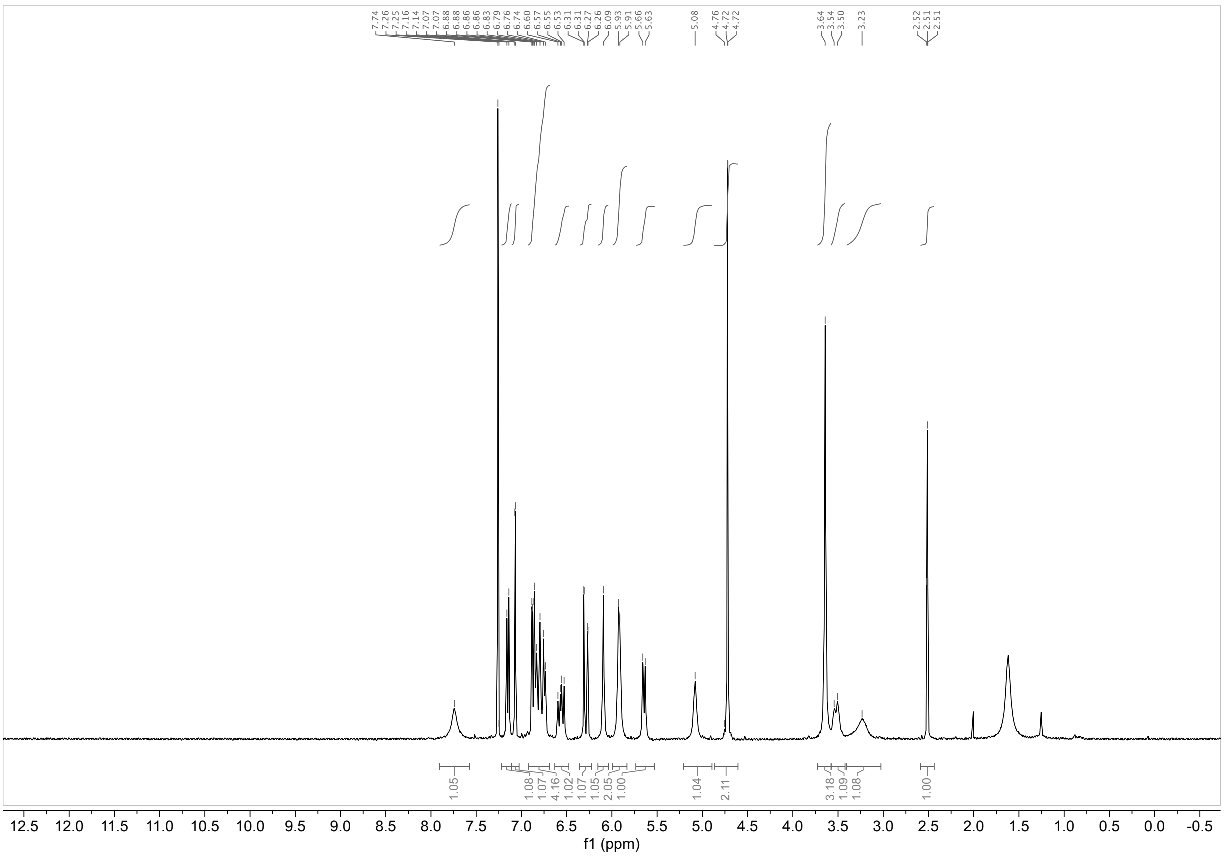

#### $^1H$ NMR of WX-02-23 (15) in $CD_3OD$

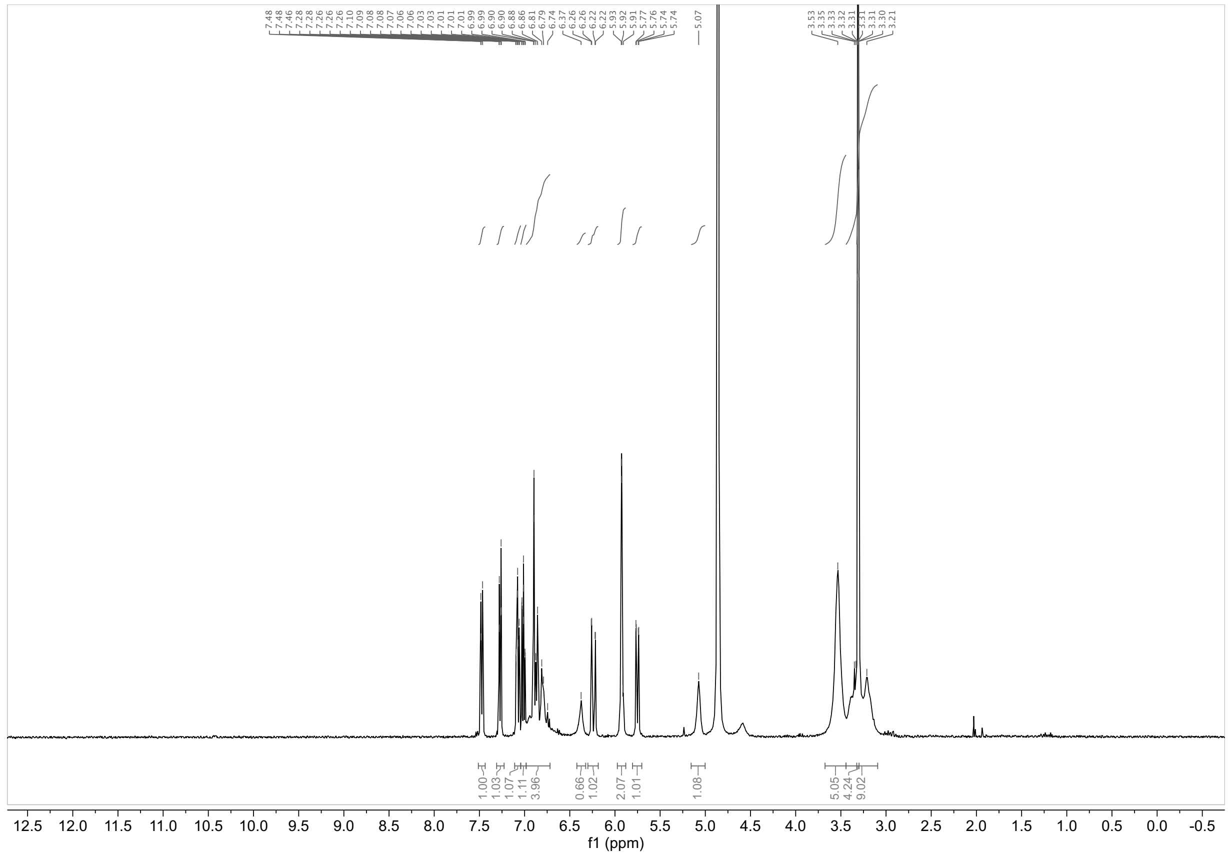

#### $^1H$ NMR of WX-02-43 (16) in $CD_3OD$

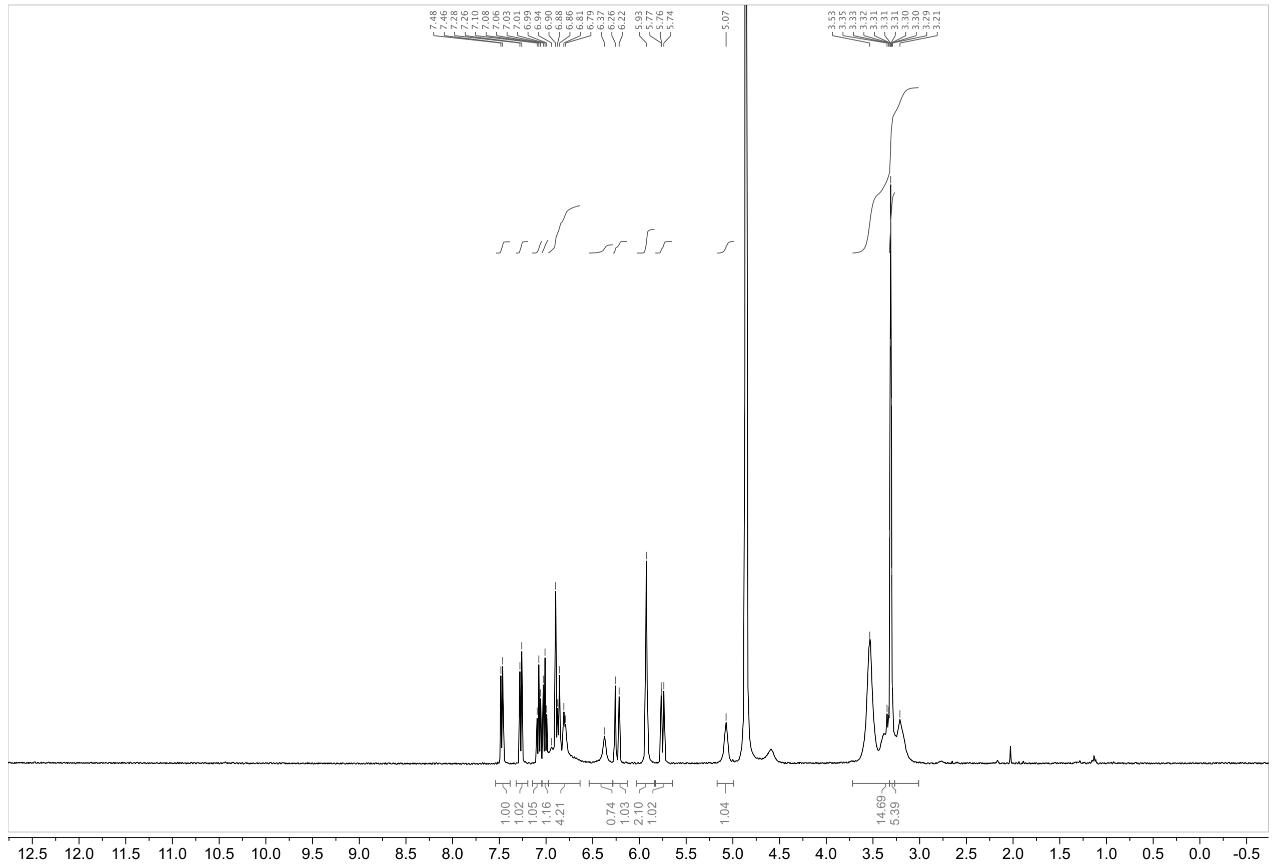

**$^1$H NMR of EV-96-ctrl (17) in $CD_3OD$**

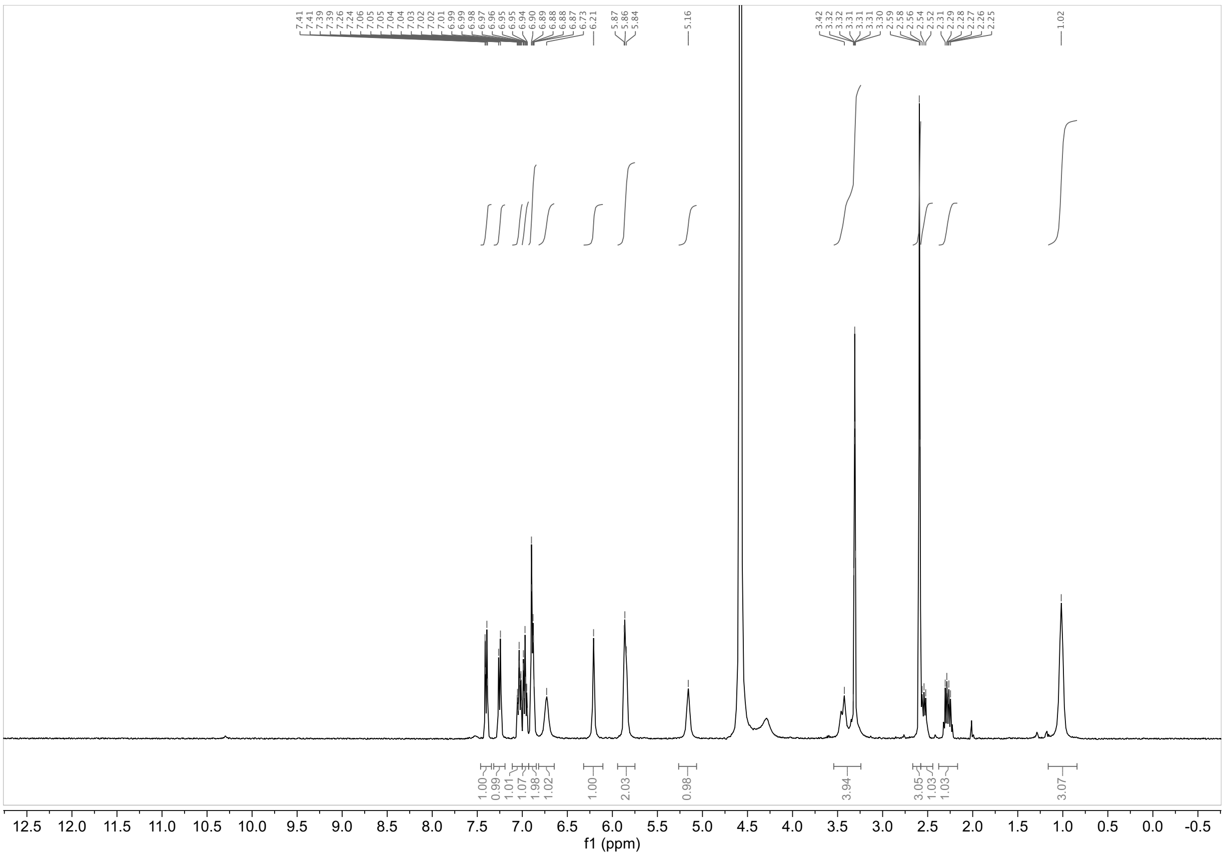

**$^1$H NMR of EV-97-ctrl (18) in $CD_3OD$**

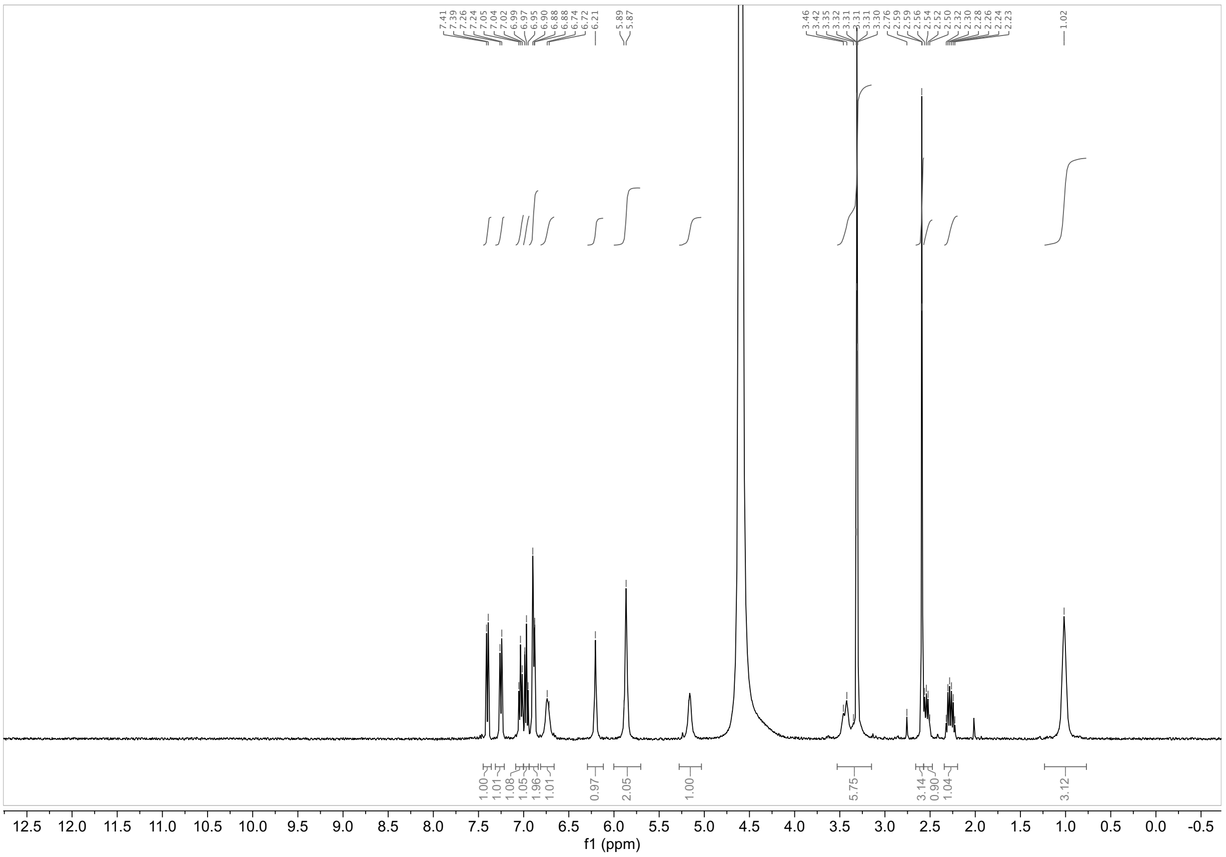

#### [1]H NMR of EV-98-ctrl (19) in $CD_3OD$

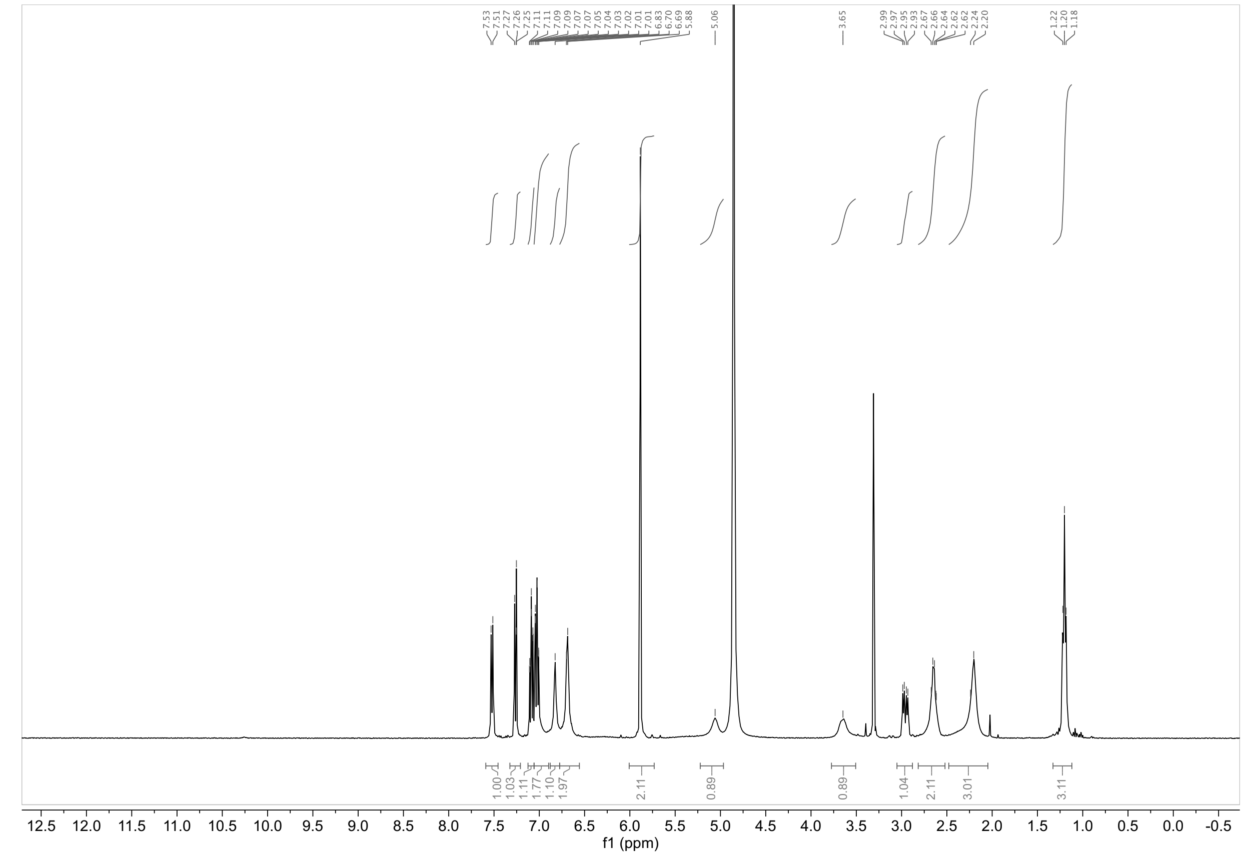

#### [1]H NMR of EV-99-ctrl (20) in $CD_3OD$

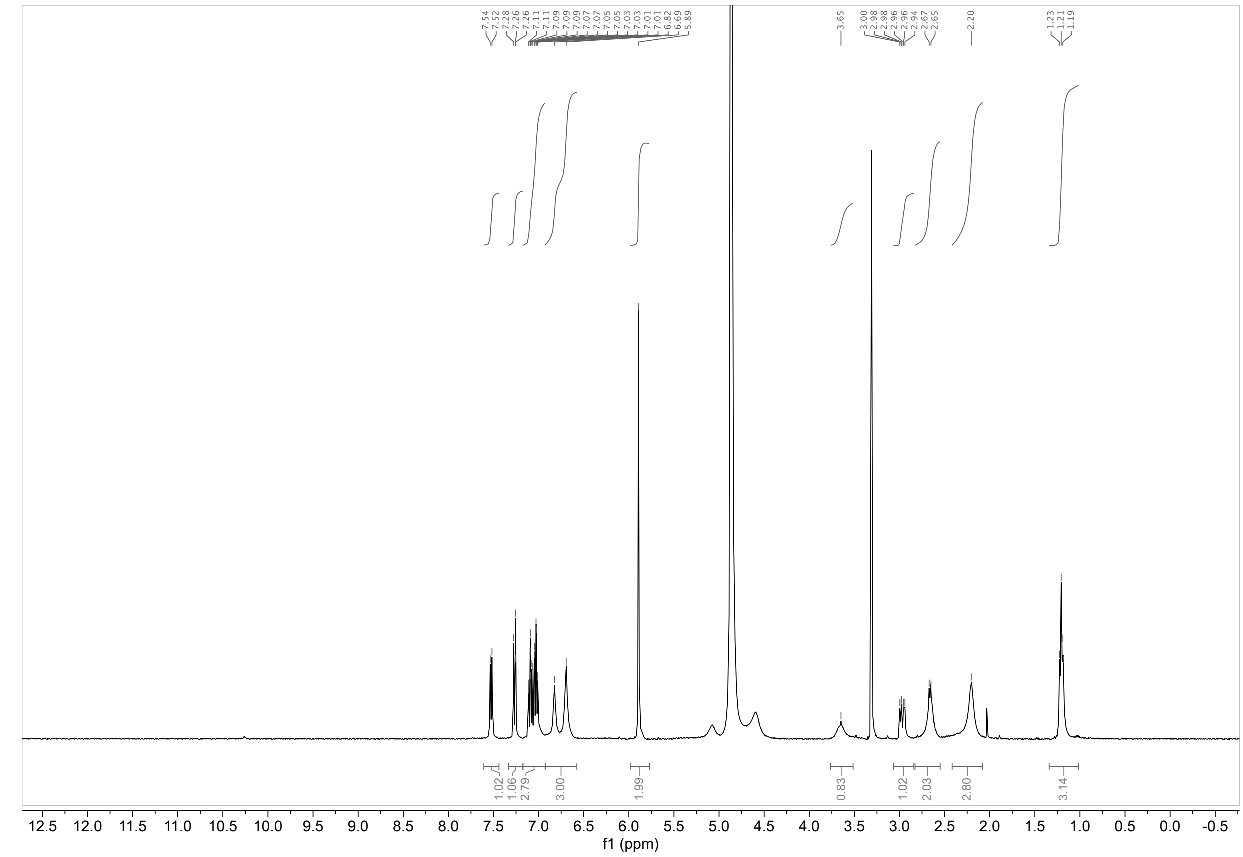

**Chiral stationary phase SFC:**

**MY-1A (1)**

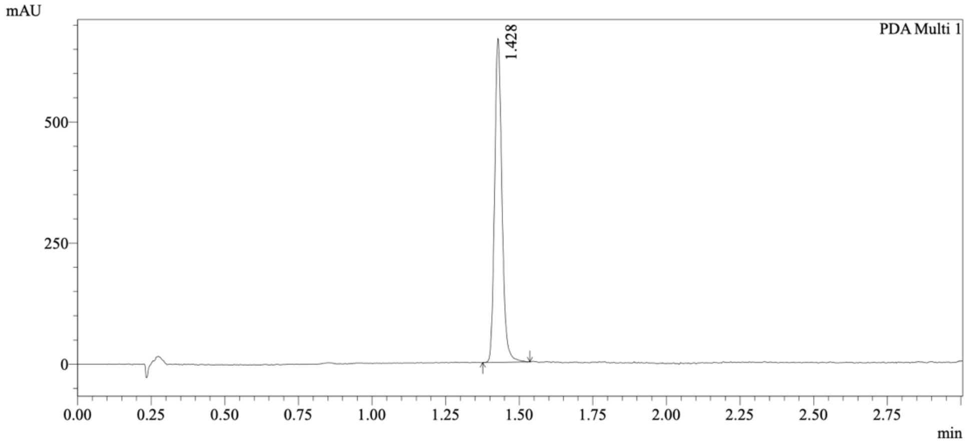

1 PDA Multi 1 / 220nm,4nm

Integration Results

PeakTable

PDA Ch1 220nm

| Peak# | Ret. Time | USP Width | Resolution | Height | Area | Area % |
|---|---|---|---|---|---|---|
| 1 | 1.428 | 0.049 | 0.000 | 652902 | 1161646 | 100.000 |
| Total | | | | 652902 | 1161646 | 100.000 |

**MY-1B (2)**

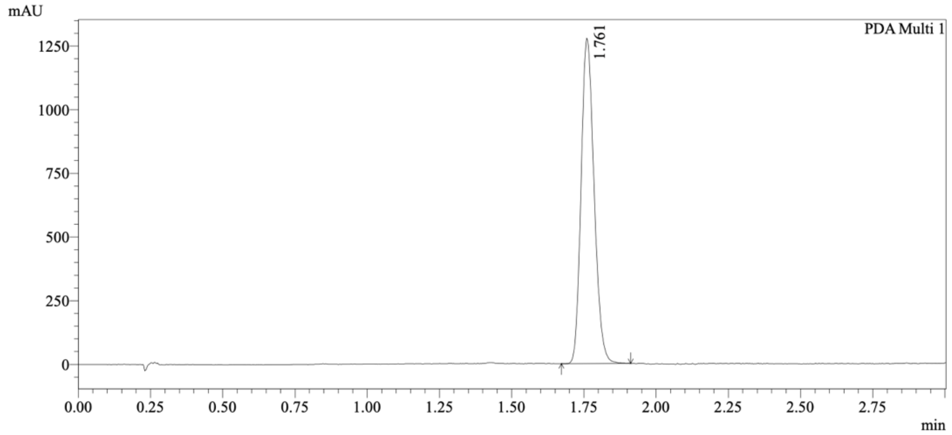

1 PDA Multi 1 / 220nm,4nm

Integration Results

PeakTable

PDA Ch1 220nm

| Peak# | Ret. Time | USP Width | Resolution | Height | Area | Area % |
|---|---|---|---|---|---|---|
| 1 | 1.761 | 0.085 | 0.000 | 1254375 | 4001669 | 100.000 |
| Total | | | | 1254375 | 4001669 | 100.000 |

**MY-1A and MY-1B mixture**

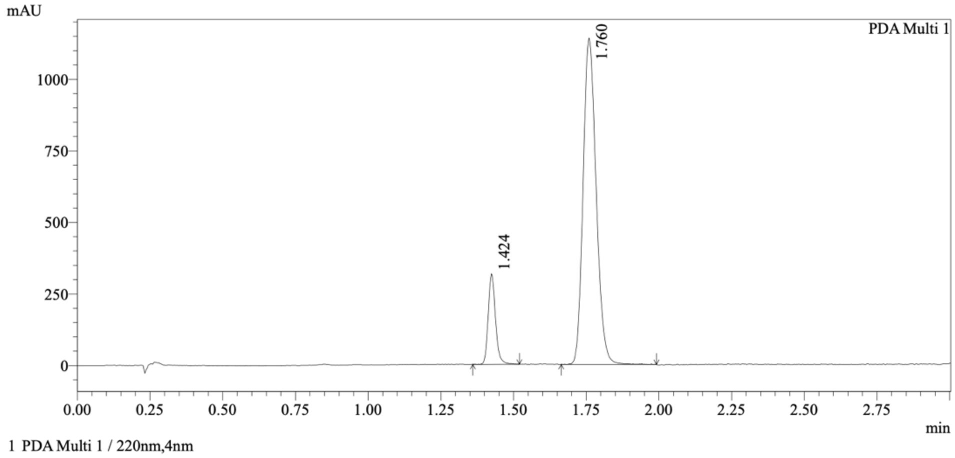

Integration Results

PeakTable

PDA Ch1 220nm

| Peak# | Ret. Time | USP Width | Resolution | Height | Area | Area % |
|---|---|---|---|---|---|---|
| 1 | 1.424 | 0.049 | 0.000 | 296527 | 545869 | 13.295 |
| 2 | 1.760 | 0.085 | 5.021 | 1125780 | 3560048 | 86.705 |
| Total | | | | 1422307 | 4105918 | 100.000 |

*Method*

Column: Chiralcel OJ-3 50×4.6mm I.D., 3 μm;

Mobile phase B: MeOH (0.05% DEA);

Gradient elution: 5% to 40% B.

**MY-3A (3)**

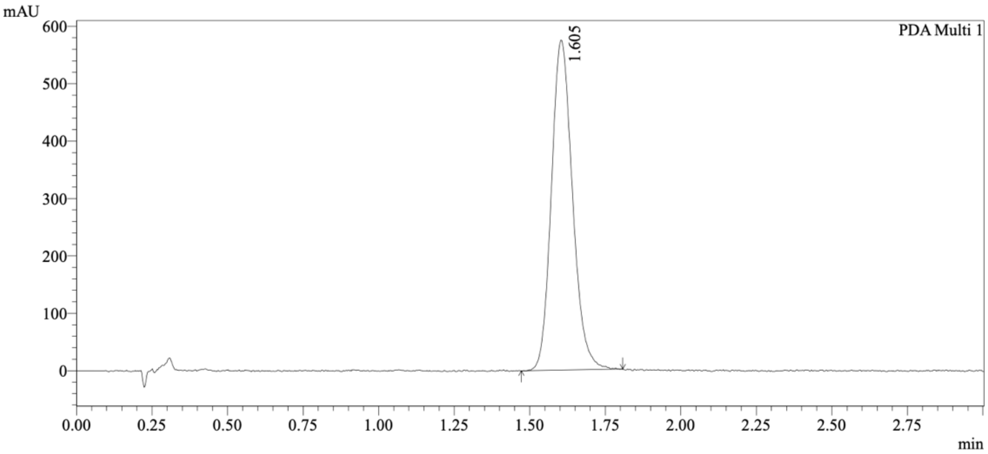

1 PDA Multi 1 / 220nm,4nm

Integration Results

PeakTable

PDA Ch1 220nm

| Peak# | Ret. Time | USP Width | Resolution | Height | Area | Area % |
|---|---|---|---|---|---|---|
| 1 | 1.605 | 0.129 | 0.000 | 561301 | 2805243 | 100.000 |
| Total | | | | 561301 | 2805243 | 100.000 |

**MY-3B (4)**

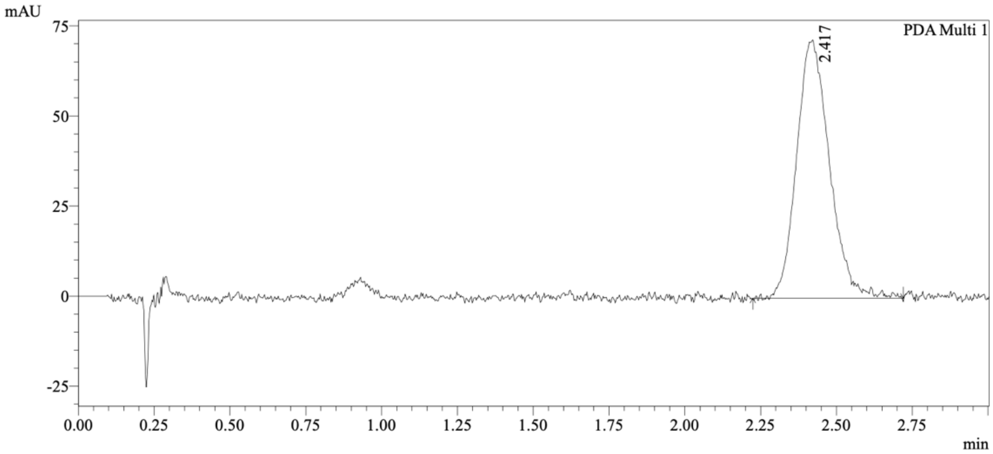

1 PDA Multi 1 / 220nm,4nm

Integration Results

PeakTable

PDA Ch1 220nm

| Peak# | Ret. Time | USP Width | Resolution | Height | Area | Area % |
|---|---|---|---|---|---|---|
| 1 | 2.417 | 0.197 | 0.000 | 70006 | 540538 | 100.000 |
| Total | | | | 70006 | 540538 | 100.000 |

**MY-3A and MY-3B mixture**

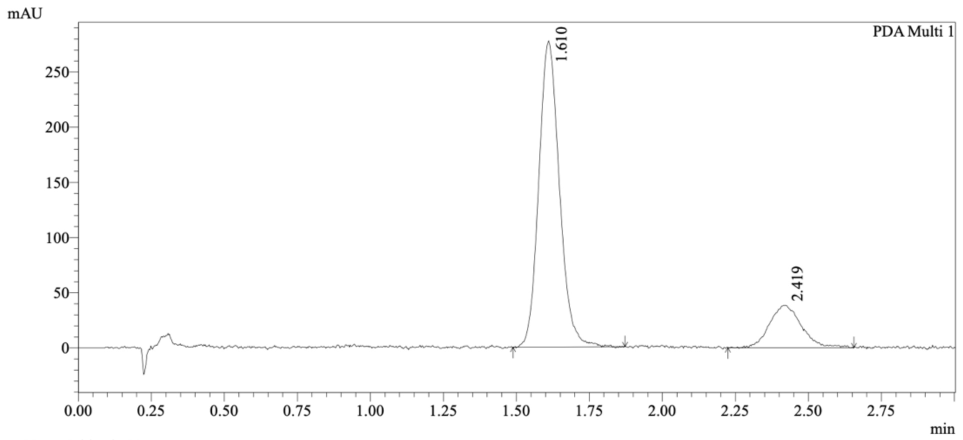

1 PDA Multi 1 / 220nm,4nm

Integration Results

PeakTable

PDA Ch1 220nm

| Peak# | Ret. Time | USP Width | Resolution | Height | Area | Area % |
|---|---|---|---|---|---|---|
| 1 | 1.610 | 0.131 | 0.000 | 273503 | 1365053 | 82.414 |
| 2 | 2.419 | 0.197 | 4.933 | 37483 | 291274 | 17.586 |
| Total | | | | 310986 | 1656326 | 100.000 |

*Method*

Column: Chiralpak AD-3 50×4.6mm I.D., 3 μm;

Mobile phase: A: $CO_2$, B: iPrOH (0.05% DEA);

Gradient elution: 40% B (isocratic).

**EV-96 (5)**

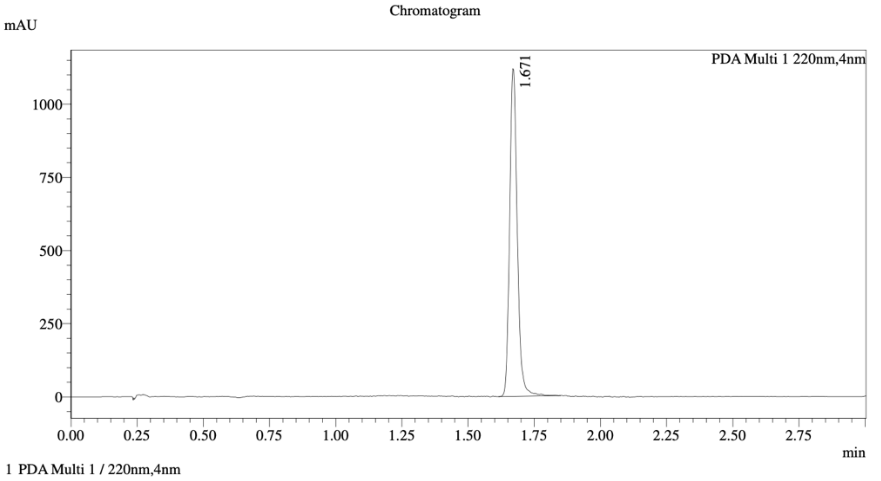

Integration Result

Peak Table

PDA Ch1 220nm

| Peak# | Ret. Time | Height | Height% | Resolution(USP) | Area | Area% |
|---|---|---|---|---|---|---|
| 1 | 1.671 | 1103556 | 100.000 | -- | 2250054 | 100.000 |

**EV-97 (6)**

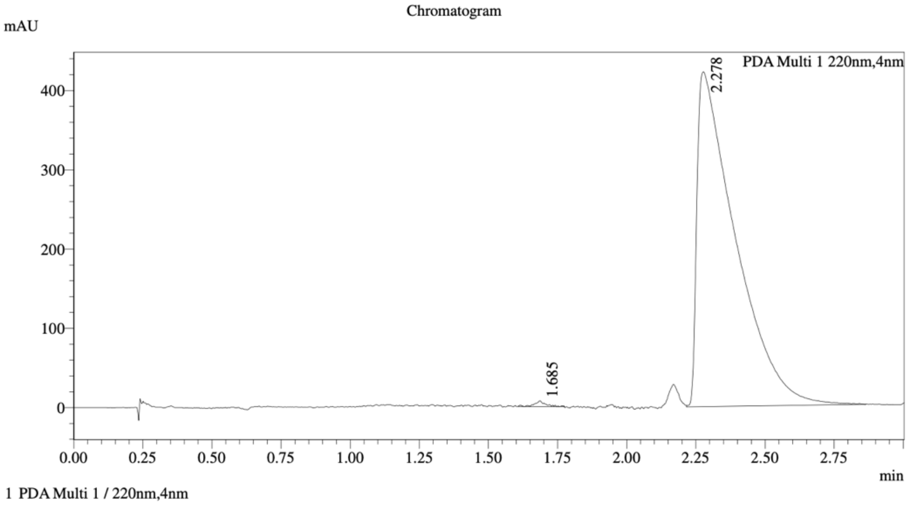

Integration Result

Peak Table

PDA Ch1 220nm

| Peak# | Ret. Time | Height | Height% | Resolution(USP) | Area | Area% |
|---|---|---|---|---|---|---|
| 1 | 1.685 | 6779 | 1.581 | -- | 18710 | 0.456 |
| 2 | 2.278 | 421874 | 98.419 | 3.818 | 4086960 | 99.544 |

**EV-96 and EV-97 mixture**

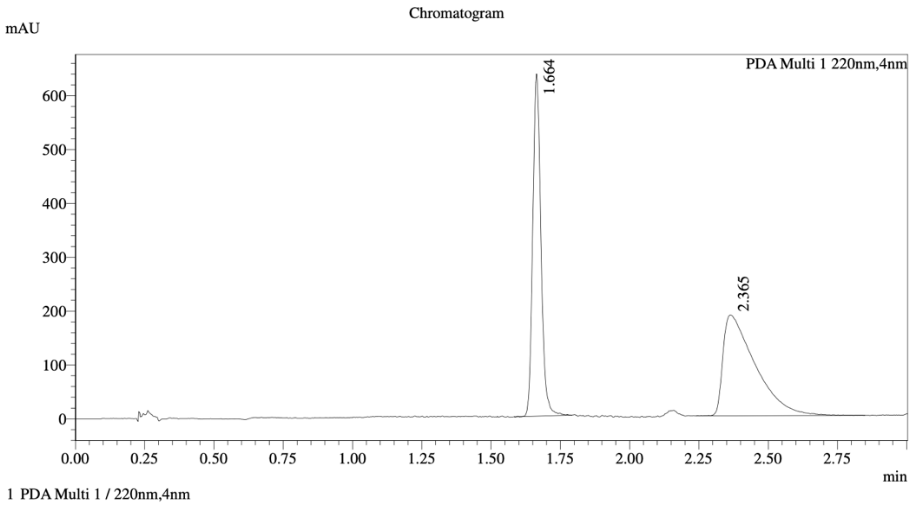

==========================================================================
Integration Result
==========================================================================

Peak Table

PDA Ch1 220nm

| Peak# | Ret. Time | Height | Height% | Resolution(USP) | Area | Area% |
|---|---|---|---|---|---|---|
| 1 | 1.664 | 610940 | 76.585 | -- | 1306910 | 46.379 |
| 2 | 2.365 | 186793 | 23.415 | 5.162 | 1511007 | 53.621 |

*Method*

Column: Chiralcel OD-3 50×4.6mm I.D., 3 μm;

Mobile phase B: MeOH (0.05% DEA);

Gradient elution: 5% to 40% B.

**EV-98 (7)**

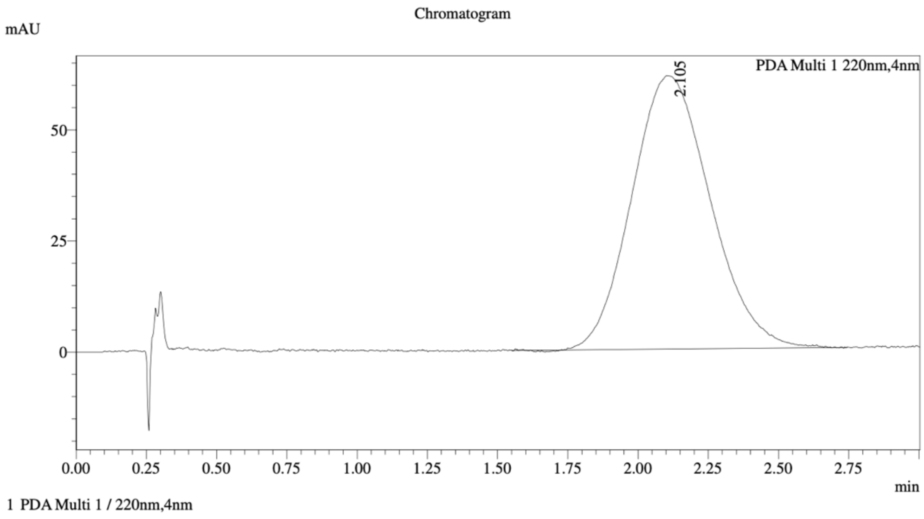

==================================================================================
Integration Result
==================================================================================

Peak Table

PDA Ch1 220nm

| Peak# | Ret. Time | Height | Height% | Resolution(USP) | Area | Area% |
|---|---|---|---|---|---|---|
| 1 | 2.105 | 61452 | 100.000 | -- | 1196846 | 100.000 |

**EV-99 (8)**

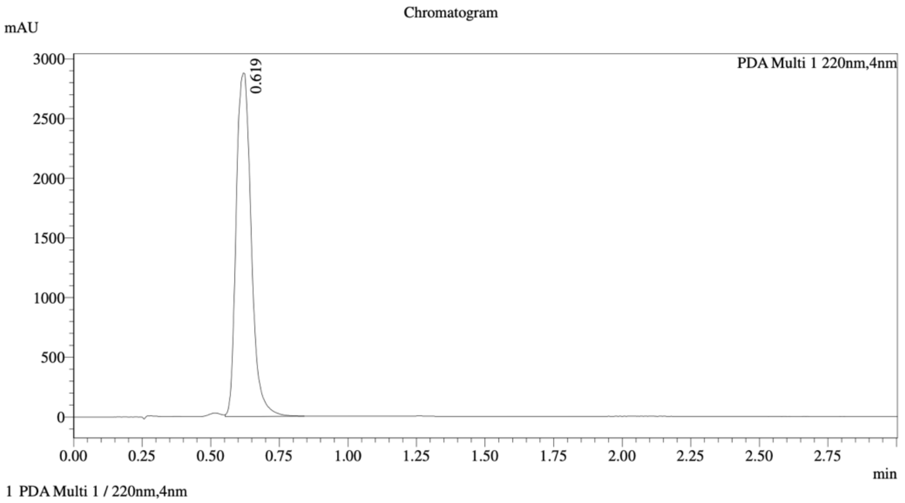

==================================================================================
Integration Result
==================================================================================

Peak Table

PDA Ch1 220nm

| Peak# | Ret. Time | Height | Height% | Resolution(USP) | Area | Area% |
|---|---|---|---|---|---|---|
| 1 | 0.619 | 2874854 | 100.000 | -- | 10981838 | 100.000 |

**EV-98 and EV-99 mixture**

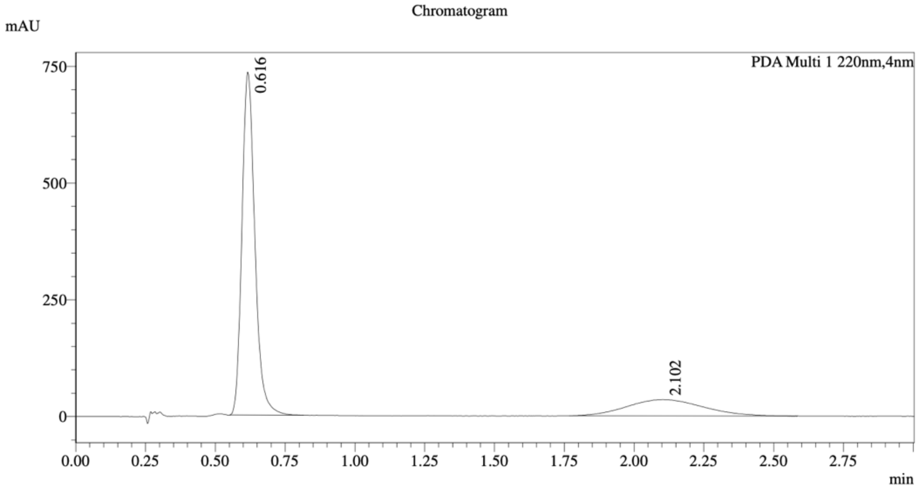

1 PDA Multi 1 / 220nm,4nm

================================================================================
Integration Result
================================================================================

Peak Table

PDA Ch1 220nm

| Peak# | Ret. Time | Height | Height% | Resolution(USP) | Area | Area% |
|---|---|---|---|---|---|---|
| 1 | 0.616 | 722077 | 95.396 | -- | 2366200 | 78.122 |
| 2 | 2.102 | 34845 | 4.604 | 5.013 | 662663 | 21.878 |

*Method*

Column: Chiralpak AD-3 50×4.6mm I.D., 3 μm

Mobile phase B: MeOH/$CH_3CN$ (0.05% DEA);

Gradient elution: 40% B (isocratic)

**MY-11A (9)**

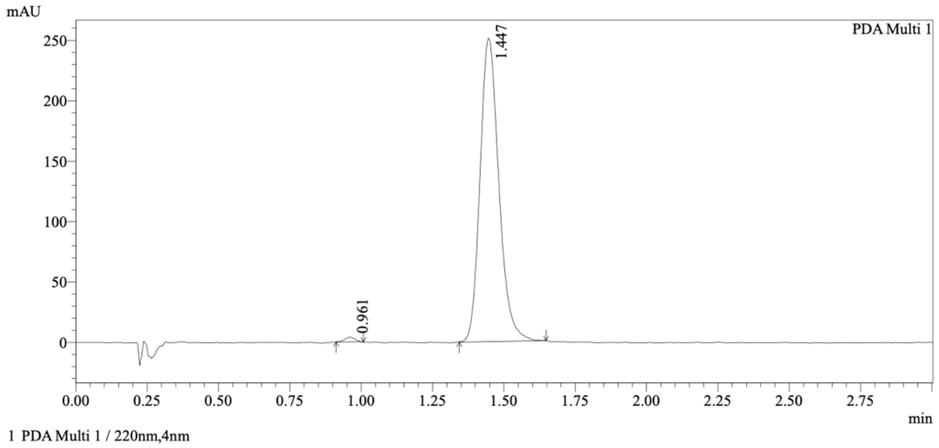

Integration Results

PeakTable

PDA Ch1 220nm

| Peak# | Ret. Time | USP Width | Resolution | Height | Area | Area % |
|---|---|---|---|---|---|---|
| 1 | 0.961 | 0.072 | 0.000 | 3299 | 9386 | 0.809 |
| 2 | 1.447 | 0.120 | 5.055 | 247033 | 1151490 | 99.191 |
| Total | | | | 250333 | 1160876 | 100.000 |

**MY-11B (10)**

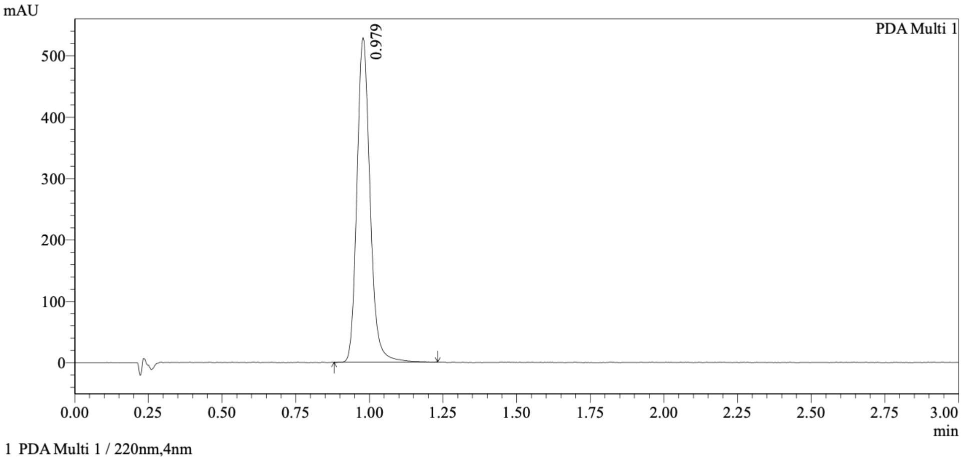

Integration Results

PeakTable

PDA Ch1 220nm

| Peak# | Ret. Time | USP Width | Resolution | Height | Area | Area % |
|---|---|---|---|---|---|---|
| 1 | 0.979 | 0.082 | 0.000 | 487407 | 1637634 | 100.000 |
| Total | | | | 487407 | 1637634 | 100.000 |

**MY-11A and MY-11B mixture**

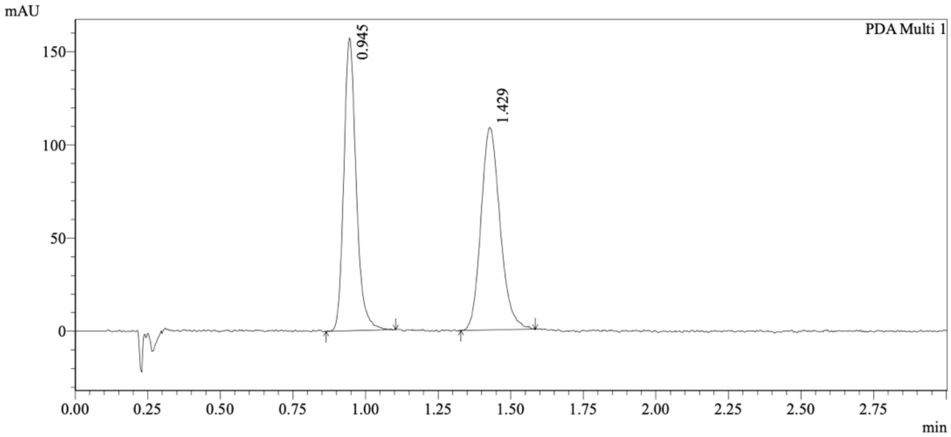

1 PDA Multi 1 / 220nm,4nm

Integration Results

PeakTable

PDA Ch1 220nm

| Peak# | Ret. Time | USP Width | Resolution | Height | Area | Area % |
|---|---|---|---|---|---|---|
| 1 | 0.945 | 0.079 | 0.000 | 140777 | 466791 | 48.372 |
| 2 | 1.429 | 0.121 | 4.848 | 105702 | 498217 | 51.628 |
| Total | | | | 246479 | 965007 | 100.000 |

*Method*

Column: Chiralpak AD-3 50×4.6mm I.D., 3 µm

Mobile phase: A: $CO_2$, B: iPrOH (0.05% DEA);

Gradient elution: 40% B (isocratic).

**MY-45A (11)**

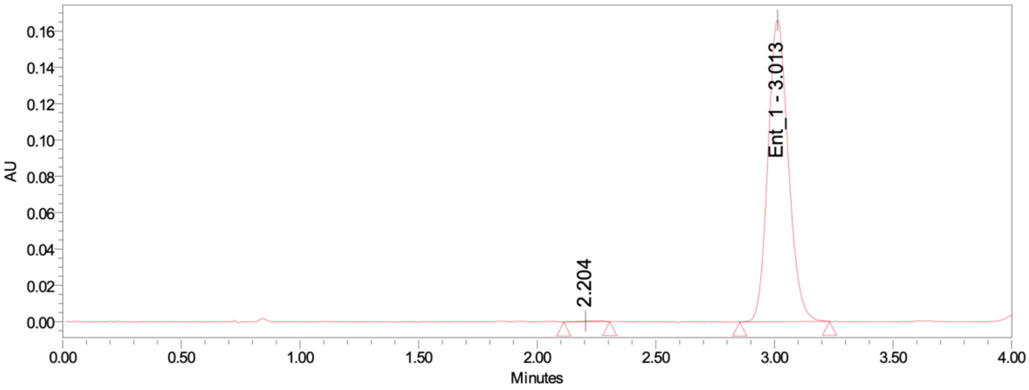

**MY-45B (12)**

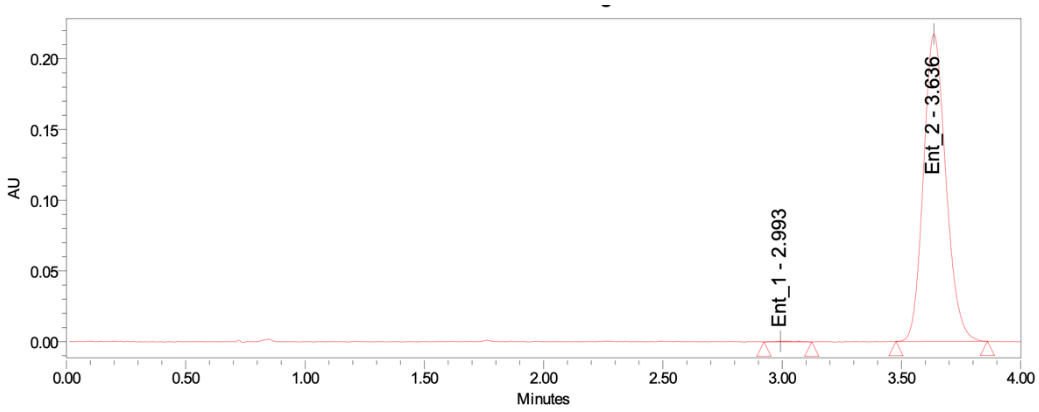

**MY-45A and MY-45B mixture**

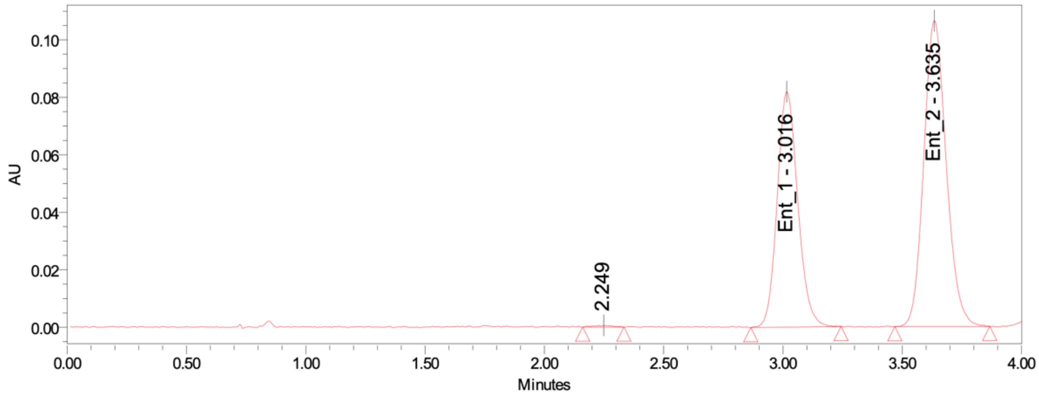

| | Peak 1 Area ($t_R$ 3.0 min) | | Peak 2 Area ($t_R$ 3.6 min) | | |
|---|---|---|---|---|---|
| **sample** | relative (%) | absolute | relative (%) | absolute | ee (%) |
| MY-45A | 100.00 | 991600 | - | - | 100.00 |
| MY-45B | 0.26 | 3707 | 99.74 | 1423004 | -99.48 |
| mixture | 41.32 | 491070 | 58.68 | 697307 | -17.35 |

*Method*

Column: Daicel IBN 250×4.6mm I.D., 3 μm

Column temperature:

Mobile phase: A: $CO_2$, B: MeOH

Gradient elution: 40% B (isocratic)

Flow rate: 3.5 mL / min

Back pressure: 110.3 bar

Detection wavelength: 235 nm

**WX-01-10 (13)**

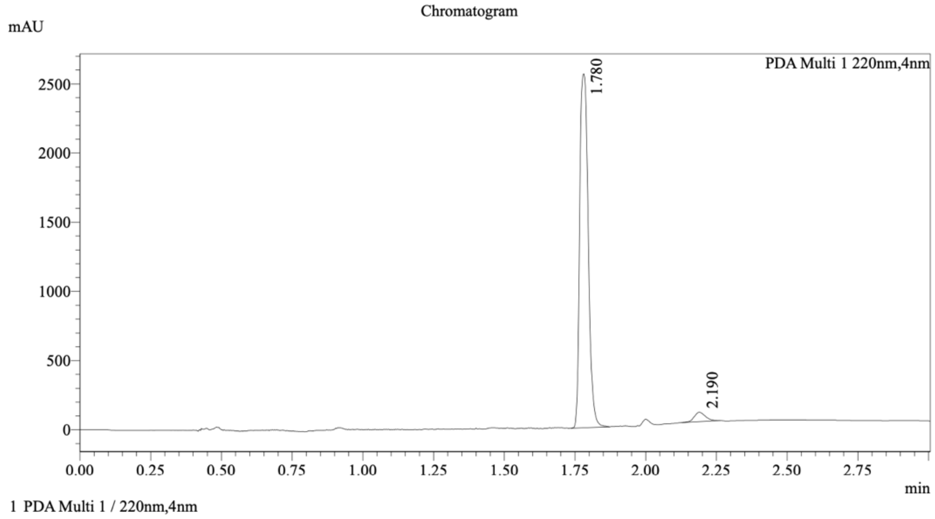

Integration Result

Peak Table

PDA Ch1 220nm

| Peak# | Ret. Time | Height | Height% | USP Width | Area | Area% |
|---|---|---|---|---|---|---|
| 1 | 1.780 | 2547962 | 97.425 | 0.055 | 5360030 | 96.588 |
| 2 | 2.190 | 67356 | 2.575 | 0.075 | 189350 | 3.412 |

**WX-01-12 (14)**

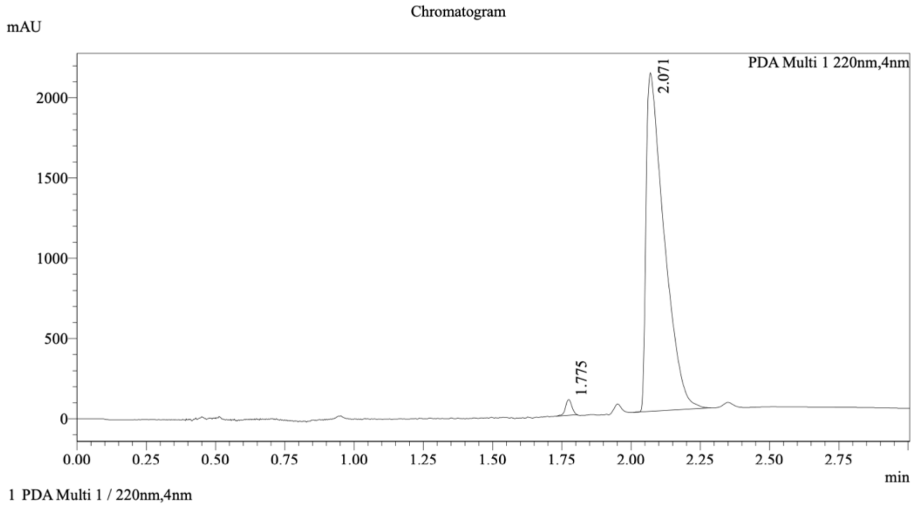

Integration Result

Peak Table

PDA Ch1 220nm

| Peak# | Ret. Time | Height | Height% | USP Width | Area | Area% |
|---|---|---|---|---|---|---|
| 1 | 1.775 | 95306 | 4.343 | 0.047 | 160514 | 1.646 |
| 2 | 2.071 | 2099003 | 95.657 | 0.122 | 9589854 | 98.354 |

**WX-01-10 and WX-01-12 mixture**

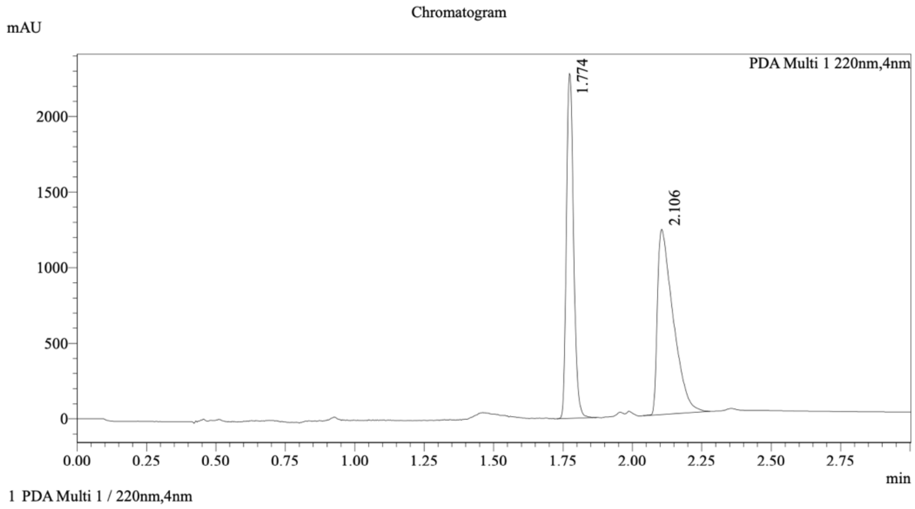

Integration Result

Peak Table

PDA Ch1 220nm

| Peak# | Ret. Time | Height | Height% | USP Width | Area | Area% |
|---|---|---|---|---|---|---|
| 1 | 1.774 | 2254220 | 65.173 | 0.050 | 4107310 | 45.914 |
| 2 | 2.106 | 1204582 | 34.827 | 0.108 | 4838435 | 54.086 |

*Method*

Column: Cellucoat 50×4.6mm I.D., 3 μm

Mobile phase B: EtOH (0.05% DEA);

Gradient elution: 5% to 40% B.

**WX-02-23 (15)**

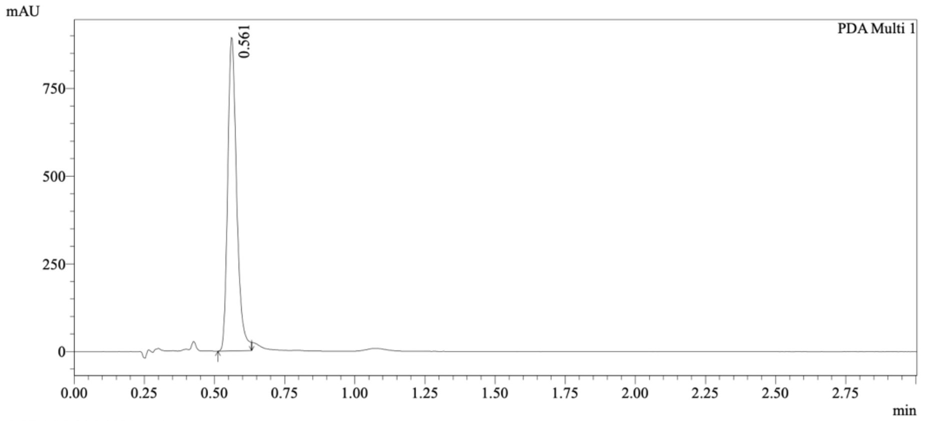

1 PDA Multi 1 / 220nm,4nm

Integration Results

PeakTable

PDA Ch1 220nm

| Peak# | Ret. Time | USP Width | Resolution | Height | Area | Area % |
|---|---|---|---|---|---|---|
| 1 | 0.561 | 0.061 | 0.000 | 852432 | 1989408 | 100.000 |
| Total | | | | 852432 | 1989408 | 100.000 |

**WX-02-43 (16)**

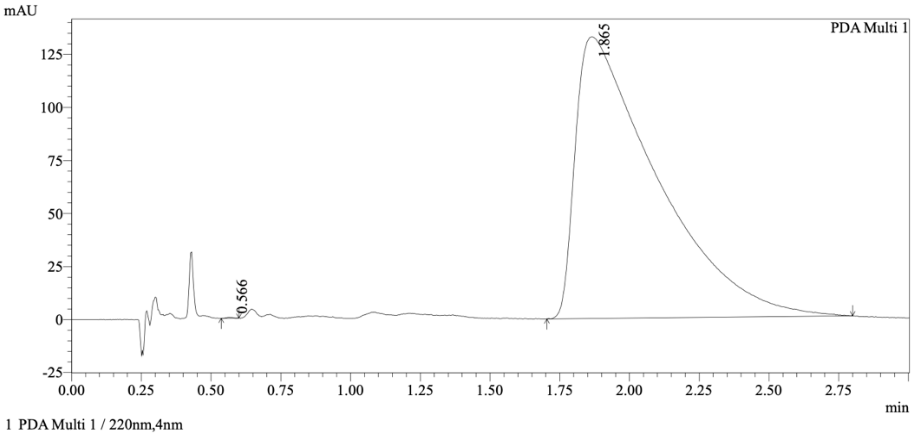

1 PDA Multi 1 / 220nm,4nm

Integration Results

PeakTable

PDA Ch1 220nm

| Peak# | Ret. Time | USP Width | Resolution | Height | Area | Area % |
|---|---|---|---|---|---|---|
| 1 | 0.566 | 0.049 | 0.000 | 390 | 643 | 0.024 |
| 2 | 1.865 | 0.553 | 4.318 | 132541 | 2696012 | 99.976 |
| Total | | | | 132931 | 2696655 | 100.000 |

**WX-02-23 and WX-02-43 mixture**

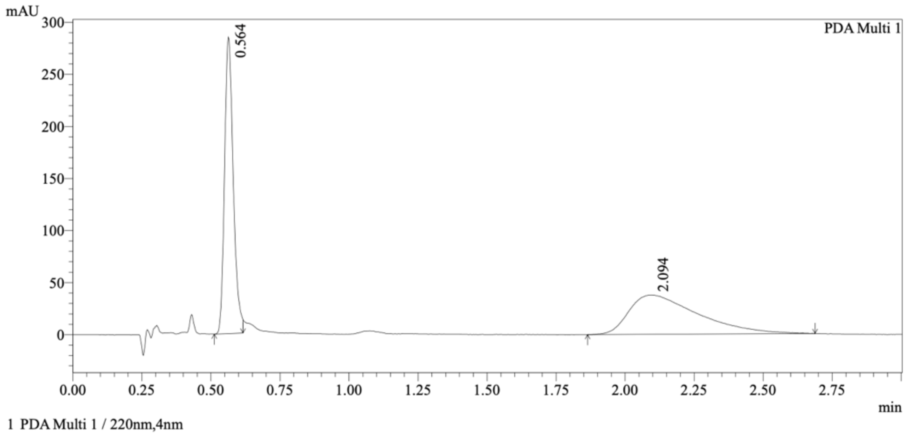

Integration Results

PeakTable

PDA Ch1 220nm

| Peak# | Ret. Time | USP Width | Resolution | Height | Area | Area % |
|---|---|---|---|---|---|---|
| 1 | 0.564 | 0.061 | 0.000 | 278808 | 626639 | 48.682 |
| 2 | 2.094 | 0.453 | 5.954 | 37794 | 660569 | 51.318 |
| Total | | | | 316601 | 1287208 | 100.000 |

*Method*

Column: Chiralcel OD-3 50×4.6mm I.D., 3 μm;

Mobile phase B: MeOH (0.05%DEA);

Gradient elution: 40% B (isocratic)

**EV-96-ctrl (17)**

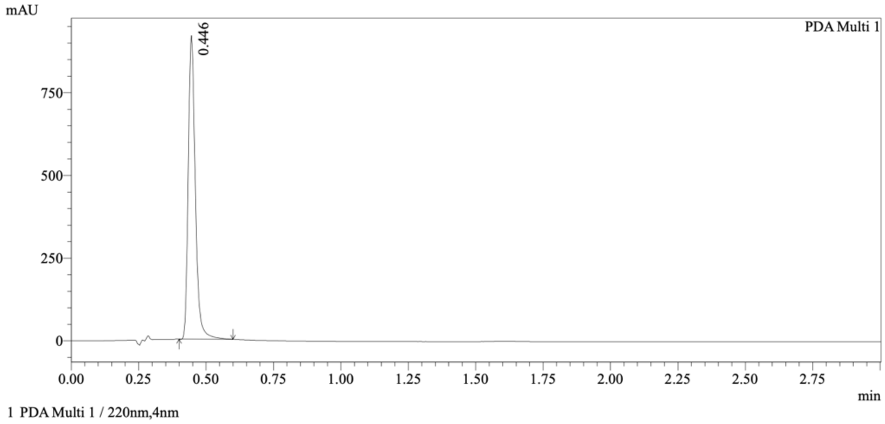

Integration Results

PeakTable

PDA Ch1 220nm

| Peak# | Ret. Time | USP Width | Resolution | Height | Area | Area % |
|---|---|---|---|---|---|---|
| 1 | 0.446 | 0.051 | 0.000 | 901243 | 1679633 | 100.000 |
| Total | | | | 901243 | 1679633 | 100.000 |

**EV-97-ctrl (18)**

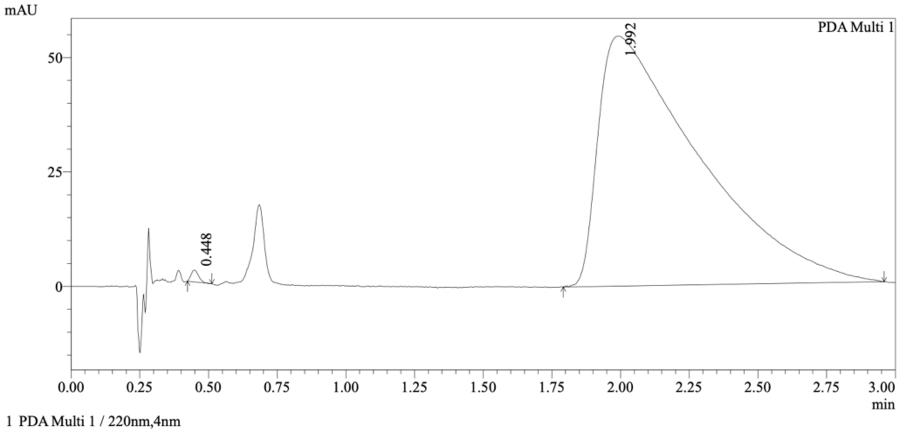

Integration Results

PeakTable

PDA Ch1 220nm

| Peak# | Ret. Time | USP Width | Resolution | Height | Area | Area % |
|---|---|---|---|---|---|---|
| 1 | 0.448 | 0.055 | 0.000 | 2504 | 4904 | 0.349 |
| 2 | 1.992 | 0.710 | 4.040 | 54596 | 1398379 | 99.651 |
| Total | | | | 57100 | 1403283 | 100.000 |

**Mixture of compounds EV-96 and EV-97**

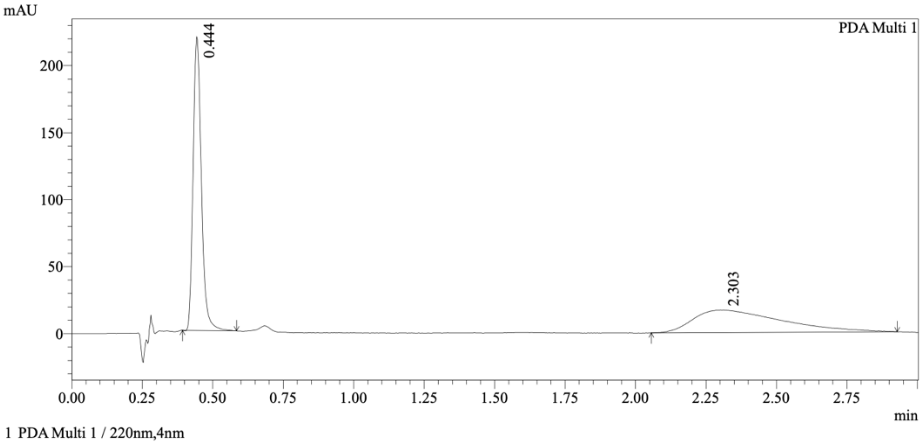

Integration Results

PeakTable

PDA Ch1 220nm

| Peak# | Ret. Time | USP Width | Resolution | Height | Area | Area % |
|---|---|---|---|---|---|---|
| 1 | 0.444 | 0.056 | 0.000 | 214301 | 447926 | 55.168 |
| 2 | 2.303 | 0.569 | 5.944 | 16753 | 364004 | 44.832 |
| Total | | | | 231054 | 811931 | 100.000 |

*Method*

Column: Chiralcel OD-3 50×4.6mm I.D., 3 μm;

Mobile phase B: MeOH (0.05%DEA);

Gradient elution: 40% B (isocratic).

#### EV-98-ctrl (19)

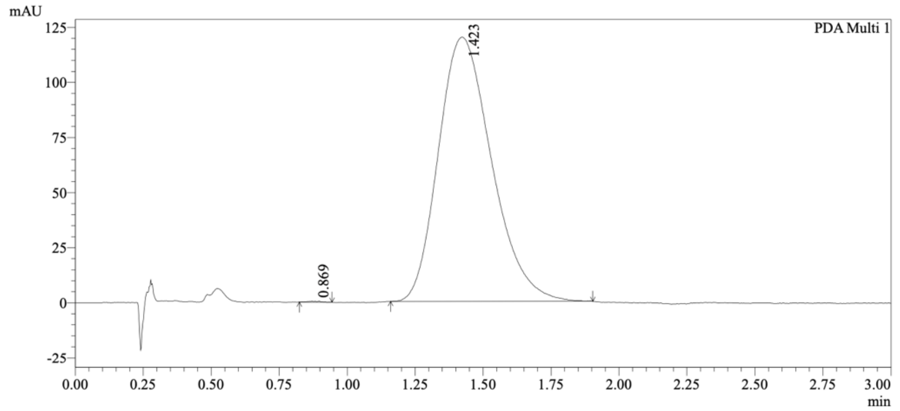

1 PDA Multi 1 / 220nm,4nm

Integration Results

PeakTable

PDA Ch1 220nm

| Peak# | Ret. Time | USP Width | Resolution | Height | Area | Area % |
|---|---|---|---|---|---|---|
| 1 | 0.869 | 0.096 | 0.000 | 348 | 1159 | 0.071 |
| 2 | 1.423 | 0.358 | 2.438 | 119881 | 1634250 | 99.929 |
| Total | | | | 120229 | 1635409 | 100.000 |

#### EV-99-ctrl (20)

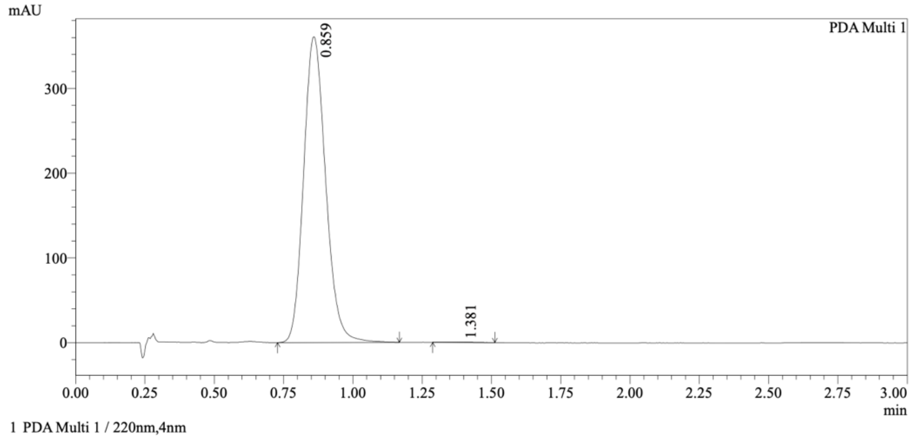

1 PDA Multi 1 / 220nm,4nm

Integration Results

PeakTable

PDA Ch1 220nm

| Peak# | Ret. Time | USP Width | Resolution | Height | Area | Area % |
|---|---|---|---|---|---|---|
| 1 | 0.859 | 0.146 | 0.000 | 359647 | 1997444 | 99.895 |
| 2 | 1.381 | 0.160 | 3.418 | 316 | 2095 | 0.105 |
| Total | | | | 359963 | 1999539 | 100.000 |

**Mixture of compounds EV-98-ctrl and EV-99-ctrl**

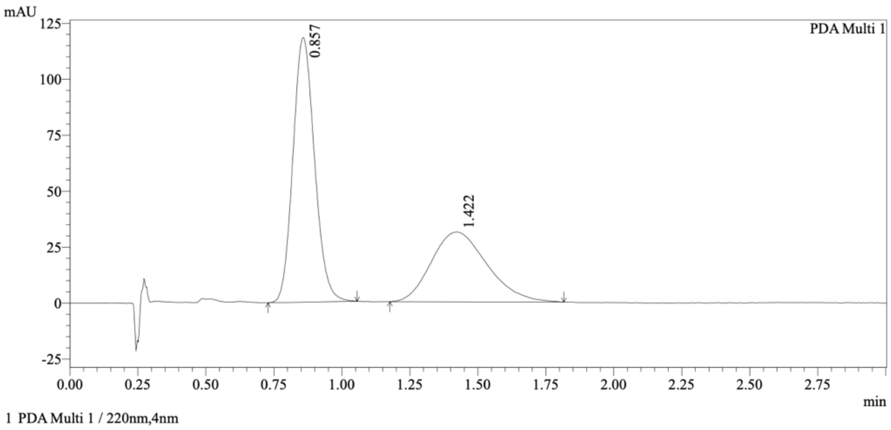

Integration Results

PeakTable

PDA Ch1 220nm

| Peak# | Ret. Time | USP Width | Resolution | Height | Area | Area % |
|---|---|---|---|---|---|---|
| 1 | 0.857 | 0.147 | 0.000 | 117235 | 653341 | 59.496 |
| 2 | 1.422 | 0.376 | 2.161 | 31162 | 444778 | 40.504 |
| Total | | | | 148397 | 1098120 | 100.000 |

*Method*

Column: Chiralpak AD-3 50×4.6mm I.D., 3 μm;

Mobile phase B: MeOH (0.05%DEA);

Gradient elution: 40% B (isocratic).
